# From macro to micro: De novo genomes of Aedes mosquitoes enable comparative genomics among close and distant relatives

**DOI:** 10.1101/2025.01.13.632753

**Authors:** Gen Morinaga, Darío Balcazar, Athanase Badolo, Diana Iyaloo, Luciano Tantely, Theo Mouillaud, Maria Sharakhova, Scott M. Geib, Christophe Paupy, Diego Ayala, Jeffrey R. Powell, Andrea Gloria-Soria, John Soghigian

## Abstract

The yellow fever mosquito (*Aedes aegypti*) is an organism of high medical importance because it is the primary vector for diseases such as yellow fever, Zika, dengue, and chikungunya. Its medical importance has made it a subject of numerous efforts to understand their biology. One such effort, was the development of a high-quality reference genome (AaegL5). However, this reference genome was sourced from a highly inbred laboratory strain with unknown geographic origin. Thus, the reference is not representative of a wild mosquito, let alone one from its native range in sub-Saharan Africa. To better understand the genetic architecture of *Ae. aegypti* and their sister species, we developed two *de novo* chromosome-scale genomes with sequences sourced from single individuals: one of *Ae. aegypti formosus* (Aaf) from Burkina Faso and one of *Ae. mascarensis* (Am) from Mauritius. Both genomes exhibit high contiguity and gene completeness, comparable to AaegL5. While Aaf exhibits high degree of synteny to AaegL5, it also exhibits several large inversions. We further conducted comparative genomic analyses using our genomes and other publicly available culicid reference genomes to find extensive chromosomal rearrangements between major lineages. Overrepresentation analysis of expanded genes in Aaf, AaegL5, and Am revealed that while the overarching category of genes that have expanded are similar, the specific genes that have expanded differ. Our findings elucidate novel insights into chromosome evolution at both microevolutionary and macroevolutionary scales. The genomic resources we present are additions to the arsenal of biologists in understanding mosquito biology and genome evolution.

**Significance:** *Aedes aegypti* is a major arboviral disease vector found throughout the tropics and sub-tropics. Its subspecies differ ecologically, as native sub-Saharan African form feeds on mammals generally and inhabit both sylvatic and domestic areas and the global invasive form preferentially feeds on humans and lives primarily domestic areas. Their medical importance has prompted the development of a high-quality reference genome, but it was sourced from an inbred laboratory strain of unknown origin. Here, we leveraged PacBio HiFi sequencing and HiC sequencing to develop the first de novo genome of *Ae. aegypti* sampled its native range in Burkina Faso. We also present a de novo genome of *Ae. mascarensis*, its sister species. Our genomes are comparably contiguous and complete to the reference genome. Comparative genomic analysis using our genomes and other culicid reference genomes reveal extensive chromosomal rearrangements.

## Introduction

Mosquitoes (family Culicidae) are major vectors of arboviruses and parasites that cause minor to severe illnesses that affect millions of people globally (Yee et al. 2022). To mitigate the impact of mosquitoes on global public health, tremendous efforts have been made to elucidate mosquito biology. One such effort is the development of numerous genomic resources of mosquito species, especially those considered medically important such as *Anopheles* and *Aedes* mosquitoes. An example of these resources is the reference genome for *Aedes aegypti* (henceforth, ‘AaegL5’; Matthews et al. 2018), the primary vector of arboviruses, including yellow fever, dengue, chikungunya, and Zika viruses (Pierson & Diamond 2020).

At the time of its development, AaegL5 utilized available technologies to achieve an accurate, complete, and contiguous chromosome-level genome assembly. Beyond improving upon the contiguity of previous versions of the reference (Dudchenko et al. 2017), AaegL5 also provided an improved set of gene annotations (Matthews et al. 2018), which allows for finer-scale mapping of genes and gene families. Indeed, the development of this reference has been invaluable—allowing researchers to study in detail *Ae. aegypti* transcriptomics (Herre et al. 2022), developmental biology (Herre et al. 2022), population genetics (Schmidt et al. 2020; Soghigian et al. 2020; Gloria-Soria et al. 2022), species distribution modeling (Rose et al. 2020), and phylogenomics (Soghigian et al. 2023). This resource is also valuable for elucidating biological differences between the subspecies *Ae. aegypti formosus* and *Ae. aegypti aegypti* (Aaf and Aaa, respectively) which differ in their bionomics—Aaf is a native to sub-Saharan Africa, inhabits both forested and domestic areas, and take blood meals from a variety of mammals; Aaa is a found globally outside of Africa, inhabits urban areas, and females have a strong affinity to feed on humans (Powell et al. 2018).

Despite its value as a genomic resource, AaegL5 is not without its shortcomings. The sequencing technology available at the time precluded the authors from using wild mosquitoes for two, co-related reasons. First, library construction requires high DNA input—a major hurdle for small insects such as flies and mosquitoes where a single individual may not be sufficient to extract enough source DNA, leading to assemblies where multiple individuals are pooled.

Second, the long-read sequencing platforms available at the time were error-prone and required polishing using other, more accurate sequencing technologies (i.e., Illumina short-reads), which again would require more input DNA from additional individuals. Matthews et al., (2018) mitigated both challenges by sequencing 80 male siblings from a highly inbred laboratory strain (Liverpool), thereby reducing mis-assemblies resulting from high levels of heterozygosity (Whibley et al. 2021) whilst also extracting enough input DNA to meet library construction requirements.

A corollary of using an inbred laboratory colony for a reference genome, of course, is the reduction of heterozygosity and genetic diversity that it represents. This is not problematic *per se*—a single individual is a mere snapshot of the complex ebbs and flows of population dynamics within a single population. However, when laboratory colonies have measurably diverged from wild or source populations, they may no longer be representative of wild populations, leading to erroneous conclusions and problems with reproducibility (Brekke et al. 2018). Recent work (Gloria-Soria et al. 2019) has shown that laboratory strains of *Ae. aegypti* exhibit significantly lower genetic diversity than their wild counterparts. Furthermore, some strains do not cluster with any wild mosquito populations (CDC, Liverpool, and Orlando) or have diverged and/or become contaminated as strains were passed among laboratories (Liverpool and Rockefeller; Gloria-Soria et al. 2019). Lastly, Gloria-Soria et al., (2019) showed that the Liverpool strain, originally thought to have been sourced from West Africa (Macdonald 1962), does not cluster with any Aaf populations.

Within Culicidae, chromosome-scale genome assemblies are taxonomically concentrated in the Anophelinae, and these assemblies have allowed biologists to uncover some of the intricate evolutionary dynamics that bridge the divide between microevolution and macroevolution (Neafsey et al. 2015; Lukyanchikova et al. 2022). Recent efforts to expand the phylogenetic sample of mosquitos have produced numerous chromosome-scale assemblies in the Culicinae (e.g., Peng et al. 2021; Ryazansky et al. 2024). This effort to expand the phylogenetic scope beyond Anophelinae allows biologists to further investigate the structural variation between members of major clades (e.g., subgenera, tribes).

Here, we present the first *de novo* genome assembly of a wild *Ae. aegypti formosus* (Aaf) specimen from West Africa and a *de novo* assembly of *Ae. mascarensis* (Am)*—*a partially reproductively isolated sister species of *Ae. aegypti* found on the island of Mauritius in the south west Indian Ocean (Hartberg & Craig Jr 1970). *Aedes mascarensis* diverged from Aaf roughly 8–10 MY (Soghigian et al. 2020). These two species are members of what is now known as the Aegypti group (Le Goff et al. 2013; Soghigian et al. 2020). Each assembly was built from a single wild-caught mosquito from their respective locales (Aaf: Burkina Faso; *Ae. mascarensis*: Mauritius) using the recently developed Pacific Biosciences high fidelity (PacBio HiFi) sequencing platform (Wenger et al. 2019). The high accuracy (99.999%) of PacBio HiFi reads allows us to sequence a relatively highly heterozygous individual while the length of the reads (>13 kbp) helps to span and resolve highly repetitive regions ubiquitous in the *Ae. aegypti* genome (Matthews et al. 2018). We also compare new genomes with those of other mosquitoes and place them in an evolutionary context to understand how genes and genomic structure have changed across major culicid lineages. Lastly, we present a method for comparing draft assemblies and implement it as a new, publicly available, *asmidx* package for R.

## Methods

### Sample collection

We sampled for wild mosquitoes in two locations: Ouagadougou, Burkina Faso, for *Aedes aegypti formosus* (Oct. 2021); and Chamarel, Mauritius for *Ae. mascarensis* (May 2022). In both locations, we collected mosquito eggs by placing ovitraps lined-up with seed-germination papr. These eggs were shipped to The Connecticut Agricultural Experiment Station (CAES) for rearing. We reared mosquitoes from eggs at CAES by submerging the paper containing the eggs in deionized water and provided TetraMin® Tropical Flakes as *ad libidum* as food source for larvae. Once mosquitoes pupated, we transferred the pupae into medicine cups filled with larval water and placed them into insect rearing cages (12 x 12 x 12 inches/28 liters) where the adults emerged. Larval trays and cages were maintained in an incubator at 27°C with a 12:8 light/dark cycle throughout the rearing process. We provided adults with *ad libitum* sugar water for 3–5 days until they were collected in dry ice for the DNA extraction protocol.

### DNA sequencing for genome assembly and Hi-C genome-wide DNA cross-linking

To generate DNA sequences for PacBio HiFi for both Aaf and Am, we collected and froze adult female mosquitoes on dry ice, then homogenized individuals with a sterile DNAse/RNAse free plastic pestle. We eluted the homogenate using 180 µl of PBS and processed the solution using the MagAttract HMW DNA Kit (Qiagen, Germantown, MD, USA) following the frozen tissue protocol from the manufacturer. We then sent the extracted DNA to the Maryland Institute of Genome Sciences of the University of Maryland for low input library preparation, where two Sequel II 8M SMRT Cell runs (CCS/HiFi mode - 30 hour movie) were used to obtain sequences.

For HiC sequencing, we pooled multiple individuals (four females for *Ae. aegypti formosus* and eight females for *Ae. mascarensis*) together and pulverized them in dry ice. We then cross-linked samples and prepared *Ae. aegypti formosus* library with the Arima High Coverage HiC kit and Arima HiC+ kit (Arima Genomics, San Diego, CA, USA), following manufacturer protocols. For *Ae. mascarensis*, we prepared HiC libraries using the xGen ssDNA & Low Input DNA Library Prep Kit (IDT, San Diego, CA, USA). We then sequenced both HiC libraries at the Yale Center for Genome Analysis to achieve 100 million 150 bp paired-end reads.

### Genome assembly

The PacBio HiFi sequencing platform and the programs (or the specific modes) built to handle these data are still nascent, so we used four different assemblers and compared the outputted draft assembly from each program. We used *HiCanu* (Nurk et al. 2020), *flye* (Kolmogorov et al. 2019), *hifiasm* (Cheng et al. 2021), and *IPA* (available at: https://github.com/PacificBiosciences/pbipa). For all programs, we specified an estimated haploid genome size of 1.3Gbp and used default settings and set flags necessary to assemble PacBio HiFi reads (Supplementary Table S1).

We also compared the performance of two purging programs designed to identify and remove duplicate haplotypes from the draft assemblies—*purge_haplotigs* (Roach et al. 2018) and *purge_dups* (Guan et al. 2020) (henceforth, *ph* and *pd* respectively). Note that *hifiasm* and *IPA* employ a purging step as part of their respective assembly pipelines by default—*hifiasm* employs a variant of *pd* with a different algorithm for haplotype identification, and *IPA* simply uses *pd*. Thus, in our workflow, assemblies output by these programs were purged twice, which allowed us to assess “out-of-the-box” performance of all programs.

To assemble the mitochondrial genomes, we used the program *mitohifi* (Uliano-Silva et al. 2023) in ‘reads’ mode and input the PacBio HiFi reads. This program was reference guided, so for both species we used the *Ae*. *aegypti* complete mitochondrion found on GenBank (OR544945.1).

### *asmidx*: A holistic approach to assessing genome quality based on user input

To quantify assembly metrics, we fed each intermediate assembly to the program *Inspector* ver. 1.0.1 (Chen et al. 2021). This program quantified basic assembly metrics, detects assembly errors at the structural (expansion, collapses, inversions, and haplotype switches ≥50Bp) and small scales (base substitutions, expansions, collapses <50Bp), and attempts to correct them. We also assessed gene content completeness for each assembly generated using *BUSCO* ver. 5.2.2 and the Diptera OrthoDB data set ver. 10 (n = 3,285 single-copy orthologs).

Common genomic metrics (e.g., N50) and gene content of an assembly can be effective indicators of assembly quality, but there is no consensus on which of these characteristics (or what set of them) best characterizes a “good” genome assembly. Furthermore, it is unknown whether each assembly program used in tandem with a purging program outputs similar quality genome assemblies. To facilitate identifying the “best” assembly derived from the same set of reads, we wrote an application using the *shiny* package ver. 1.7.4 for the R statistical programming language ver. 4.2.1, which we call *asmidx* (available at https://github.com/genmor/asmidx). The application takes as input a user-generated data set in tabular format, with column headers where each row contains an assembly, and each column contains a metric describing that assembly. The user can then select assembly metrics that should be maximized and minimized to assess quality. Additionally, the user can identify a genome size column and input a known genome size (e.g., from a reference genome) which will be used to compute relative genome size differences, where smaller differences in relative genome size will be considered better. Each chosen metric is then feature-normalized to be between 0 (the worst) and 1 (the best). A row sum is then computed and multiplied by 100 resulting in normalized scores which are used to rank each assembly by quality. We also allow users to differentially weight each of the selected characters. To make this process intuitive, we allow for any positive value for weighting. The weights are multiplied to their respective feature-normalized columns, row sums re-computed, and multiplied by 100 to output a weighted score. For both weighted and unweighted scores, higher values will be associated with better assemblies based on the quality metrics supplied (and weighted) by the user. The application outputs these rankings in tabular format and visually represents them using a lollipop plot, both of which the user can download. In addition to this shiny application, we wrote helper functions to convert output from Inspector into tabular format.

To rank our assemblies, we considered four sets of metrics: gene content (duplicated, fragmented, and missing BUSCOs), structural errors in the assembly (total number of expanded bases, collapsed bases, and inverted bases), contiguity (N50), and genome size difference relative to the reference. Our rankings maximized N50 and minimized all other metrics. We weighted all structural errors and N50 by 0.1, duplicated and missing BUSCOs by 0.15, fragmented BUSCOs by 0.125, and relative genome size difference by 0.175. For each specimen, we used the assembly with the highest weighted score for scaffolding using Hi-C.

### Scaffolding contigs to chromosomes via Hi-C and post processing

To prepare cross-linked, paired-end Illumina short-reads for use in scaffolding, we trimmed the first five bases from the reads using the program *fastp* (ver. 0.23.4; Chen et al. 2018). We then followed the Arima Genomics mapping pipeline (available at https://github.com/ArimaGenomics/mapping_pipeline). This pipeline relies on *BWA* (ver. 0.7.17-r1198-dirty; Li and Durbin 2009), *samtools* (ver. 1.15; Danecek et al. 2021), *picard* (ver. 2.2.4; Broad Institute 2019), and custom Perl scripts for aligning the short-reads to a draft contig-level genome assembly and preparing it for scaffolding (Table Program details). To scaffold, we used the program *Yahs* (ver. 1.2a.2; Zhou et al. 2023). We manually curated the outputted scaffold-level assembly using the programs *Juicer tools* and *Juicebox* (ver. 1.11.08; Durand et al. 2016) to further remove duplicate contigs and correct mis-assemblies (i.e., inverted and mis-joined contigs and scaffolds) and generated finalized contact maps for visualization purposes using *HapHiC* (ver. 1.0.5; Zeng et al. 2024). After manually curating the scaffolds, we used *TGS-gapcloser* (ver. 1.2.1; Xu et al. 2020) to close gaps in the assembly. We inspected gene content of this assembly using *BUSCO* and then finalized the gap-filled assemblies by using a custom script to remove “debris” sequences (i.e., contigs and scaffolds with duplicate HiC signal) and scaffolds/contigs containing only duplicate BUSCOs that were not located on the chromosomal scaffolds. We concatenated these “debris” sequences together with the duplicates detected by the purging program. We then fed these assemblies to the *BlobToolKit* (ver. 4.2.1; Challis et al. 2020) suite to determine whether our assemblies contained sequences from contaminants or endosymbionts and to output final summary statistics.

### Validating structural variation and departures from AaegL5

To ascertain the validity of the structural variations we observed in Aaf and Am (relative to AaegL5), we used NCBI BLAST (ver. 2.12.0+; Camacho et al. 2009) to create databases from our scaffolded *Aaf* and *Am* assemblies to find positional hits of 88 *Ae. aegypti* bacterial artificial chromosome (BAC) clone sequences (Matthews et al. 2018; Supplementary table S2). We retained only the best hits (i.e., highest bit score) for each BAC clone and visualized their positional order in both assemblies using the R package *ChromoMap* (ver. 4.1.1; Anand and Rodriguez Lopez 2022; Supplementary figure S1). After this validation step, we aligned the scaffolded *Aaf* assembly to AaegL5 using *minimap2* (Li 2018), and the resultant alignment file fed into *SyRI* (ver. 1.6.3; Goel et al. 2019) which identified nucleotide synteny and structural variation (i.e., duplications, translocations, and inversions). We performed the same analysis for Am but chose not to interpret nucleotide synteny because it is too divergent from *Ae. aegypti*, even after using less stringent *minimap2* settings (i.e., -asm20) and found the resulting output uninterpretable (see Supplementary figure S2).

### Genome structural annotation

We used the *RepeatModeler* pipeline (ver. 2.0.4; Flynn et al. 2020) to model and identify repetitive elements in the genomes. After generating a *de novo* repeat database from our draft assemblies, we soft-masked the assemblies using *RepeatMasker* (Smit et al. 2013) in four iterations, passing each outputted soft-masked fasta files to the subsequent step: 1) mask only simple repeats; 2) mask repeats using the -species flag with ‘diptera’ which queries the Dfam database (ver. 3.7; Storer et al. 2021) for dipteran repeat sequences; 3) mask repeats identified in *Ae. aegypti* (Nene et al. 2007) from TEfam repeat database; 4) mask repeats based on the *de novo* repeat database created from *RepeatModeler*. We then quantified the diversity and divergence relative to the consensus sequences of the repetitive content in Aaf and Am using the ‘*calcDivergenceFromAlign.pl’* script included with *RepeatMasker.* This script estimated divergence using the Kimura (K81) model of sequence evolution modified to account for the high mutability of “CG” sites (Tsunoyama et al. 2001).

We input the final, soft-masked assemblies into the *BRAKER2* genome annotation pipeline (ver. 3.0.3; Stanke et al. 2006b, 2008; Hoff et al. 2016, 2019; Brůna et al. 2021) with the Arthropoda protein data set obtained from *OrthoDB* (ver. 11; Kuznetsov et al. 2023). *Braker* uses *GeneMark-ES* (Lomsadze et al. 2005) and *ProtHint* (Brůna et al. 2020) to predict protein coding genes in the assembly, then aligns these predicted proteins and regions using *DIAMOND* (ver. 0.9.24.125; Buchfink et al. 2015) and *SPALN* (Iwata & Gotoh 2012). High-confidence hits output by these programs are then fed into *GeneMark-EP+* (Brůna et al. 2020) and *Augustus* (ver. 3.4.0; Stanke et al. 2006a) to output gene predictions. Following recommendations from the program authors, we used the final output from *Augustus*. We then used a Python script ‘*selectSupportedSubsets.py’* included with *Braker* to exclude genes predicted without any external support (i.e., no support from OrthoDB). We used this output for all analysis that involved the proteome. We assessed the quality of these annotations using *BUSCO* in protein mode and *AGAT* (ver. 1.2.0; Dainat et al. 2023) to quantify annotation metrics after we subset the output from *BRAKER2* to exclude genes supported only through computational predictions and keeping only the longest isoforms.

### Comparative genomic analysis

We compared the Aaf and Am assemblies with other Culicidae, including AaegL5, and nine (eight culicid; one outgroup) other annotated, high-quality, chromosome-level reference assemblies available on NCBI that represent the breadth of phylogenetic diversity of the family. These assemblies were: *Anopheles cruzii* (subgenus *Kerteszia*)*, An. darlingi* (subgenus *Nyssorhynchus*)*, An. gambiae* (subgenus *Cellia*)*, An. ziemanni* (subgenus *Anopheles*)*, Armigeres subalbatus* (tribe Aedini)*, Sabethes cyaneus* (tribe Sabethini)*, Culex pipiens pallens* (tribe Culicini)*, Cx. quinquefasciatus* (tribe Culicini), and the sandfly, *Phlebotomous papatasi* (Supplementary table S3). We assessed gene order synteny using the R package *GENESPACE* (ver. 1.2.3; Lovell et al. 2022). *GENESPACE* uses *Orthofinder* (Emms & Kelly 2019) to identify orthologous genes across a set of species and assesses synteny and collinearity of the orthologs between all pairwise combinations of species using *MCScanX* (Wang et al. 2012). We set AaegL5 as the reference for the riparian plot output by *GENESPACE*.

We also assessed the evolution of gene families across these genomes using part of the *compare_genomes* pipeline (Paril et al. 2023). This pipeline chained several programs to: 1) identify orthologous genes (*Orthofinder*); 2) infer a dated phylogeny using the orthologs (*IQ-Tree2*; Minh et al. 2020); 3) assess gene family expansion and contraction (*Cafe5*; Mendes et al. 2021); and 4) use the PANTHER classification system (Mi et al. 2013) to assign gene family and function. For phylogenetic inference, the pipeline set IQ-Tree2 to use a multi-partition model (Chernomor et al. 2016) and performed model selection using ModelFinder (Kalyaanamoorthy et al. 2017). It additionally used ultrafast bootstrap approximation (Hoang et al. 2018) to estimate branch support and a least squares algorithm to date the inferred phylogeny (To et al. 2016). To analyze evolutionary gene expansion and contraction, we modeled genes families to evolve at different rate categories through a γ-parameter with K = 4 categories in *Cafe5*. For gene function and ontology, we used the biological process set (GO:0008150) from *An. gambiae* (taxon ID: 7165), and the pipeline classified gene function using PantherHMM 17 (Mi et al. 2013, 2019). Although *compare_genomes* performs GO enrichment and overrepresentation as part of the pipeline, we elected to use the intermediate output to perform our own. We did this by querying the PANTHER DB web tool (www.pantherdb.org; ver. 18.0; accessed: Apr. 22, 2024) with a list of significantly expanded orthogroups (from *Cafe5*) for AaegL5, Aaf, and Am (separately and together) to perform a statistical overrepresentation test using Fisher’s exact test using the ‘GO biological process complete’ annotation set of *An. gambiae* and corrected for multiple testing by accounting for false discovery rate (Benjamini & Hochberg 1995). We downloaded the full data table of results for each taxon and filtered each list in R to include only those where P_FDR_ < 0.01 for a given taxon. We then examined the semantic similarity of these overrepresented GO terms based on the method of Wang et al. (2007), used K-means clustering to determine similar sets of terms, then corrected the P-values a final time for multiple comparisons (again accounting for false discovery rate) using the R package *simplifyEnrichment* (ver. 1.12; Gu and Hübschmann 2023). For a more holistic analysis of the Aegypti group, we repeated the above overrepresentation test on PANTHER DB, this time included all three expanded sets and outputting only significantly overrepresented GO terms (P_FDR_ < 0.01). Lastly, we took advantage of the fact that we identified gene families and their positions in the Aaf genome, so we mapped the ones specifically located in putatively inverted regions (relative to AaegL5). We matched gene identity and names by creating a local blast database from AaegL5 and queried the orthologs in the inverted regions, limiting output to a single alignment with e-values less than 1e-60.

## Results

### Identifying the best combination of HiFi assembler and purging program

We generated twelve draft assemblies each for Aaf and Am—four outputs directly from the assemblers and four of each output from purging programs *ph* and *pd* after taking the outputs from the assemblers as input and removing duplicated contigs and haplotigs. We provide detailed comparisons in the supplement (Supplemental results; Supplementary table S4). In brief, *asmidx* allowed us to identify *HiCanu* paired with *ph* output the best Aaf assembly, and *hifiasm* paired with *ph* output the best Am assembly (Fig. 1). While not free of arbitrary decisions, *asmidx* allows users to compare draft assemblies transparently and flexibly.

**Fig. 1.**
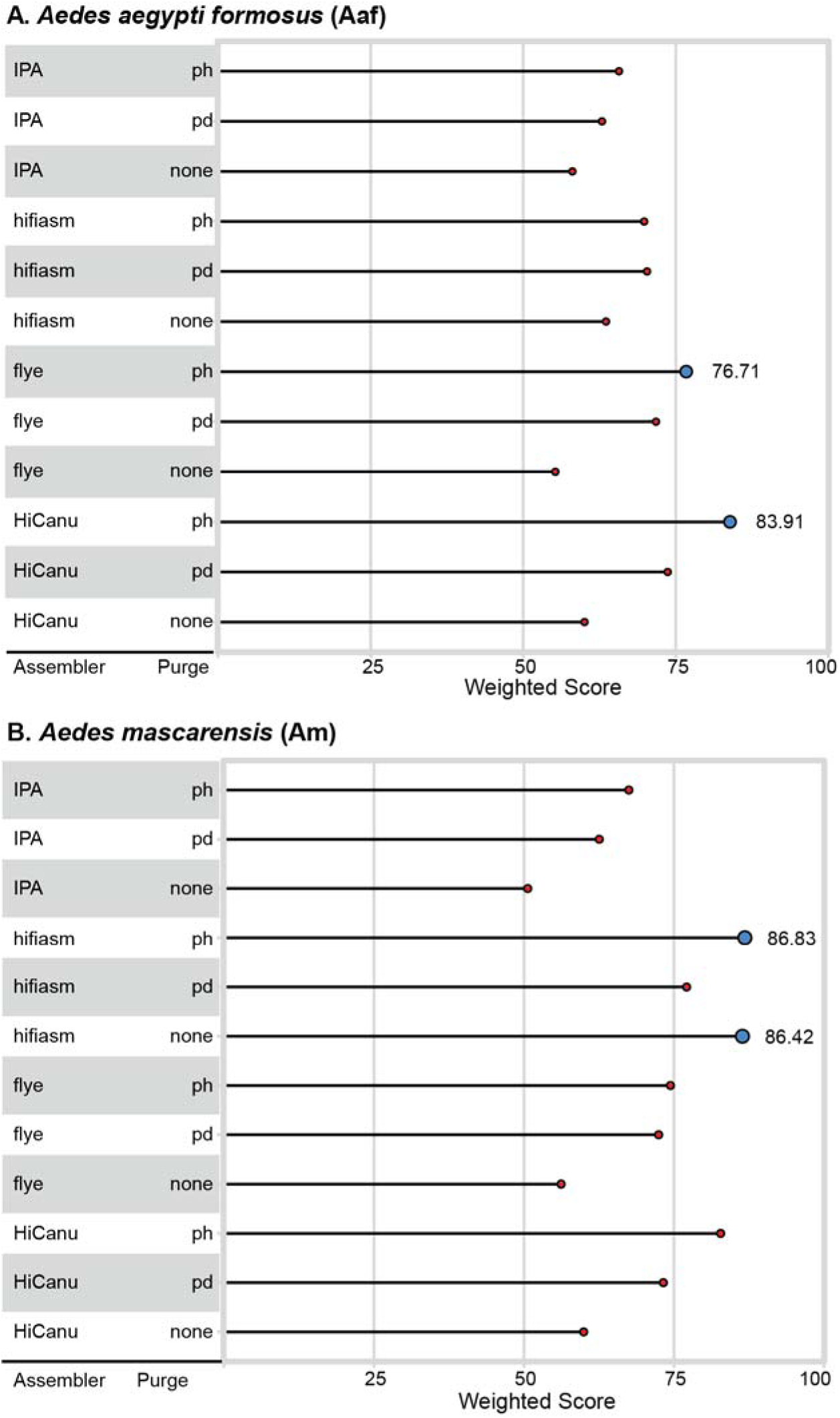
Lollipop plot ranking each draft assemblies for (A) *Aedes aegypti formosus* (Aaf) and (B) *Ae. mascarensis* (Am). The two best assemblies for each taxon is indicated by larger, blue circles. The scores based on duplicated (0.15), fragmented (0.125), missing (0.15), collapses (0.10), expansions (0.1), inversions (0.1), N50 (0.1), and relative genome size (0.175) [metric (weight)].

### Final assembly characteristics

Scaffolding the assemblies using Hi-C, and manually curating the scaffolds using *Juicebox* (Durand et al. 2016) substantially improved the contiguity of the assemblies. For Aaf, N50 saw an 82-fold improvement and reduced L50 from 82 to 2, yielding an assembly with 706 scaffolds, 1,124 contigs, and total assembly size of 1.24 Gbp (Supplementary figures S3 and S4; Table 1). In terms of gene content, we detected 3,127 (95.2%) complete single-copy, 55 (1.7%) duplicated, 45 (1.4%) fragmented, and 58 (1.7%) missing BUSCOs from Diptera orthodb10 ortholog set (n = 3,285). For Am, scaffolding improved N50 by 30-fold and reduced L50 from 22 to 2, outputting an assembly with 74 scaffolds, 269 contigs, and a total assembly size of 1.29 Gbp (Supplementary figures S3 and S4; Table 1). We detected 2,990 (91%) single-copy, 179 (5.4%) duplicated, 48 (1.5%) fragmented, and 68 (2.1%) missing BUSCOs. We attempted to fill gaps in both assemblies using the HiFi reads, but neither resulted in dramatic reduction in scaffold count, although we were able to fill some gaps between contigs in both assemblies (Supplementary figure S3; Table 1). We additionally assembled the mitochondrial genome of these individuals, which yielded mitogenomes at roughly 16 kbp total size for both species.

**Table 1.** Final assembly metrics for Aaf (*Aedes aegypti formosus*) and Am (*Ae. mascarensis*) compared to those of AaegL5 (*Ae. aegypti*).

| Metric | AaegL5 | Aaf | Am |
| --- | --- | --- | --- |
| # Scaffolds | 2,310 | 706 | 74 |
| # Contigs | 2,539 | 1,124 | 269 |
| Total length (Gbp) | 1.279 | 1.239 | 1.289 |
| Gap % | 0.002 | 0.007 | 0.003 |
| L50 | 2 | 2 | 2 |
| N50 (Mbp) | 40.978 | 39.859 | 42.609 |
| GC ± SD | 0.382 ± 0.029 | 0.381 ± 0.074 | 0.381 ± 0.04 |
| Mitogenome length (Mbp) | 16,790 | 16,617 | 16,428 |
| # Exons | 94,104 | 63,767 | 62,375 |
| # Genes | 19,203 | 17,672 | 17,009 |

Structural annotation of both assemblies showed their compositions were proportionally similar to one another (Fig. 2). Repetitive elements comprise the majority of genomic content (up to nearly 80%)—approximately 20% of the genomic content are classified as LTR retrotransposons and roughly 15% attributed to non-LTR retrotransposons (Fig. 2). In both assemblies, DNA transposons accounted for 6% of the genomic content, and approximately 1% of the assemblies were classified as helitrons (Fig. 2). Additionally, approximately 30% of both assemblies were considered repetitive, but unable to be classified (Fig. 2). For both assemblies, genomic repeat contents appear to have accumulated recently relative to the consensus repeat sequences with the peak occurring at 5% and 3% (Aaf and Am, respectively; Fig. 2). Unmasked, genomic content for both assemblies accounted for 22% (274 Mbp) and 21% (276 Mbp) of their genomes (Aaf and Am, respectively; Fig. 2). Exonic content accounted for 28 and 27 Mbp for Aaf and Am respectively (roughly 2% for both genomes) and intronic content accounting for 15% (211 Mbp) and 16% (230 Mbp) for Aaf and Am respectively. Lastly, monoexonic genes constituted 17% (3,120/17,009) and 16% (2,764/17,672) of the gene contents of Aaf and Am respectively.

**Fig. 2.**
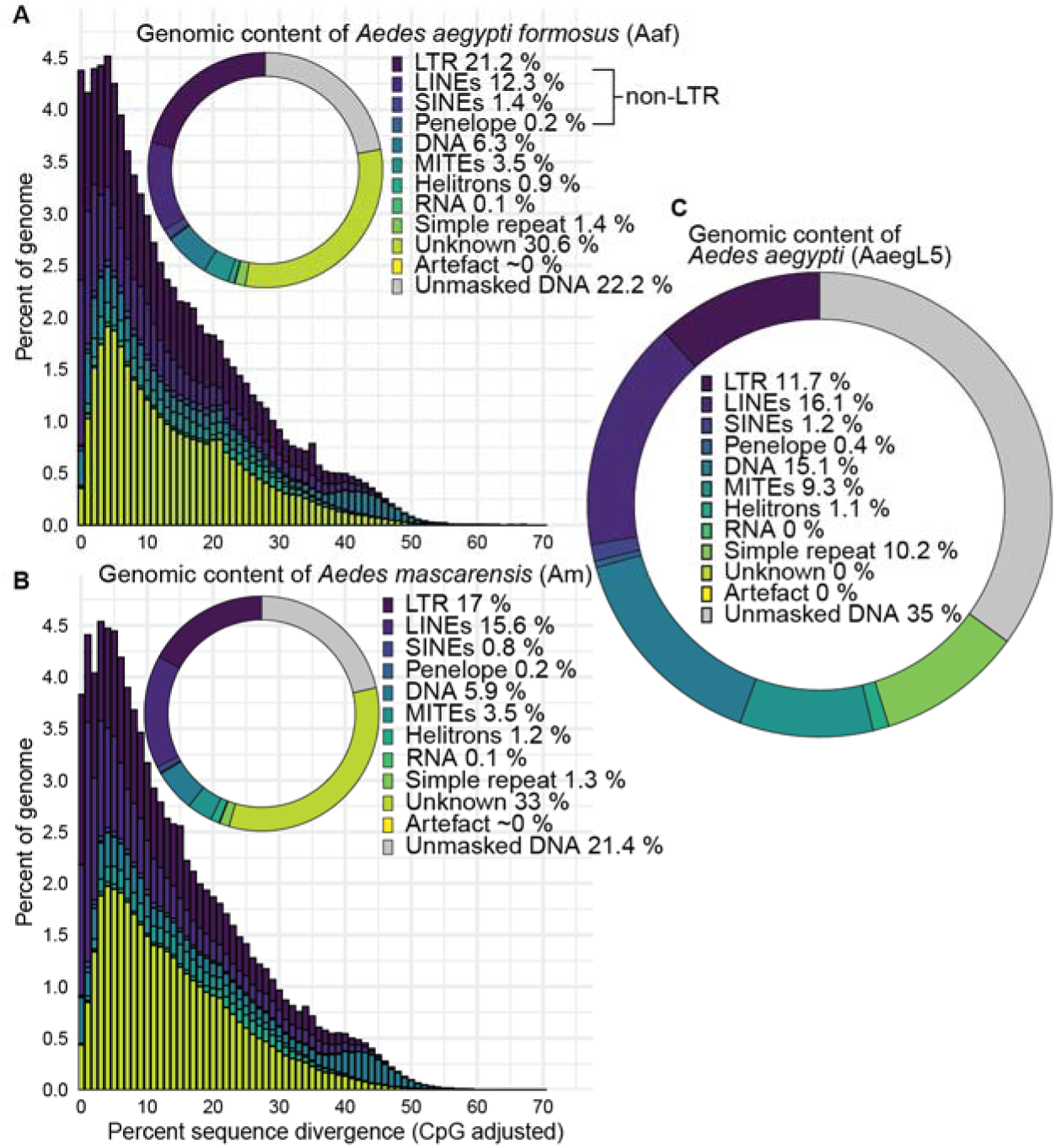
Genomic content for *Aedes aegypti formosus* (Aaf; A), *Ae. mascarensis* (Am; B) *Aedes aegypti* reference genome found by Matthews et al., (2018) [AaegL5; C] and repeat landscape plots (A and B only). In the landscape plots, sequence divergence is shown in 1% intervals. Sequence divergence of the landscape plot was estimated using the Kimura model of sequence evolution modified to account for the high mutability of CpG sites. Landscape plots do not account for “Unmasked DNA”. Categories with trivial number of bases are shown as “∼0%” (A and B), while “0%” for C are actual 0s.

### Identifying putative endosymbiont and contaminant sequences

We assessed both the gap-filled assemblies and the alternate assemblies (e.g., assemblies consisting of haplotigs, duplicates, and ‘debris’) for any potential endosymbionts using the Blobtoolkit pipeline (Challis et al. 2020). Of the sequences in the primary Aaf assembly, we traced 1.18 Gbp to Arthropoda, while the rest (54.4 Mbp) yielded no hits. In the alternate assembly, we detected two sequences, a contig and a scaffold, totaling 1.59 Mbp originating from α-proteobacteria (Supplementary figure S5). We isolated and removed this sequence from the alternate Aaf assembly and queried the first 4400 bp of the scaffold sequence on BLAST. The highest hit (99.89% identity) was for a *Rickettsia* endosymbiont found in *Cimex lectularius* (GenBank Accession #: CP084572.1), while the next two highest hits (88.73%, 88.32%), were similarly for *Rickettsia* endosymbionts isolated from *Oedothorax gibbosus* and *Culicoides impuctatus* respectively (GenBank Accession #:OW370493.1; CP084573.1). In the primary Am assembly, we traced 1.28 Gbp of the sequences to Arthropoda and the rest (7.74 Mbp) yielded no hits. Unlike the Aaf alternate assembly, the Am alternate assembly did not contain any sequences from that did not originate from either Arthropoda or yielded no hits.

### Assessing structural rearrangement between AaegL5 and Aaf

We found a substantial degree of synteny between AaegL5 and Aaf, amounting to 1.08 Gbp or 87% of the total Aaf assembly exhibiting synteny with AaegL5 (Fig. 3). The Aaf assembly also exhibited 175 inversions (most of which are small), totaling 44 Mbp or 3.5% of the total assembly length. We found 870 translocations totaling 4.2 Mbp in length (0.34% of total assembly length). The Aaf assembly also exhibited 490 instances of duplicated sequences which totaled 1.9 Mbp in length (0.15% of total assembly length). Approximately 5% (61 Mbp total length) of the Aaf assembly was not syntenic with AaegL5. We note two relatively large inversions on chromosome 1—one located at 1*p*34 and one at 1*q*42 (1.63 and 1.84 Mbp long, respectively; Fig. 3; Supplementary Table S5). We also detect a series of smaller inversions on the telomeric end of the q-arm of chromosome 1. Additionally, we detected smaller inversions on chromosome 3 located near 3*p*44 and two located on the telomeric end of the q-arm. As noted in the methods, we make no attempt to interpret nucleotide synteny or structural variation at the nucleotide level between Am and AaegL5 because they are too diverged. Nevertheless, we show the synteny map between these assemblies in supplementary figure S2.

**Fig. 3.**
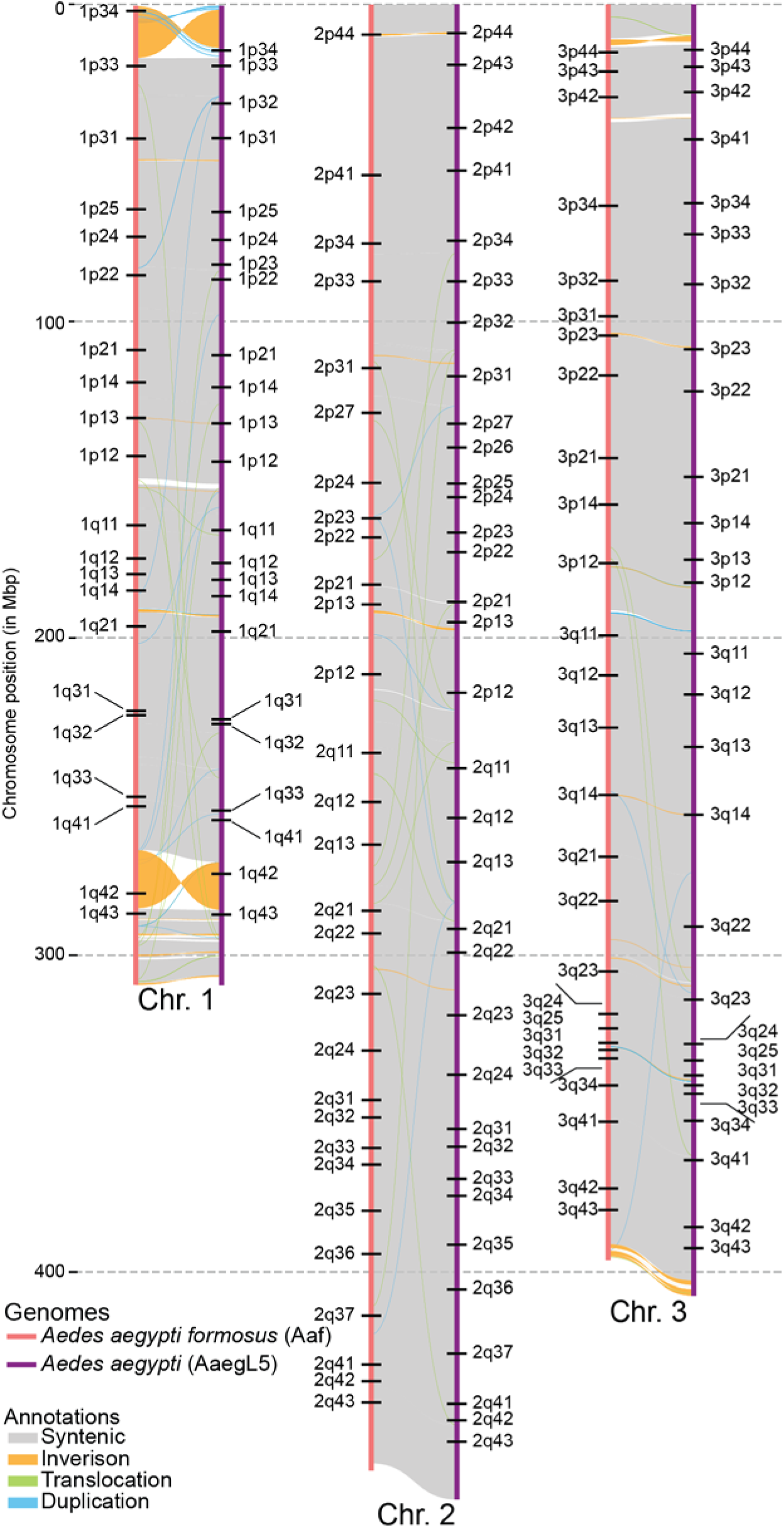
Synteny between chromosomes of *Aedes aegypti* (AaegL5) and *Ae. aegypti formosus* (Aaf). Bacterial artificial chromosomes (BAC; Matthews et al. 2018) positions represented as black horizontal bars.

### Gene order synteny across the Culicidae

Our analysis of gene order synteny revealed largely conserved patterns of chromosome evolution within mosquito clades, but patterns between clades showed substantial chromosomal rearrangement (Fig.5; Supplementary figure S6). Within the Aedini (*Aedes* and *Armigeres*) and Culicini (*Culex*), our analysis showed largely syntenic patterns (i.e., similar gene order) between the assemblies. However, *Anopheles* mosquitoes showed substantial chromosomal rearrangements between the subgenera that we included (Fig. 5). We found whole-arm translocations between chromosomes 2 and 3. In fact, translocation whole-arm translocations between chromosomes 2 and 3 appear to be pervasive when viewed at greater evolutionary scales. Without account for the translocation of chromosome 1 genes in Aedini onto chromosome 3 in *Anopheles,* we find five distinct arm associations in chromosomes 2 and 3 (Fig. 5; Supplementary figure S6)—one arm association among the Aedini, one for *Sabethes*, one for *Culex*, one for *An. cruzii* and *An. darlingi*, and one for *An. gambiae.* Interestingly, our analysis suggests *An. ziemanni* have the same autosomal arm associations as the Aedini while maintaining the same chromosome composition as other *Anopheles* (Fig. 5; Supplementary figure S6). Our analysis showed largely syntenic gene order arrangement between AaegL5, Aaf, and Am. This analysis found a similar set of inversions between AaegL5 and Aaf on chromosome 1 (Fig. 5; Supplementary figure S6). The same regions appeared to be inverted between Aaf and Am (Fig. 5; Supplementary figure S6). We detected an additional inversion between Aaf and Am at the distal end of the q-arm of chromosome 1. Lastly, we note that *P. papatasi* has N=5 chromosomes—three macrochromosomes (> 40 Mbp) and two microchromosomes (< 20 Mbp), but only three macrochromosomes are shown in figure 5. This was likely because too few orthologs were detected on the microchromosomes to adequately assess synteny. Nevertheless, we detected considerable gene order rearrangement in *P. papatasi* (Fig. 5)—genes originating from AaegL5 chromosome 2 comprised much of chromosomes 2 and 3 in *P. papatasi*, while chromosome 1 of *P. papatasi* was composed of genes originating from chromosomes 1 and 3 of AaegL5.

### Gene family evolution in the Culicidae

We found 6,559 common orthologs between the twelve species included in our analysis (Supplementary figure 7), 3,537 of which were single copy (Fig. 5). We found that both Aaf and Am had more orthologs in total (17,672 and 17,009 respectively) when compared to AaegL5 (14,626) (Fig. 5). We also found that Aaf and Am had more orthologs exclusively in common with one another than with AaegL5 (Fig. 5). Our analysis found 12,924 multi-copy orthologs in Aaf, 12,639 multi-copy orthologs in Am, and 10,538 multi-copy orthologs in AaegL5.

Furthermore, we detected 639 paralogs unique to Aaf, 447 paralogs unique in Am, and 285 unique paralogs in AaegL5 (Supplementary figure S7. Our analysis failed to assign orthology to 639 genes in Aaf, 447 genes in Am, and 285 genes in AaegL5 (Supplementary figure S7).

Across the culicid assemblies we included, *Ar. subalbatus* had the highest ortholog count (19,040), followed by Aaf and Am (Supplementary figure S7). These three assemblies also comprised the top three in terms of unique paralogs and unassigned genes. Anophelines had 138 orthologs exclusively common among them, while species in the Culicini had 320 orthologs exclusively common among them. At more granular evolutionary scales, *Culex* species exhibited 953 exclusive orthologs, while *Aedes* species exhibited 218 exclusive orthologs.

We used the maximum likelihood phylogeny output from IQ-Tree2 with 8,742,672 sites aligned across 3,534 partitions for the taxa in our analyses. We re-rooted the outputted phylogeny using *P. papatasi* as the outgroup, which showed strong monophyly of the Culicidae (Fig. 5). The topology of our phylogeny was largely congruent to that of Soghigian et al., (2023) with the exception of the placement of *Sabethes* (Fig. 5). Our phylogeny placed *Sa. cyaneus* (thus, Sabethini) as sharing a more recent common ancestor with the Aedini than the Culicini, contrary to the results of Soghigian et al. (2023), and more similar to those of Reidenbach et al. (2009).

Our analysis of gene family expansion and contraction showed substantial variability in gene family gains and losses, the majority of which occurred at the species level (Fig. 5). In addition to raw numbers of gains and losses of gene families, rapid gene family evolution (i.e., gene families with statistically significant changes in count AND categorized to have higher than average rates – reflected by blue numbers to the right of nodes in Fig. 5) appears to have happened at or near the tips rather than toward the root (Fig. 5). Indeed, our analysis showed that deeper internal nodes tended to have very few quickly evolving gene families than more relatively shallow nodes, such as the nodes leading to *Aedes* and the ancestor of *Culex* (Fig. 5). Despite the lack of quickly evolving gene families, our analysis did suggest an overall gene family expansion in the subfamily Culicinae, and contraction in the subfamily Anophelinae (Fig. 5). Between tribes in the Culicinae, our results indicated that the Aedini saw much greater gene family expansion than contractions (Fig. 5). Of the three *Aedes* genomes, both Aaf and AaegL5 saw roughly similar number of gene family contractions and expansions, Am saw a higher number of expansions than contractions (Fig. 5). Among tips, Aaf and AaegL5 have the highest number of rapidly evolving genes—most other taxa had roughly half or fewer rapidly evolving genes (Fig. 5). The distribution of these appear to differ between assemblies, though in general, rapid changes in gene count appear to be gains (Supplementary figure S8). Notably, the assemblies that exhibit rapid losses tend to be those derived from laboratory strains (i.e., AaegL5, *Cx. pipiens pallens*, *Cx. quinquefasciatus*, *Sa. cyaneus;* Supplementary figure S8).

We assessed the biological process gene ontology (GO) terms associated with each gene family and highlighted the most reoccurring, significantly expanded or contracted gene families with GO annotations by total number of copies represented among the species included. In general, while many gene families have expanded and contracted since diverging from each taxon’s recent common ancestor, we did not detect any changes that would be indicative of a pattern particular of any group of taxa (Supplementary figure S9), reflecting that most differences appeared to be between tips, rather than between genera or higher taxonomic rankings. In our analysis, we detected two sets of orthologs encoding odorant receptors, totaling five instances of significant count changes (Supplementary figure S6). We detected two different orthologs of rho-guanine exchange factor-related protein that had three total instances of significant copy number changes (Supplementary figure S9). E3 ubiquitin-protein ligase trip12 and glucose-methanol-choline oxidoreductase were each assigned to two different orthologs, each significantly changing in copy number once (Supplementary figure S9). We found seven other protein families that were each assigned orthologs whose copy number significantly changed twice: cyclic nucleotide-gated cation channel subunit A, fatty acid acyl transferase-related, malic enzyme-related, nipped-b-like protein delangin SCC2-related, oxidoreductase Glyr-1-related, scaffold attachment factor B-related, and voltage gated potassium channel (Supplementary figure S9).

### Comparison of gene ontology between Aaf, Am, and AaegL5

We assessed gene function at a finer scale across the three *Aedes* genomes by assessing significantly overrepresented GO terms among the set of significantly expanded genes common to these genomes. We used K-means clustering to group the 194 GO-terms common across the three genomes into 11 clusters, three of which appeared to describe metabolic processes (Fig. 6). The remaining eight clusters were loosely described as processes vital to behavior—sensory perception and detection of chemicals, ion transport, male mating and reproductive behavior, synaptic signaling and signal transduction, and cellular organization and biogenesis (Fig. 6). We did not detect any notable commonalities between the taxa and the sets of overrepresented GO terms (Fig. 6). When we repeated the overrepresentation test with all three taxa in a single analysis, we found similar results, albeit with substantially fewer GO terms overrepresented (Table 2; Supplementary table S6).

**Table 2.**
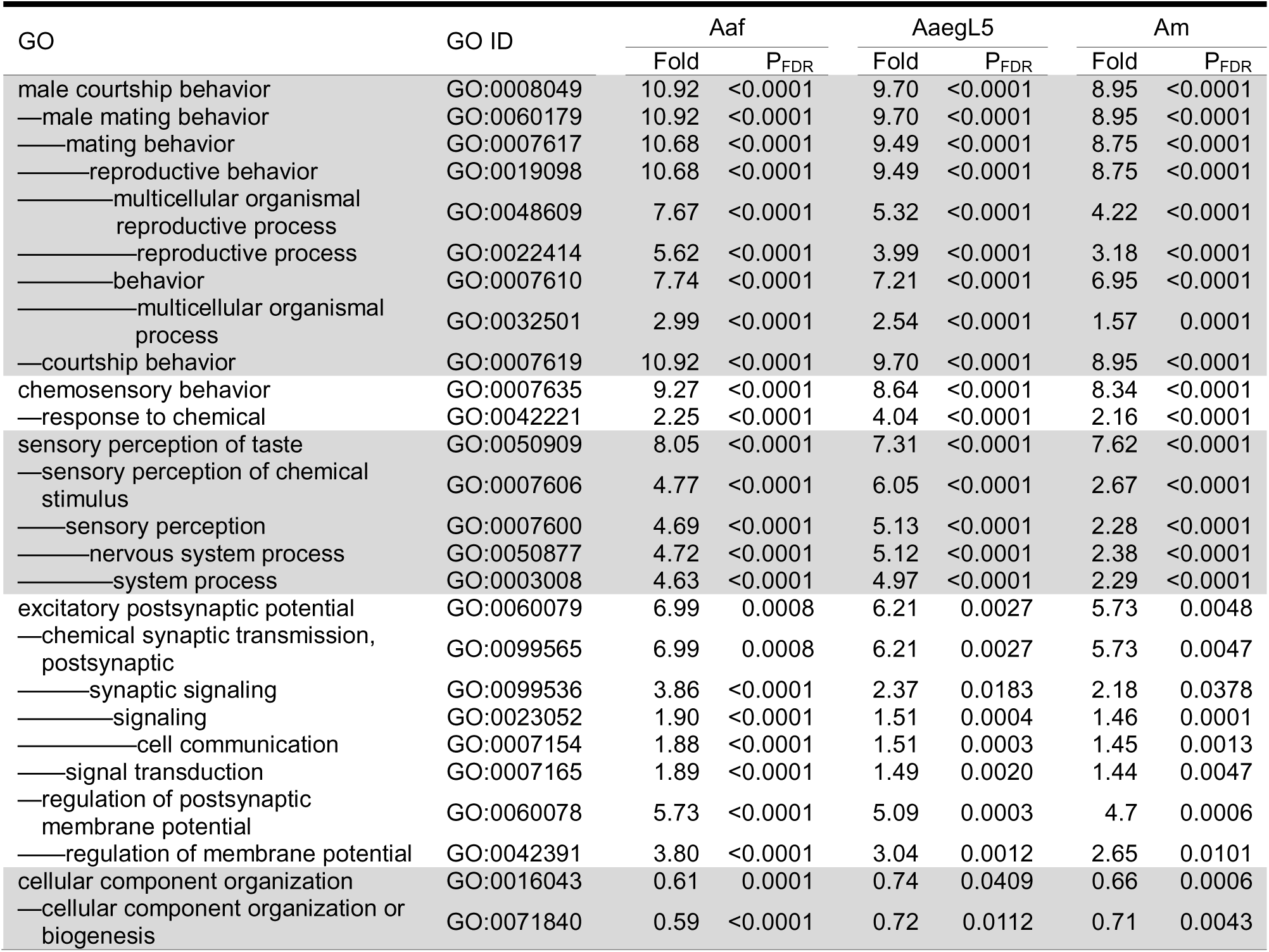
Overrepresentation test of terms common to all three Aegypti group assemblies (*Aedes aegypti formosus:* Aaf; *Ae. aegypti* AaegL5; *Ae. mascarensis* Am) done on the PANTHER DB web interface (release 20240807). The reference annotation set is from *Anopheles gambiae*. Significance testing done using Fisher’s Exact Test and the resulting P-values were corrected using false discovery rate (FDR). Here, ‘Fold’ refers to fold enrichment calculated as the actual number in the sample divided by the expected number from the same relative to the reference set. Here, the em-dashes (“—”) reflect the nested nature of GO terms and alternating shading of the rows separate basal-most GO terms.

### Gene families located in putative inversions in Aaf

By taking advantage of the gene families identified, we mapped 413 genes in the putatively inverted regions of Aaf chromosomes (relative to AaegL5), of which 354 were also identified in AaegL5 (Supplementary table S7) Notable genes among those found (Supplementary figure S10; supplementary table S8) were different odorant receptors (*Or4*) odorant binding proteins (Obp 56a and d), and ion channels (Shaker, *NaCh*) [important for signal transduction (Bohbot et al. 2007; McBride et al. 2014; Matthews et al. 2016)]; E3 ubiquitin ligases (*RNF 19B* and *RNF 126*) [similar E3 ubiquitin ligases are implicated in susceptibility to flavivirus infection (Giraldo et al. 2020; Dubey et al. 2022)]; heat shock proteins [important for heat and dehydration tolerance (Zhao et al. 2009; Benoit et al. 2010, 2011)]; and cytochrome P450 and adult and larval cuticle proteins [important for insecticide resistance (Poupardin et al. 2010)].

## Discussion

### *De novo* assembly of wild, individual *Aedes* mosquitoes

Small body size and a high input DNA requirement have been major hurdles to producing high-quality, chromosome-scale genome assemblies from many wild insects. Recent advances in sequencing technologies that generate highly accurate, long-reads, like PacBio HiFi (Wenger et al. 2019), allowed us to obtain enough high quality reads from a single mosquito to use for the first *de novo* genome assembly of *Aedes aegypti formosus* and *Ae. mascarensis*, avoiding the need of rearing colonies in the laboratory to obtain sufficient material. One additional roadblock, albeit minor compared to issues such as DNA input requirements, is determining which combinations of varied software tools produces the best assembly, particularly when considering numerous genome assembly metrics. *asmidx* allowed us to overcome this hurdle, choosing the best assembly from a range of excellent assemblies produced by a variety of genomic tools. Combined with Hi-C aided scaffolding (Burton et al. 2013; Dudchenko et al. 2017), the resulting genome assemblies from our pipelines are both highly contiguous and highly complete (Table 2; Supplementary figures S3 and S4). Using these chromosome-level assemblies together with other high quality Culicid reference genomes, we conducted a series of phylogenomic and comparative genomic analyses. The phylogenomic analysis (Fig. 5) showed minor differences to those recently published (Soghigian et al. 2023), likely due to substantial differences in taxonomic (i.e., number of taxa and lineages) and genetic sampling (i.e., whole genomes vs. sequence capture). The comparative genomic analysis revealed notable structural differences across large phylogenetic distances (Fig. 4) and numerous insights on the evolution gene families in the Culicidae (Figs. 5 and 6, Supplementary figures S8 and S9). We detail the implications of these findings below.

**Fig. 4.**
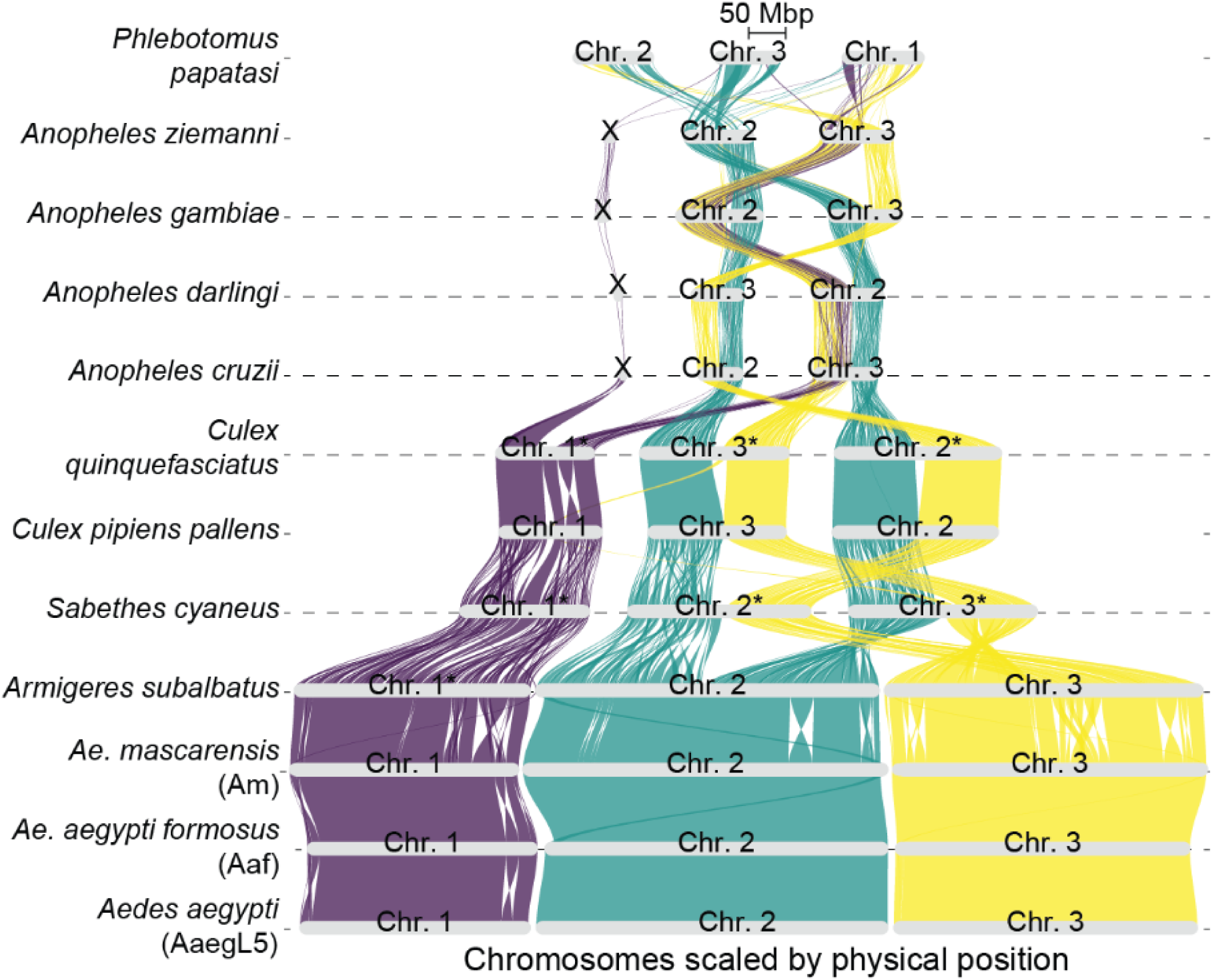
Riparian plot showing gene order synteny between eleven culicid genomes and the outgroup *Phlebotomus papatasi*. Synteny is assessed relative to the *Aedes aegypti* reference genome (AaegL5)—all genes originating from chromosome 1 on AaegL5 are shown in purple, all genes originating from chromosome 2 on AaegL5 are shown in green, and all genes originating from chromosome 3 are shown in yellow. Chromosomes with “*” in their names have been inverted to facilitate visibility. Similarly, some chromosomes appear out of order to facilitate visibility. Note that chromosomes 4 and 5 of *P. papatasi* are microchromosomes and not shown because too few orthologs were detected on them to adequately assess synteny with any of the chromosomes of AaegL5.

### Comparing Aaf, Am, and AaegL5

The assemblies of *Aedes aegypti formosus* (Aaf) and *Ae. mascarensis* (Am) had genome sizes comparable to AaegL5 assembly size (Supplementary figure S3; Table 1) and within range of genome size estimates from flow cytometry (Matthews et al. 2018). Both assemblies exhibited comparable gene content and accuracy to the *Ae. aegypti* reference genome, AaegL5 (Supplementary figure S3; Table 1) with a high degree of synteny between homologous chromosomes (Fig. 3). However, we also found evidence of inversions on Aaf chromosomes relative to AaegL5, though they require further testing via PCR for veracity. Recent studies (Redmond et al. 2020; Liang et al. 2024) have described numerous inversions in each of the chromosomes African and global populations of *Ae. aegypti*. In our *Ae. aegypti formosus* assembly (Aaf), we detected several of the same relatively large inversions that they discovered on 1*p*34, 1*q*42, and 3*q*43 (Fig. 3; Liang et al. 2024). The inversions on 1*p*34 and 3*q*43 are common among African populations of *Ae. aegypti* (referred to as 1pA and 3qG respectively by Liang et al. 2024). Furthermore, the inversion we detected on 1*q*42 is positioned similar to 1qF or 1qG detected in Burkina Faso populations Ouahigouya (OHI) and Ouaga-dougou (OGD).

Structural variations, such as inversions (Supplementary table S5), rearrange gene order, which in turn can lead to adaptive phenotypes that are shielded from recombination (reviewed in: Wellenreuther & Bernatchez 2018; Wellenreuther et al. 2019). This phenomenon is well-documented among *Anopheles* mosquitoes, wherein inversions are associated with numerous local adaptations (Powell et al. 1999; Cheng et al. 2018; Ayala et al. 2014) and genomic diversity (The Anopheles gambiae 1000 Genomes Consortium 2020). Inversions in *Ae. aegypti* have a long history of interest (Macdonald & Sheppard 1965) with recent research focusing on identifying inversions in different populations in both subspecies (Dickson et al. 2016; Redmond et al. 2020; Liang et al. 2024). The adaptive effect of these inversions remains unclear, however some genes that have been identified occur in inverted regions and have identified phenotypes. For example, odorant receptor 4 (*Or4*; McBride et al. 2014) located near the 1*q*42 position have been linked to preference for human odor. Other genes may have implications for vector management and adaptation. For example, over-expression and diversification of cuticle proteins are implicated in insecticide resistance in many insects (reviewed in: Balabanidou et al. 2018). Similarly, upregulation or increased copy number of heat shock proteins may contribute to more readily adaptable populations under increasing global temperatures (but see: Ware-Gilmore et al. 2023). A targeted, transcriptomic approach is necessary to further interrogate how genes in these inverted regions are expressed and their phenotypic consequences.

*Aedes aegypti* and *Ae. mascarensis* diverged approximately 8–10MY (Soghigian et al. 2020), thus we expected overall similarity between genome structure, especially in genic regions between the three *Aedes* assemblies. Indeed, our results showed a high degree of gene order synteny between these assemblies (Fig. 5) and the holistic set of GO term clusters show key clusters of genes with similar functions appear to be overrepresented (Fig. 6; Table 2; Supplementary table S6). For example, GO term clusters that describe mating and reproductive behavior as well as sensory perception appeared to be commonly overrepresented in all three taxa, perhaps because they are highly consequential to fitness (Cabrera & Jaffe 2007), and could represent differences in mating behaviors unique either to *Aedes* mosquitoes or to the Aegypti Group. However, an equally intriguing observation is that while many of the overarching clusters (i.e., mating/reproductive behavior, sensory perception, metabolism) are similar, the specific set of genes and GO terms assigned to them appear to vary in the Aegypti group, even between Aaf and AaegL5 (Fig. 6). These differences may be the manifestations of local adaptations—to environmental conditions where the mosquitoes were sampled (for Aaf and Am) or to the laboratory (for AaegL5). Indeed, numerous studies have documented local adaptations in *Aedes aegypti aegypti* to climatic/environmental (Soudi et al. 2023), altitude (Kramer et al. 2023), and vector competency for dengue serotypes (Lambrechts et al. 2009). Another possibility is that the source population for AaegL5 is not necessarily representative of wild *Ae. aegypti* (Gloria-Soria et al. 2019). Expanding the taxonomic sampling to other members of the Aegypti group may shed light on what functional genetic differences exist within the group.

**Fig. 5.**
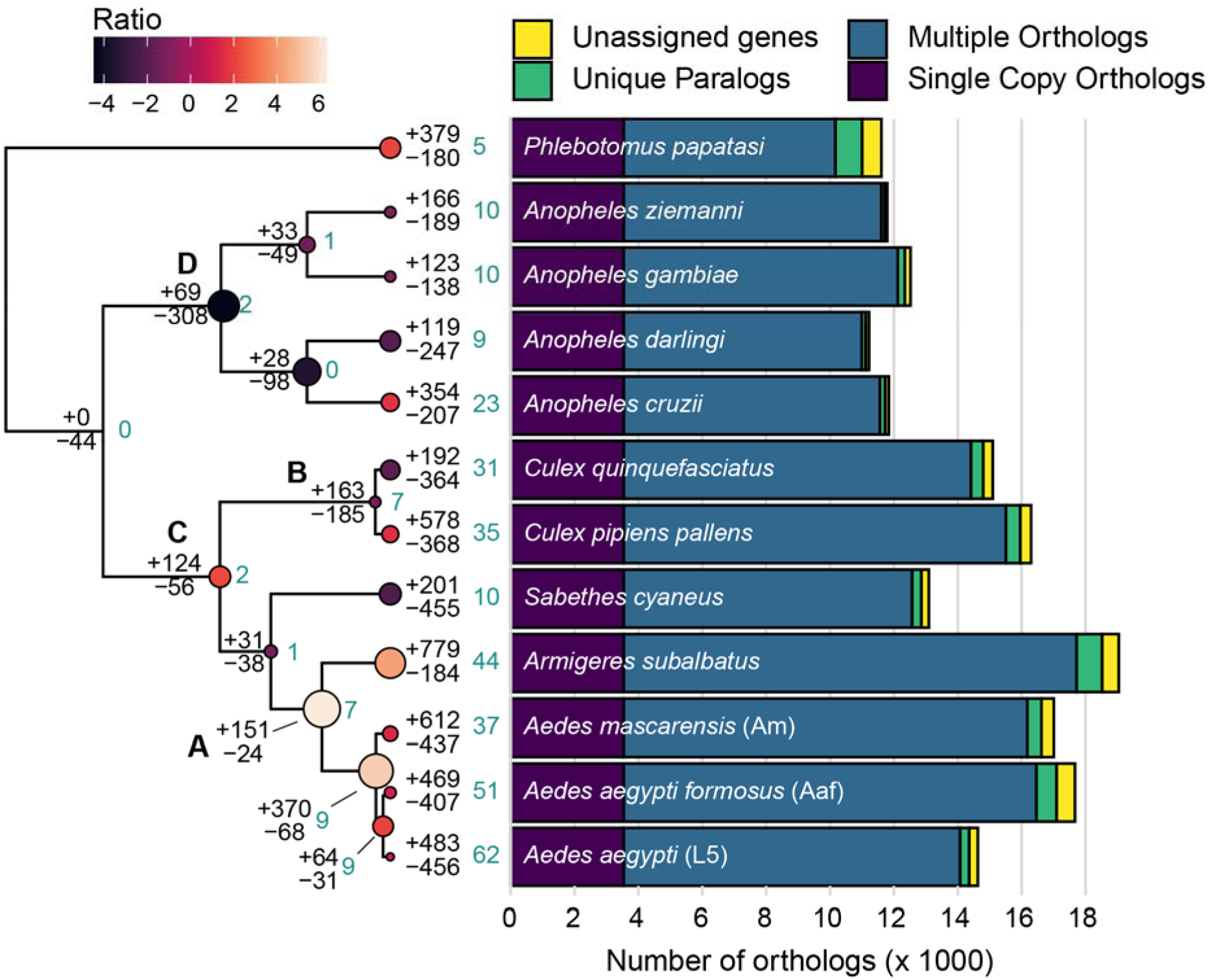
Maximum likelihood phylogeny output from IQTree2, re-rooted (to *Phlebotomus papatasi*) and ultrametricized using the *ape* package for R. Values shown at all nodes and tips represent rapidly evolving orthologs (blue) and expansions or contractions. All circles represent the ratio of expansions and contractions of ortholog copy number at each node and tip. Size of each circle represents the magnitude of the ratio (calculated as the greater of the two numbers divided by the lesser), while color represents direction (contraction-biased ratios are negative and darker, expansion-biased ratios are lighter and positive). Bold letters denote ancestral nodes for taxonomic groups: Aedini (A), Culicini (B), Culicinae (C), Anophelinae (D).

**Fig. 6.**
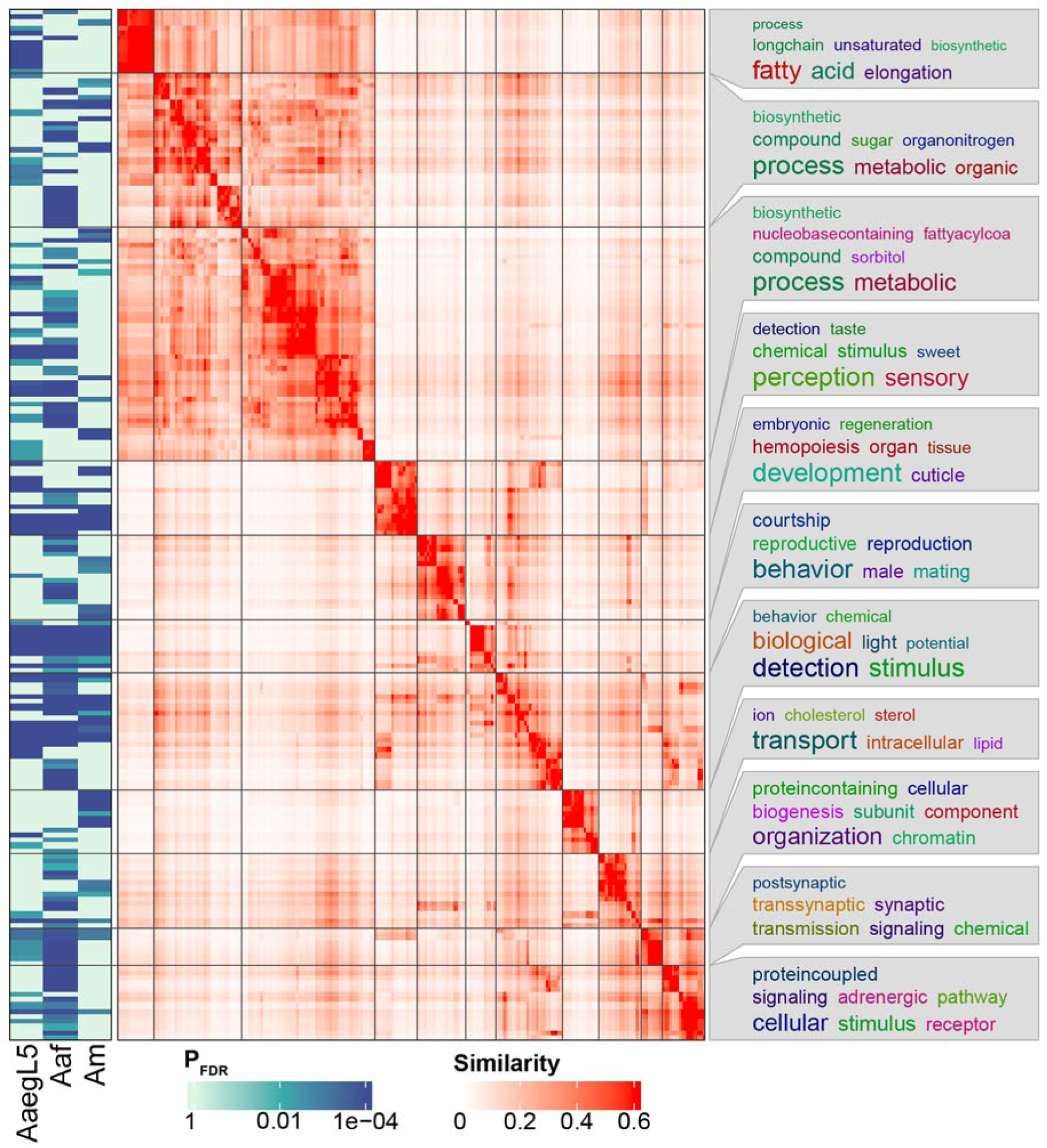
Heatmaps of GO term similarity and overrepresentation for the *Aedes aegypti* reference genome (AaegL5), *Ae. aegypti formosus* (Aaf), and *Ae. mascarensis* (Am). Similarity of 194 GO terms were assessed using semantic similarity outlined by Wang et al. (2007) and used K-means clustering to form eleven clusters of GO terms. Significance (P_FDR_ < 0.01) of overrepresentation of each GO term is shown in the heat map to the left. The heat map on the right shows similarity of individual GO terms, wherein GO terms are clustered together by similarity and divided by horizontal and vertical lines. The most commonly occurring descriptive terms for each cluster is shown to the right, where more frequent terms are shown in larger text.

An examination of the repetitive content in Aaf and Am found a notable departure from what was originally reported in AaegL5 (Fig. 2; Matthews et al. 2018). Indeed, whereas Matthews et al. (2018) found that 65% of the AaegL5 genomic sequence was considered repetitive, our assemblies showed nearly 80% of the genomic content to be repetitive. Other *Ae. aegypti* assemblies whose repeat content is characterized (Aag2 cell line: Whitfield et al. 2017; ROCK chromosome-scale assembly: Fisher et al. 2022) reported levels similar to Matthews et al. (2018). However, a recent re-examination found 78% of the AaegL5 genome to be repetitive DNA (Ryazansky et al. 2024) and thus similar to what we detected in Aaf and Am. The repeat landscapes for both Aaf and Am assemblies (Fig. 2 A, B) are similar to those reported by Whitfield et al. (2017), with sequence divergence peaking close to zero relative to the consensus sequences. These landscapes, particularly that of long terminal repeat (LTR) retrotransposons, represent recent activation and thus may represent recent infection from an RNA virus (Whitfield et al. 2017). In similar vein, the

### A Rickettsia endosymbiont found in Aedes aegypti formosus

We discovered an endosymbiont in our *Aaf* assembly (Supplementary figure S5). The genome size of this endosymbiont (1.59 Mbp) was similar to the Torix-group *Rickettsia* endosymbiont first detected in *Culicoides impunctatus* (Davison et al. 2022). While *Rickettsia* are better known for causing typhus fever and spotted fever (Raoult & Roux 1997), work conducted in the past 20+ years has revealed their extensive association with invertebrates and their tendency to manipulate host reproduction (see Perlman et al. 2006 and sources cited therein). This body of work has revealed their taxonomic diversity (Perlman et al. 2006; Weinert et al. 2009; Davison et al. 2022) and the diversity of their host range (e.g., Kikuchi et al. 2002; Thongprem et al. 2021), but their effects on hosts are still understudied. The most documented effect of many invertebrate-affecting *Rickettsia* is host reproductive manipulation, similar to those of *Wolbachia* (reviewed in: Werren 1997; Perlman et al. 2006). These endosymbionts are vertically transmitted from infected mother to her offspring, thus hijacking host reproduction to benefit themselves. These effects generally lead to a female-biased sex-ratio either by killing males (Werren et al. 1994) or inducing parthenogenesis (Hagimori et al. 2006; Aguin-Pombo et al. 2021). Torix-group *Rickettsia* have a direct effect on body size of host leech species, wherein infected individuals exhibiting larger body sizes (Kikuchi et al. 2002), and low dispersal among infected spiders (Goodacre et al. 2006). The prevalence or the effect of this *Rickettsia* in *Ae. aegypti formosus* from Burkina Faso is unknown.

### Repeated genomic rearrangement in Culicidae

Recent advances in sequencing technologies, physical mapping, and three dimensional chromosomal structure inference have made evident the extensive evolutionary rearrangement of chromosomes in Culicidae (Sharakhov et al. 2002; Neafsey et al. 2015; Palatini et al. 2020; Yurchenko et al. 2023; Ryazansky et al. 2024; Lukyanchikova et al. 2022). Microscale structural variants that confer local adaptations (Ayala et al. 2014; Powell et al. 1999; Cheng et al. 2018) and macroscale whole-arm translocations detected between species are well-studied in *Anopheles* (Sharakhov et al. 2002, 2016; Neafsey et al. 2015; Wei et al. 2017; Artemov et al. 2018). In the Culicinae, Arensburger et al. (2010) detected whole-arm translocations between *Ae. aegypti* and *Cx. quinquefasciatus,* which Ryazansky et al. (2024) recently confirmed. Our analysis expand on the scope of these studies by including more taxa from across the Culicidae and show several intriguing trends. First, associations of chromosome arms have repeatedly changed throughout the evolutionary history of the Culicidae (Fig. 4; Supplementary figure S6). Indeed, relative to AaegL5, all non-Aedini genomes we investigated showed whole-arm translocations between chromosomes 2 and 3 (Fig. 4, Supplementary figure S6; also see: Arensburger et al. 2010; Neafsey et al. 2015; Ryazansky et al. 2024). Second, in stark contrast to our first point, chromosomes 2 and 3 showed stability in the Aedini, as none of the assemblies in the tribe showed whole-arm translocations (Fig. 4; Supplementary figure S6). Our taxonomic sample cover roughly 50MY of evolutionary history in the Aedini, and in a similar timeframe, each of the anophelines in our dataset evolved to exhibit unique chromosomes 2 and 3 arm associations, again in line with the findings of Neafsey et al. (2015), wherein syntenic blocks rapidly decayed in that timeframe. Lastly, despite multiple major rearrangements, most culicids exhibit a karyotype 2N = 6, with the sole exception being *Chagasia bathana* (subfamily Anophelinae), where 2N = 8 (Rai & Black 1999). This level of conservation is remarkable in the context of other arthropods such as Coleoptera (Blackmon et al. 2024), Lepidoptera (Wright et al. 2024), and within Diptera (Morelli et al. 2022). To better-interrogate chromosome evolution within Culicidae, more high quality, chromosome-scale genome assemblies are required, especially within the Culicinae.

### Evolutionary changes in distribution and copy number of gene families

Changes in copy number of key gene families may play key adaptive roles in mosquitoes leading to differences in vector effectiveness between species (Arcà et al. 2017; Palatini et al. 2017; Catapano et al. 2023) and populations (Lambrechts et al. 2009; Bennett et al. 2021). We compared our *Ae. aegypti formosus* (Aaf) and *Ae. mascarensis* (Am) genomes to high quality mosquito genome assemblies that were publicly available and found striking differences both in the distribution of rapidly evolving orthologs (Supplementary figure S8) and ortholog copy number (Supplementary figure S9).

Rapid changes in copy number may also be an indication of adaptation (Simon et al. 2015; Xie et al. 2018). Rapid gains, in particular have been attributed to adaptative evolutionary changes, as duplicated gene copies are “released” from stabilizing selective pressures may respond adaptively as the environment or the context in which they are expressed changes (Guo & Kim 2007; Vieira et al. 2007), however other works have also shown rapid losses to also lead to adaptation (McBride & Arguello 2007; Goldman-Huertas et al. 2015). Rapid gains in orthologs appear in all mosquito assemblies in our data set (Supplementary figure S8), however rapid losses appear to have happened more often among laboratory strains. It is unclear whether this is a pattern of adaptation to laboratory conditions (Gloria-Soria et al. 2019; Ross et al. 2019) or an artifact (e.g., sampling bias) of the available genomic resources of mosquitoes. A more detailed examination with more diverse sampling of laboratory strains would be necessary if there is a tendency for rapid gene loss among laboratory strains compared to wild populations.

At more granular levels, we detected eleven different reoccurring rapidly evolving gene families in our data set (Supplementary figure S9). Among them, we detected those involved in sensory processes (odorant receptors) and signaling cascades (CNG cation channels) reoccurred most often (Fig. 5). These proteins are crucial for detecting and transducing olfactory signal (Zwiebel & Takken 2004; Sato et al. 2008) and thus key for host detection. Our analysis showed two sets of orthologs that encode odorant receptors—one that has contracted in *Cx. quinquefasciatus* and *Ar. subalbatus* but expanded in *Cx. pipiens pallens*, and another that has expanded in *Cx. quinquefasciatus* and *Ae. aegypti* (AaegL5) (Supplementary figure S9). No clear pattern of blood host affinity (based on: Soghigian et al. 2023) arises from these combination of taxa and the orthologs we detected.

## Conclusion

Here, we presented a *de novo* genome assembly of a wild caught *Ae. aegypti formosus* (*Aaf*) and *Ae. mascarensis* (*Am*) each derived from a single, wild-caught individual. Our assemblies are comparable to the reference *Ae. aegypti* assembly (AaegL5; Matthews et al. 2018) in terms of contiguity and gene content but differ in that the Aaf assembly exhibits genomic structural variation particular to West Africa (Liang et al. 2024), and notable differences in ortholog counts and their functions. At 8–10 MY diverged, we view the nucleotide synteny between Am and AaegL5 as unreliable, but gene order in *Ae. mascarensis* is highly conserved and show a high degree of synteny with *Ae. aegypti*. With the three *Aedes* genome assemblies we also showed variation in gene family expansion, and the function of those expanded gene families, showing population- (between AaegL5 and Aaf) and species- (between *Ae. aegpyti* and *Ae. mascarensis*) wide differences. These assemblies will be valuable assets for future studies to understand the biology and evolution of *Ae. aegypti*—the *Aaf* assembly more closely reflects natural populations of *Ae. aegypti* in the ancestral range, and the Am assembly provides the closest genome output to date for this species. We used our newly assembled genomes along with ten other reference genomes evolutionary changes within the Culicidae and find repeated bouts of major chromosomal rearrangements, particularly between chromosomes 2 and 3. The genomes we present here represent initial steps toward the development genomic resources for all of the currently described taxa in the Aegypti group (Soghigian et al. 2020). Further development within this group can elucidate genomic architecture that differentiates the ecology and behavior of the African and the global invasive sub-species.

## Data Availability

The PacBio HiFi reads generated for this project will be deposited in GenBank within BioProjects (Am_MascCH02 principal: PRJNA1199517, Am_MascCH02 alternate: PRJNA1199516, Aaf_Bf05 principal: PRJNA1199519, Aaf_Bf05 alternate PRJNA1199518) under SRA accession XXXXX and XXXXX. The GenBank accession number for the assemblies we generated are: Am_MascCH02 principal: XXXX, Am_MascCH02 alternate: XXXXXX, Aaf_Bf05 principal: XXXXXX, Aaf_Bf05 alternate: XXXXXXX, and all available at NCBI.

## Supporting information

Supplementary Results

## Acknowledgements

Financial support for this project was provided by NIAID R01 AI101112 awarded to JRP. We also acknowledge the support of the Natural Sciences and Engineering Research Council of Canada (NSERC), funding reference number RMS21-73779779 [Cette recherche a été financée par le Conseil de recherches en sciences naturelles et en génie du Canada (CRSNG), numéro de référence RMS21-73779779]. GM was supported by NIAID R01 AI155562 awarded to JRP, AG-S, and JS. We thank T. Petruff, J. Brophy, S. Arent, and R. Pellegrini for support with sample processing. We also thank L. Jackson for providing comments on an earlier version of this manuscript. Lastly, we thank the Research Computing Services group at the University of Calgary.

## Notes

### Competing Interest Statement

The authors have declared no competing interest.

### Summary of Updates

We have updated our author list, as previous versions unintentionally omitted a contributor. No changes to text have occurred.

