## Supplementary Results for "From macro to micro: De novo genomes of Aedes mosquitoes enable comparative genomics among close and distant relatives"

We generated twelve assemblies each for Aaf and Am—four directly output from the assemblers (*HiCanu*, *flye*, *hifiasm*, and *IPA*), and eight which resulted from passing assembler output into either *pd* or *ph* for purging duplicates and haplotigs from the assemblies. In general, assembly size metrics did not differ substantially between purging programs, with one exception (see below). When examining genome size relative to the AaegL5 (Matthews et al. 2018), results should approach a predicted genome size (Supplementary Table 4). Assembly outputs from *flye* and *HiCanu*—neither of which include a purging step—are nearly double the size of the AaegL5 reference (Supplementary Table 4). Only the *hifiasm* output for Am had a genome size that was reasonably close to AaegL5. In fact, all other draft assemblies directly output from assembly programs (i.e., no external purging) fell outside the range of genome size estimates from flow cytometry (Nene et al. 2007; Matthews et al. 2018). Thus, not only is purging a necessary step, internal purging alone also does not obviate the need for an external purging step in our case. In nearly all other cases purging duplicate contigs and haplotigs reduced genome sizes to within ±10% of the reference genome. *Am* assembly using *hifiasm* and purged using *pd* led to an over-purged genome, reducing its total length by nearly 500 Mbp. For both species, *hifiasm* output assemblies with the highest N50s. *HiCanu* and *IPA* show similar performance to one another, and *flye* output the lowest N50*.* In terms of the number of contigs that comprise each assembly, purged assemblies showed lower contig counts after purging with either *pd* or *ph*.

Since nearly all outputs from assemblers required external purging, it was important to assess whether external purging introduced errors into the purged assemblies. We did this using *Inspector*, by quantifying the number of base pairs associated with structural errors (i.e., >50bp) for all assemblies we generated. For both specimens, every combination of *IPA* generated assemblies with the greatest number of collapsed bases (Supplementary Table 4), while all other combination of assembler and purging program performed similarly. Base expansions generally increased for all assemblers after external purging, with Aaf assembled using *hifiasm* and purged using *ph* exhibiting substantial expansion (>300kb). *Inspector* also detected a substantial number of inverted bases, especially for *hifiasm* and *IPA* after external purging (Supplementary Table 4). Notably, *hifiasm* had the highest number of inverted bases for both specimens prior to external purging. Conversely, *flye* assembled the fewest inverted bases, with external purging only modestly increasing the number of inverted bases (if at all). While *HiCanu* also showed increased number of inverted bases after purging, the number of bases involved was substantially lower than that found in *hifiasm* or *IPA.*

We assessed gene content of all assemblies using *BUSCO*. Assemblies with no external purging generally had a high number of duplicated orthologs (Supplementary Table 4). Externally purged assemblies had similar numbers of complete, fragmented, and missing orthologs, though external purging modestly increased the number of missing orthologs compared to assemblies that were not purge externally. However, as noted above, the *Am* assembly output from *hifiasm* and purged using *pd* exhibited roughly 40% of orthologs missing, indicating that *pd* over-purged the assembly.

We feature normalized each of the assembly characteristics for each specimen to determine which of assemblies generated was considered the “best”, and thus a good candidate for scaffolding. Based on a weighted set of characteristics which included gene content, assembly accuracy, contiguity, and relative assembly size, we determined that the *HiCanu* assembly purged with *ph* was the best assembly for Aaf and the *hifiasm* assembly purged with *ph* to be the best Am assembly (Fig. 1).
