## Supplementary material for "From macro to micro: De novo genomes of Aedes mosquitoes enable comparative genomics among close and distant relatives": Figure S1

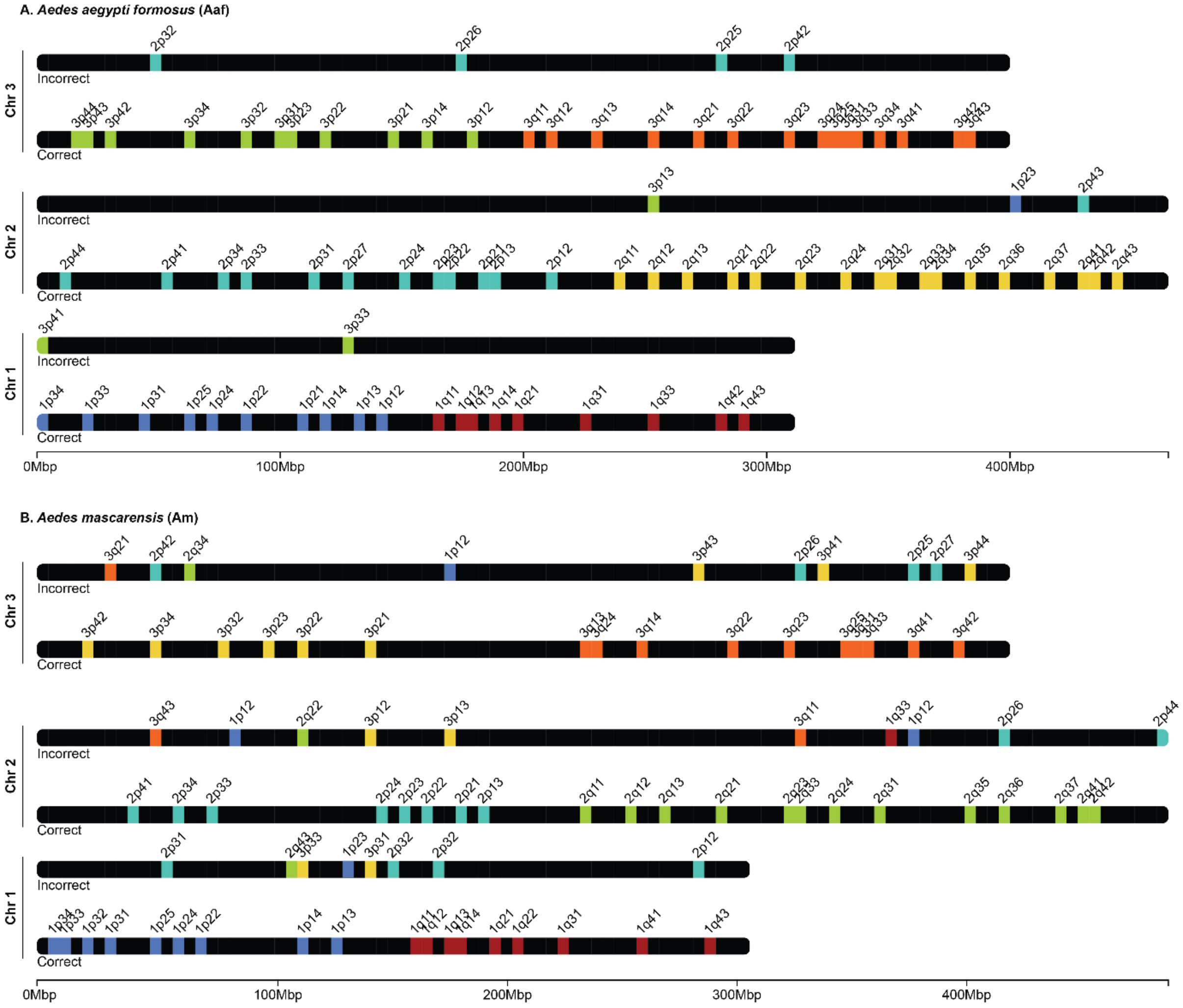


**Supplementary figure S1**. A simple visualization of the three primary (i.e. chromosome-level) scaffolds for *Aedes aegypti formosus* (Aaf; A) and *Ae. mascarensis* (Am; B) with bacterial artificial chromosomes (BAC) positions (from Matthews et al. 2018) mapped (color bars). Each scaffold is shown twice: the top scaffold shows the scaffolds with erroneously mapped BAC positions, while the bottom scaffolds show correctly mapped BAC positions. Each scaffold with correctly mapped BAC positions should have two colors, one each for the p and q arms of the chromosome, and the positional numbering should sequentially decrease from the telomeric end to the centromere for each p and q arm. Note that band width is not to scale.
