## Supplementary material for "From macro to micro: De novo genomes of Aedes mosquitoes enable comparative genomics among close and distant relatives": Figure S2

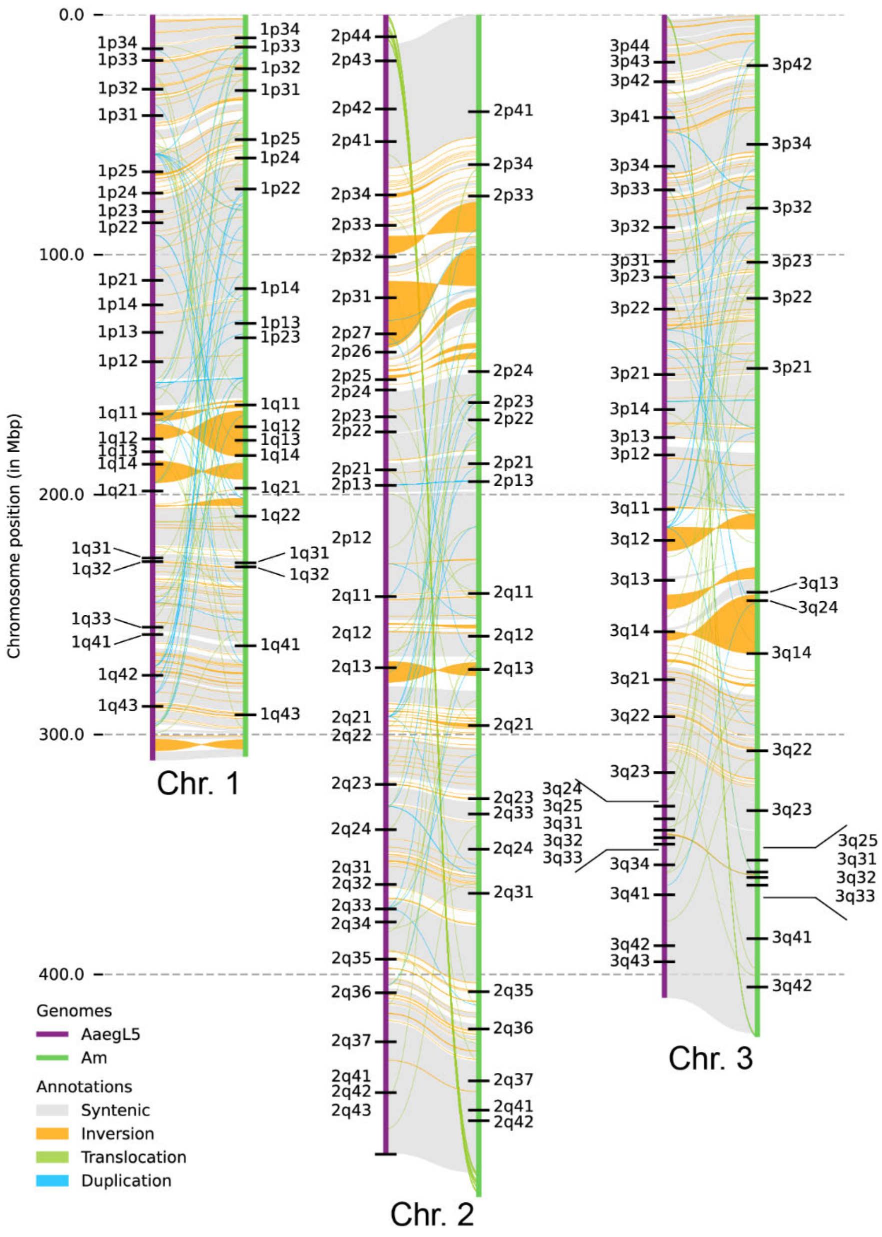


**Supplementary figure S2.** Nucleotide synteny between *Aedes aegypti* reference genome (AaegL5) and *Ae. mascarensis* (Am). The two species are approximately 8 MY diverged, and thus required more stringent settings for genome alignment (*minimap2* settings -asm20). As noted in the main text, we made no attempt to interpret this output because we deemed the two species to be too diverged from one another to reliably assess structural variation at the nucleotide level. Unlike nucleotide identity, gene order along the chromosomes appears to be more conserved (see Fig. 4 in the main text).
