## Supplementary material for "From macro to micro: De novo genomes of Aedes mosquitoes enable comparative genomics among close and distant relatives": Figure S3

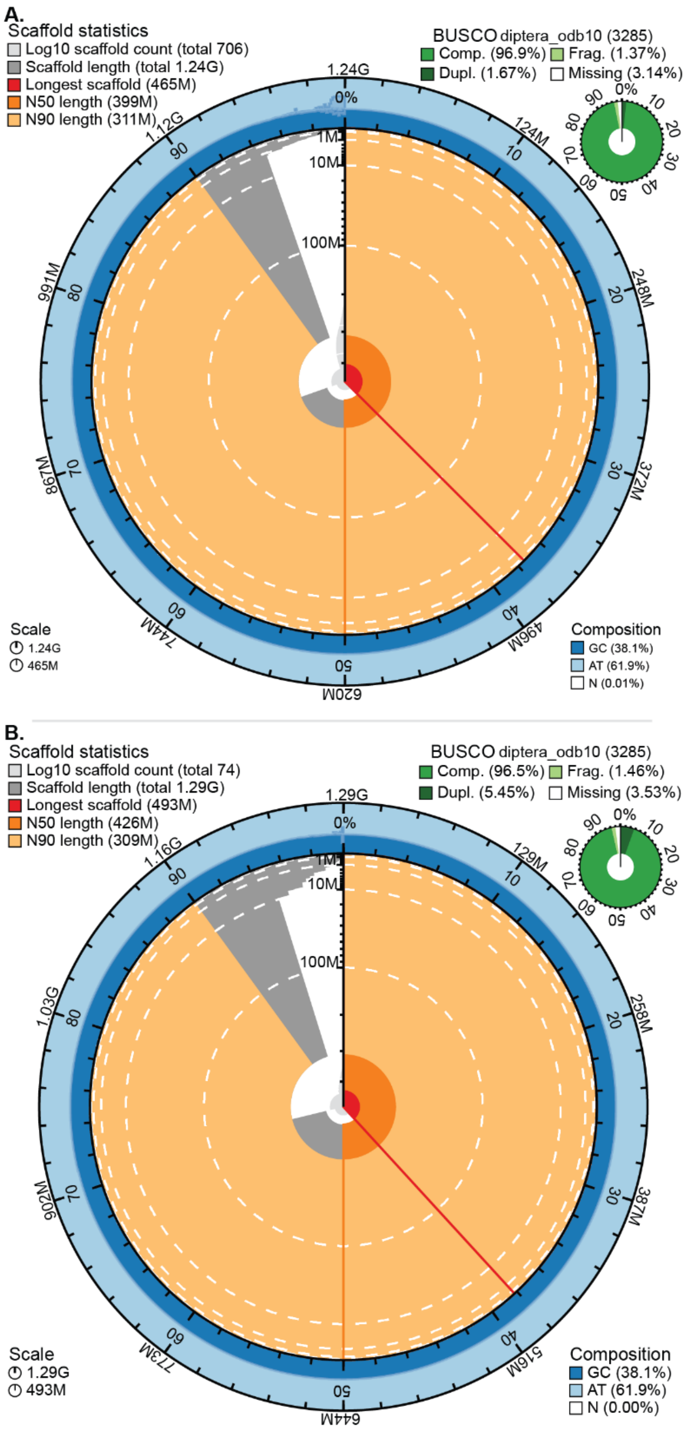


**Supplementary figure S3.** Snail plots for (A) *Aedes aegypti formosus* (Aaf) and (B) *Aedes mascarensis* (Am). The light blue and dark blue rings show AT and GC content along the length of the genome respectively. The sector lengths represent N90 (light orange), N50 (dark orange), longest scaffold (red), cumulative scaffold length (dark grey), and scaffold count (light grey). BUSCO scores based on diptera_odb10 are shown to the right.
