## Supplementary material for "From macro to micro: De novo genomes of Aedes mosquitoes enable comparative genomics among close and distant relatives": Figure S4

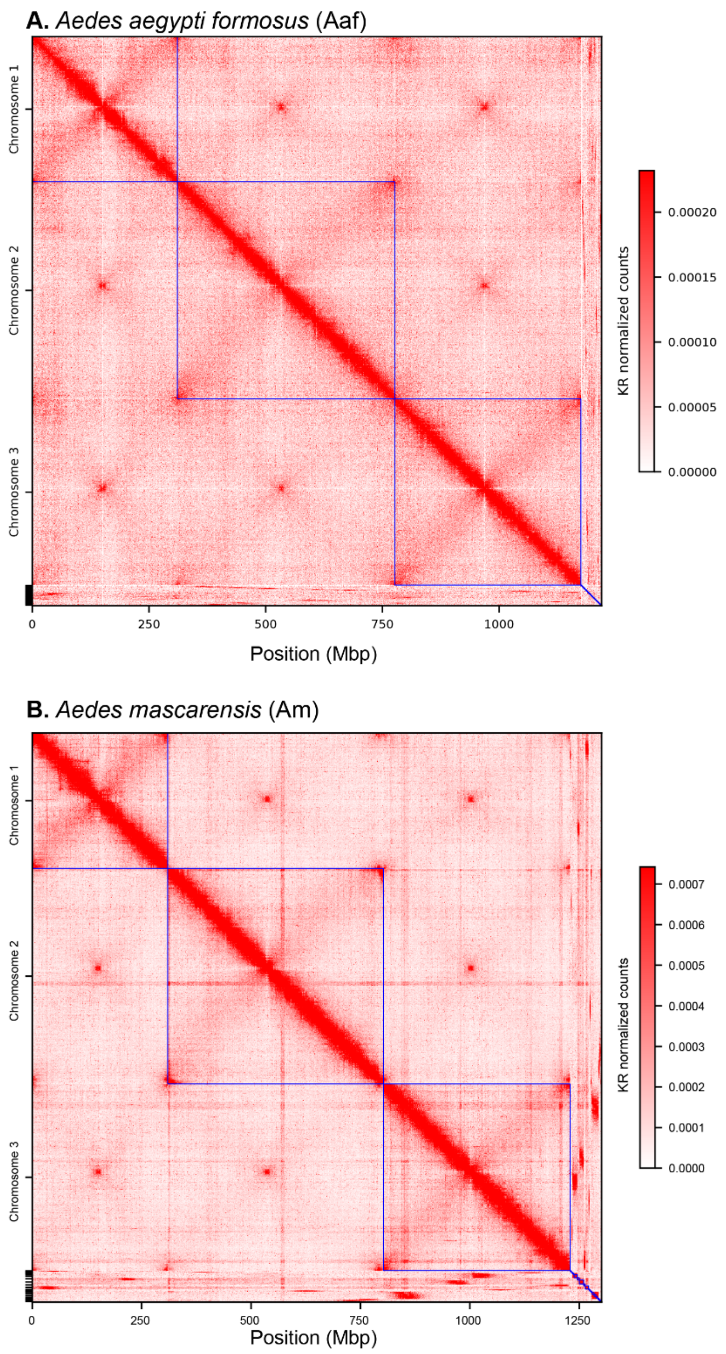


**Supplementary figure S4.** Finalized Hi-C contact map for (A) *Aedes aegypti formosus* (Aaf) and (B) *Ae. mascarensis* (Am). Blue boxes indicate boundaries of scaffolds. After mapping cross-linked paired-end short reads, we scaffolded both assemblies using *Yahs* and manually curated the assembly in *Juicebox*. We used HapHiC to visualize the final output shown here.
