## Supplementary material for "From macro to micro: De novo genomes of Aedes mosquitoes enable comparative genomics among close and distant relatives": Figure S5

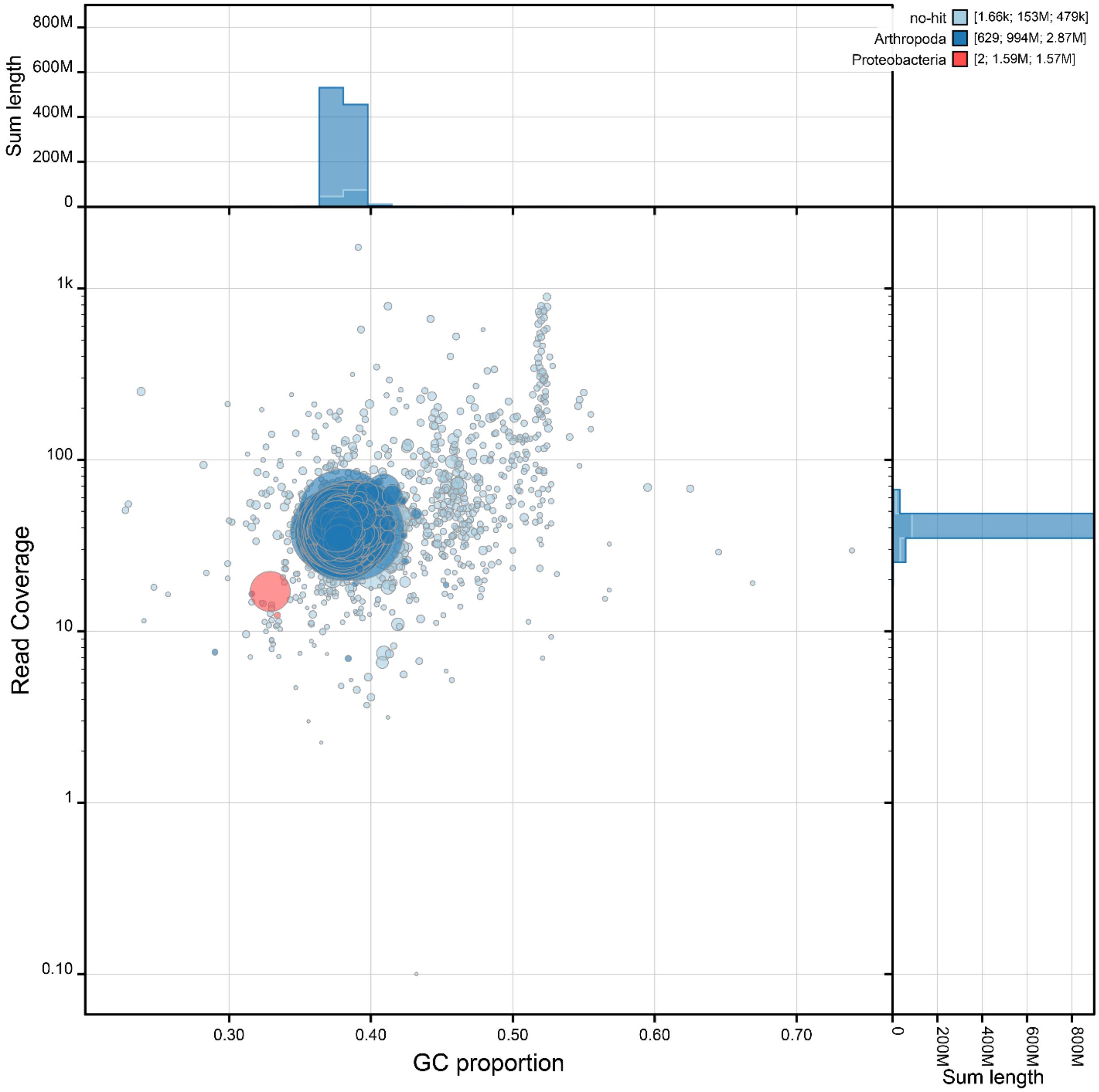


**Supplementary figure S5.** Bubble plot for the debris/duplicates of the *Aedes aegypti formosus* (Aaf) assembly. Bubble plot showing the taxonomic identities of each sequence in the concatenated fasta file containing duplicate haplotigs and debris contigs of *Aedes aegypti formosus* (Aaf). Each circle represents a contig or scaffold in the assembly, and its size represents its size. Two sequences shown in red originate from an α-proteobacterium endosymbiont which we determined to be *Rickettsia.*
