## Supplementary material for "From macro to micro: De novo genomes of Aedes mosquitoes enable comparative genomics among close and distant relatives": Figure S6

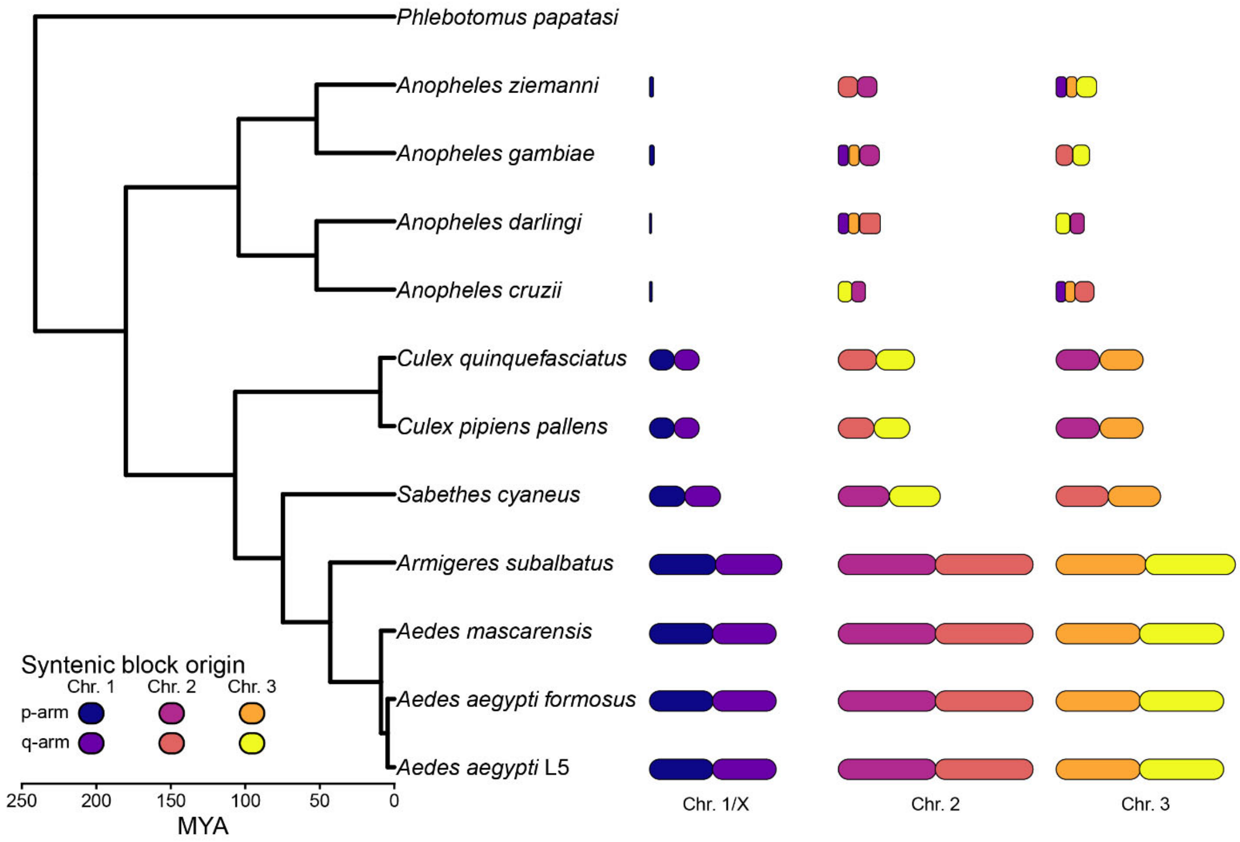


**Supplementary figure S6.** A simplified schematic representation of the chromosome arm arrangement detected in Figure 5 plotted alongside the maximum likelihood phylogeny inferred from IQTree2. The combined length of the chromosomes for each assembly are scaled to the *Aedes aegypti* reference genome (AaegL5), and the length of each chromosome is scaled to chromosome 2 of each assembly. Chromosome arms are colored according to their originating chromosome arms relative to AaegL5. For the *Anopheles* species, the x chromosomes all originate from the p-arm of chromosome 1 of AaegL5. Note that here, the chromosomes for all taxa are ordered numerically.
