## Supplementary material for "From macro to micro: De novo genomes of Aedes mosquitoes enable comparative genomics among close and distant relatives": Figure S7

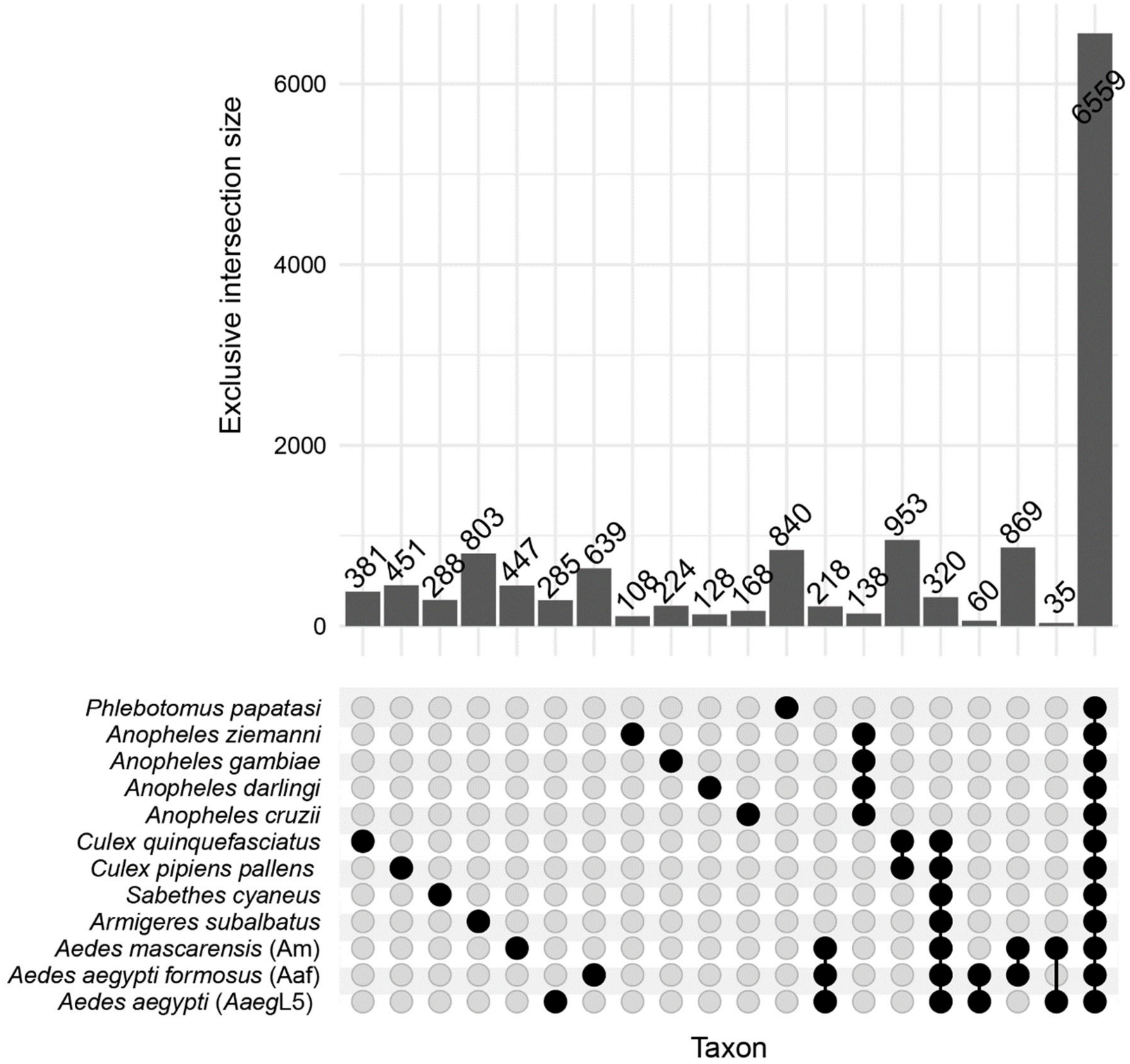


**Supplementary figure S7.** Upset plot of orthologs common among the twelve taxa included in the analysis. All unions between multiple taxa (i.e., dots connected by a line) are exclusive.
