## Supplementary material for "From macro to micro: De novo genomes of Aedes mosquitoes enable comparative genomics among close and distant relatives": Figure S8

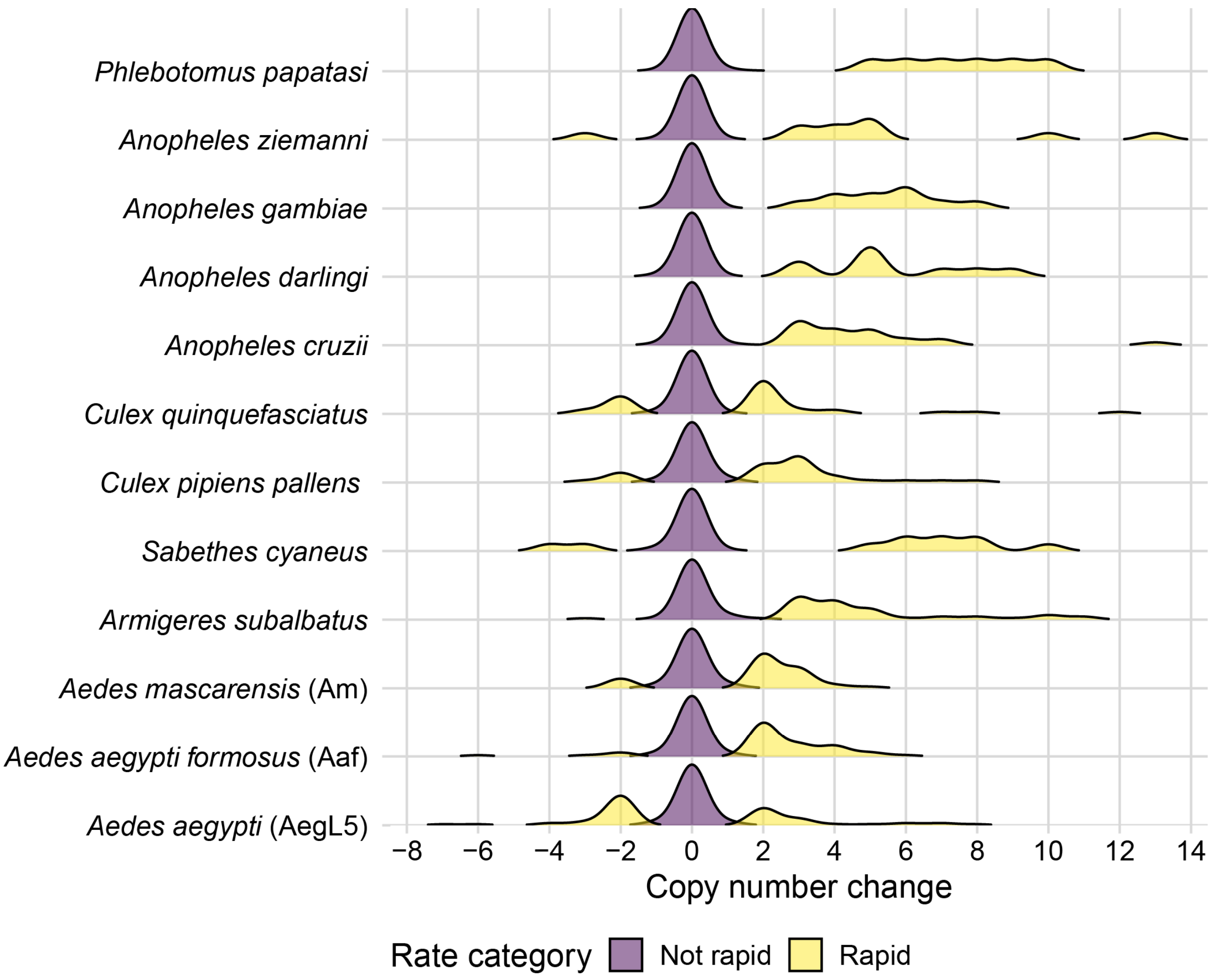


**Supplementary figure S8.** A density plot for each species showing the distribution of rapidly evolving orthologs modeled using a birth-process. Rapid or not rapid was determined using a γ-distribution with four rate categories in Cafe5.
