## Supplementary material for "From macro to micro: De novo genomes of Aedes mosquitoes enable comparative genomics among close and distant relatives": Figure S9

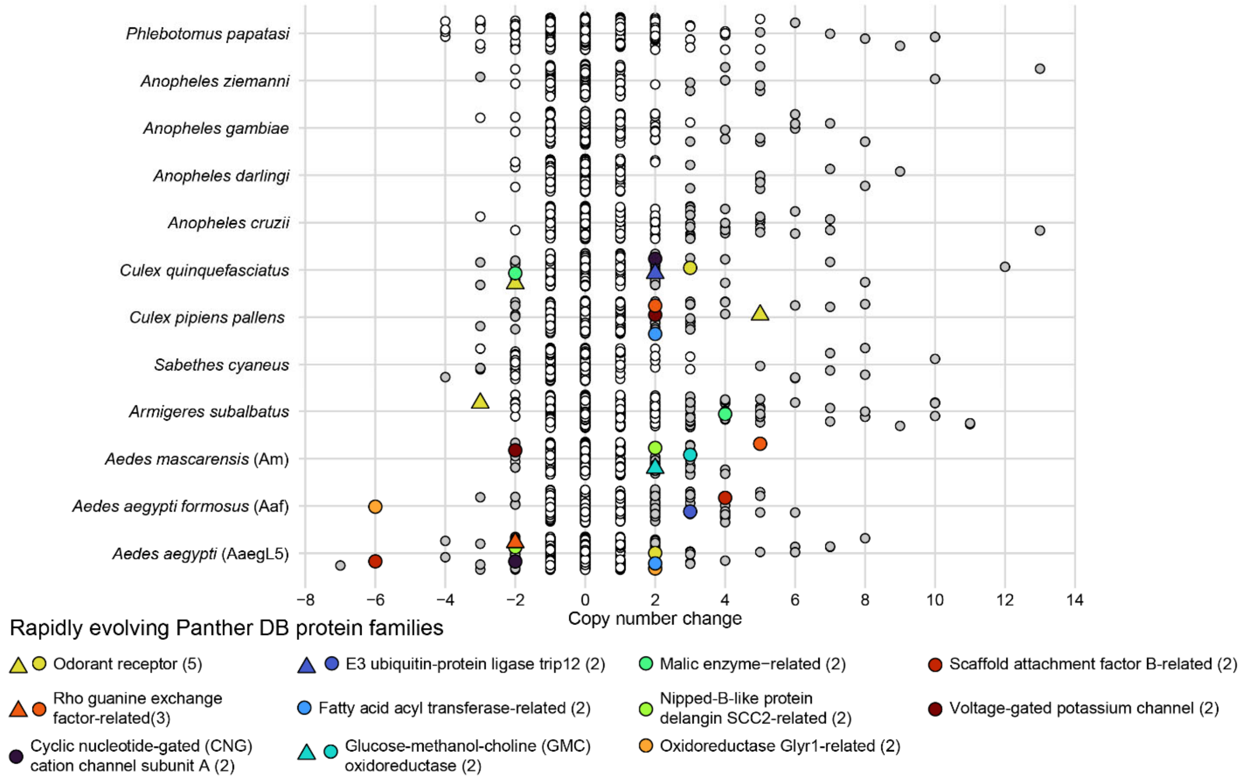


**Supplementary figure S9.** Dot plot showing each ortholog for each assembly. White dots represent orthologs whose copy numbers did not change significantly since diverging from its parent node. Gray dots are orthologs of taxa whose copy numbers have changed significantly since diverging from its parent node once. Colored shapes are orthologs of taxa whose copy numbers have significantly and rapidly changed more than once (shown in parentheses). When two different orthologs encoded a protein in the same family, they are represented by different shapes.
