## Supplementary material for "From macro to micro: De novo genomes of Aedes mosquitoes enable comparative genomics among close and distant relatives": Figure S10

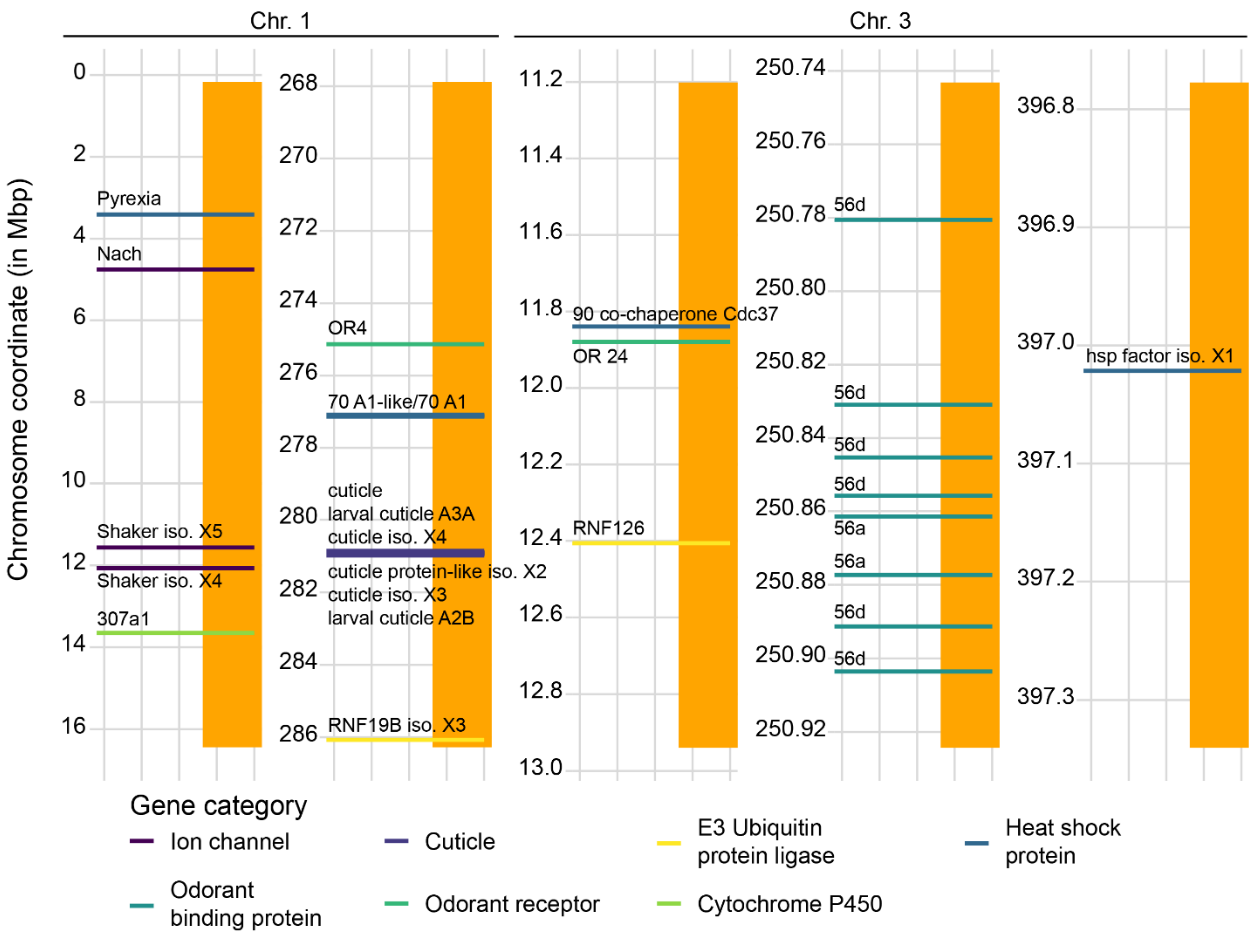


**Supplementary figure S10** Select genes found in inverted regions of chromosomes 1 and 3 of *Aedes aegypti formosus* (Aaf). Orange bar represents the inverted regions (relative to the reference genome AaegL5 in Fig. synteny plot) and colored lines show the start positions of each gene. Some lines are thicker than others due to the proximity of similar gene categories. Here, ‘iso.’ stands for ‘isoform’.
