## Supplementary material for "From macro to micro: De novo genomes of Aedes mosquitoes enable comparative genomics among close and distant relatives": Table S1

**Supplementary Table S1**. Programs used to assemble drafts genomes from PacBio HiFi reads.

| Program | Version | Flags | Notes |
|---|---|---|---|
| | Assembling from reads to contigs | | |
| HiCanu | 2.2 | -pacbio-hifi | |
| flye | 2.9-b1768 | --pacbio-hifi | |
| hifiasm | 0.16.1-r375 | --primary | |
| IPA | 1.8.0 | | local mode |
| mitohifi | 3.0.0 | -a animal -o 6 | |
| | Purging duplicated haplotigs | | |
| purge_dups | 1.2.5 | | |
| purge_haplotigs | 1.1.2 | -a 60 | |
| minimap2 | 2.23 | -I6G -x map-hifi (pd); -ax map-hifi (ph) | |
| samtools view | 1.15 | -hF 256 | As part of ph workflow |
| samtools sort | 1.15 | | |
| | Scaffolding | | |
| fastp | 0.23.4 | --html -f 5 --detect_adapter_for_pe | |
| bwa index | 0.7.17-r1198-dirty | | |
| bwa mem | | -a bwtsw | |
| samtools view | 1.15 | -h | |
| samtools view | 1.15 | -bS -t | |
| samtools view | 1.15 | -Sb | |
| samtools sort | 1.15 | | |
| picard | 2.27.4-SNAPSHOT | | |
| samtools index | 1.15 | | |
| YaHS | 1.2a.2 | | |
| Juicer tools | 1.22.01 | | |
| Juicebox | 1.11.08 | | |
| | Gap closing | | |
| TGS-gapcloser | 1.2.1 | --ne --tgstype pb --minimap_arg '-x map-hifi' | |
