## Supplementary material for "From macro to micro: De novo genomes of Aedes mosquitoes enable comparative genomics among close and distant relatives": Table S2

Supplementary table S2: BAC region and arms. NCBI accession IDs for bacterial artificial chromosomes (BAC) used to identify positions along each chromosome. Positional information comes from Matthews et al., (2017).

| BAC | region | arm |
|---|---|---|
| CC138747 | 1p34 | 1p |
| CC858654 | 1p33 | 1p |
| CC151019 | 1p32 | 1p |
| CC117157 | 1p31 | 1p |
| CC866872 | 1p25 | 1p |
| CC139319 | 1p24 | 1p |
| CC848005 | 1p23 | 1p |
| CC844306 | 1p22 | 1p |
| CC866118 | 1p21 | 1p |
| CC125499 | 1p14 | 1p |
| CC112157 | 1p13 | 1p |
| CC857841 | 1p12 | 1p |
| CC864441 | 1q11 | 1q |
| CC867763 | 1q12 | 1q |
| CC118334 | 1q13 | 1q |
| CC846091 | 1q14 | 1q |
| CC845923 | 1q21 | 1q |
| CC121404 | 1q22 | 1q |
| CC122455 | 1q31 | 1q |
| CC120911 | 1q32 | 1q |
| CC114409 | 1q33 | 1q |
| CC856659 | 1q41 | 1q |
| CC143747 | 1q42 | 1q |
| CC134552 | 1q43 | 1q |
| CC853005 | 2p44 | 2p |
| CC869613 | 2p43 | 2p |
| CC112565 | 2p42 | 2p |
| CC109310 | 2p41 | 2p |
| CC845577 | 2p34 | 2p |
| CC134315 | 2p33 | 2p |
| CC858708 | 2p32 | 2p |
| CC859287 | 2p31 | 2p |
| CC119500 | 2p27 | 2p |
| CC130380 | 2p26 | 2p |
| CC846525 | 2p25 | 2p |
| CC135433 | 2p24 | 2p |
| CC842466 | 2p23 | 2p |
| CC122253 | 2p22 | 2p |
| CC123263 | 2p21 | 2p |
| CC115469 | 2p13 | 2p |
| CC140594 | 2p12 | 2p |
| CC126753 | 2q11 | 2q |
| CC860163 | 2q12 | 2q |
| CC843415 | 2q13 | 2q |
| CC125439 | 2q21 | 2q |
| CC871173 | 2q22 | 2q |
| CC852408 | 2q23 | 2q |
| CC119901 | 2q24 | 2q |

| | | |
|---|---|---|
| CC861676 | 2q31 | 2q |
| CC864156 | 2q32 | 2q |
| CC868859 | 2q33 | 2q |
| CC121699 | 2q34 | 2q |
| CC133314 | 2q35 | 2q |
| CC137303 | 2q36 | 2q |
| CC860273 | 2q37 | 2q |
| CC117851 | 2q41 | 2q |
| CC142176 | 2q42 | 2q |
| CC844953 | 2q43 | 2q |
| CC847484 | 3p44 | 3p |
| CC847550 | 3p43 | 3p |
| CC858180 | 3p42 | 3p |
| CC869102 | 3p41 | 3p |
| CC848236 | 3p34 | 3p |
| CC114739 | 3p33 | 3p |
| CC143387 | 3p32 | 3p |
| CC132563 | 3p31 | 3p |
| CC863970 | 3p23 | 3p |
| CC116852 | 3p22 | 3p |
| CC137246 | 3p21 | 3p |
| CC855081 | 3p14 | 3p |
| CC855641 | 3p13 | 3p |
| CC844150 | 3p12 | 3p |
| CC129101 | 3q11 | 3q |
| CC139997 | 3q12 | 3q |
| CC859791 | 3q13 | 3q |
| CC123692 | 3q14 | 3q |
| CC862836 | 3q21 | 3q |
| CC870767 | 3q22 | 3q |
| CC133183 | 3q23 | 3q |
| CC129236 | 3q24 | 3q |
| CC140983 | 3q25 | 3q |
| CC871489 | 3q31 | 3q |
| CC117052 | 3q32 | 3q |
| CC133051 | 3q33 | 3q |
| CC865974 | 3q34 | 3q |
| CC116137 | 3q41 | 3q |
| CC864644 | 3q42 | 3q |
| CC867094 | 3q43 | 3q |
