## Supplementary material for "From macro to micro: De novo genomes of Aedes mosquitoes enable comparative genomics among close and distant relatives": Table S3

**Supplementary Table S3**. Reference assemblies for different Culicidae species.

| Taxon | Assembly Name | NCBI RefSeq Assembly ID |
|---|---|---|
| *Aedes aegypti* | AaegL5.0 | GCF_002204515.2 |
| *Anopheles cruzii* | idAnoCruzAS_RS32_06 | GCF_943734635.1 |
| *Anopheles darlingi* | idANoDarIMG_H_01 | GCF_943734745.1 |
| *Anopheles gambiae* | idAnoGambNW_F1_1 | GCF_943734735.2 |
| *Anopheles ziemanni* | idAnoZiCoDA_A2_x.2 | GCF_943734765.1 |
| *Armigeres subalbatus* | GZ_Asu_2 | GCF_024139115.2 |
| *Sabethes cyaneus* | idSabCyanKW18_F2 | GCF_943734655.1 |
| *Culex pipiens pallens* | TS_CPP_V2 | GCF_016801865.2 |
| *Culex quinquefasciatus* | VPISU_Cqui_1.0_pri_paternal | GCF_015732765.1 |
| *Phlebotomus papatasi* | Ppap_2.1 | GCF_024763615.1 |
