## Supplementary material for "From macro to micro: De novo genomes of Aedes mosquitoes enable comparative genomics among close and distant relatives": Table S4

**Supplementary Table S4**. Metrics for each draft assembly output from Inspector and BUSCO. Collapses, expansions, and inversions are shown errors detected after mapping reads back to the draft assemblies (all measured in bp). Here, 'Sp' is species, 'Asm' is assembler used to assemble reads, and 'Purge' is the purging program used to remove duplicate and haplotig contigs (pd: purge_dups; ph: purge_haplotigs; none: no external purging; internal: purging done internally by assembler). The *Aedes aegypti* reference (AaegL5), has a genome size of approximately 1.3 Gbp.

| | | | Inspector | | | | | | BUSCO (diptera_ODB10; n = 3285) | | | | |
|---|---|---|---|---|---|---|---|---|---|---|---|---|---|
| Sp | Asm | Purge | #contigs | Total length (Gbp) | N50 (Mbp) | Collapses | Expansions | Inversions | C % | S % | D % | F % | M % |
| Aaf | HiCanu | none | 3487 | 2.388 | 2.993 | 159 | 242 | 0 | 97.7 | 5.7 | 91.9 | 1 | 1.3 |
| Aaf | HiCanu | pd | 929 | 1.251 | 4.140 | 22655 | 119094 | 77343 | 96.6 | 91 | 5.5 | 1.5 | 1.9 |
| Aaf | HiCanu | ph | 969 | 1.274 | 4.250 | 8314 | 116510 | 58036 | 96.6 | 89.4 | 7.2 | 1.4 | 2 |
| Aaf | flye | none | 5016 | 2.310 | 1.640 | 202 | 439 | 0 | 97.4 | 10.8 | 86.5 | 1.3 | 1.3 |
| Aaf | flye | pd | 2075 | 1.233 | 2.021 | 4263 | 96176 | 8774 | 96 | 91.3 | 4.7 | 1.8 | 2.2 |
| Aaf | flye | ph | 2005 | 1.249 | 2.045 | 4825 | 101749 | 8774 | 96.5 | 91.2 | 5.2 | 1.8 | 1.7 |
| Aaf | hifiasm | internal | 3631 | 1.799 | 5.341 | 4103 | 111250 | 346539 | 97.6 | 56.9 | 40.7 | 1.1 | 1.3 |
| Aaf | hifiasm | pd | 2403 | 1.292 | 7.526 | 3964 | 96602 | 462957 | 96.2 | 88.5 | 7.7 | 1.4 | 2.4 |
| Aaf | hifiasm | ph | 1478 | 1.370 | 6.930 | 18497 | 325665 | 409347 | 96.7 | 85 | 11.6 | 1.5 | 1.9 |
| Aaf | IPA | internal | 4360 | 2.062 | 2.616 | 108173 | 71430 | 2299 | 96.6 | 92.2 | 4.4 | 1.6 | 1.8 |
| Aaf | IPA | pd | 2645 | 1.378 | 3.131 | 84899 | 97371 | 557616 | 96.1 | 93.9 | 2.2 | 1.6 | 2.3 |
| Aaf | IPA | ph | 2257 | 1.273 | 3.242 | 77172 | 94540 | 554975 | 96.2 | 93.5 | 2.7 | 1.7 | 2.1 |
| Am | HiCanu | none | 2614 | 2.499 | 4.674 | 322 | 107 | 0 | 97.5 | 3.9 | 93.6 | 1.1 | 1.4 |
| Am | HiCanu | pd | 520 | 1.391 | 5.925 | 3481 | 88628 | 132519 | 96.8 | 83 | 13.8 | 1.3 | 1.9 |
| Am | HiCanu | ph | 492 | 1.320 | 6.000 | 3557 | 80161 | 133933 | 96.7 | 88.7 | 8 | 1.4 | 1.9 |
| Am | flye | none | 2818 | 2.461 | 2.125 | 2268 | 3108 | 0 | 97.3 | 5.8 | 91.5 | 1.2 | 1.6 |
| Am | flye | pd | 1162 | 1.391 | 2.884 | 3392 | 77980 | 13457 | 96 | 82.5 | 13.5 | 1.6 | 2.4 |
| Am | flye | ph | 1083 | 1.310 | 2.913 | 3307 | 65708 | 13582 | 96.2 | 89.1 | 7.1 | 1.6 | 2.2 |
| Am | hifiasm | internal | 466 | 1.330 | 14.034 | 4097 | 71041 | 264190 | 97 | 89.3 | 7.7 | 1.3 | 1.7 |
| Am | hifiasm | pd | 180 | 0.747 | 11.268 | 1416 | 31750 | 0 | 58.8 | 57.5 | 1.4 | 1.2 | 39.9 |
| Am | hifiasm | ph | 221 | 1.307 | 14.047 | 4362 | 65393 | 364438 | 96.9 | 90.6 | 6.2 | 1.3 | 1.9 |
| Am | IPA | internal | 906 | 1.904 | 4.215 | 132426 | 65145 | 27696 | 97 | 42.6 | 54.4 | 1.2 | 1.7 |
| Am | IPA | pd | 489 | 1.303 | 5.202 | 46231 | 62975 | 337574 | 94.4 | 86.3 | 8.2 | 1.4 | 4.2 |
| Am | IPA | ph | 452 | 1.308 | 5.453 | 50462 | 84835 | 327316 | 94 | 85.2 | 8.7 | 1.3 | 4.7 |
