## Supplementary material for "From macro to micro: De novo genomes of Aedes mosquitoes enable comparative genomics among close and distant relatives": Table S5

Supplementary table S5. Genomic coordinates and lengths (in bp) of inversions detected in *Aedes aegypti formosus* (Aaf) relative to the *Aedes aegypti* reference genome (AaegL5) from Syri.

| Invesion | AaegL5_seq | Aaf_seq | inv_length | AaegL5_start | AaegL5_end | Aaf_start | Aaf_end |
|---|---|---|---|---|---|---|---|
| INV1430 | AaegL5_1 | scaffold_3 | 16265696 | 1319181 | 13241301 | 171941 | 16437637 |
| INV1431 | AaegL5_1 | scaffold_3 | 520 | 30758215 | 30758740 | 30659028 | 30658508 |
| INV1432 | AaegL5_1 | scaffold_3 | 263199 | 49000124 | 49269139 | 48794185 | 49057384 |
| INV1433 | AaegL5_1 | scaffold_3 | 408 | 51002920 | 51003329 | 50481977 | 50481569 |
| INV1434 | AaegL5_1 | scaffold_3 | 515 | 60769790 | 60770318 | 60208163 | 60207648 |
| INV1435 | AaegL5_1 | scaffold_3 | 451 | 70268758 | 70269209 | 69365235 | 69364784 |
| INV1436 | AaegL5_1 | scaffold_3 | 261 | 78483330 | 78483591 | 77390550 | 77390289 |
| INV1437 | AaegL5_1 | scaffold_3 | 541 | 102768900 | 102769440 | 101220880 | 101220339 |
| INV1438 | AaegL5_1 | scaffold_3 | 332 | 109423624 | 109423955 | 107781821 | 107781489 |
| INV1439 | AaegL5_1 | scaffold_3 | 1640 | 121152885 | 121154528 | 119579727 | 119578087 |
| INV1440 | AaegL5_1 | scaffold_3 | 21235 | 132481439 | 132502669 | 130788493 | 130767258 |
| INV1441 | AaegL5_1 | scaffold_3 | 467 | 135221826 | 135222290 | 133409280 | 133408813 |
| INV1442 | AaegL5_1 | scaffold_3 | 659 | 139005751 | 139006410 | 137154218 | 137153559 |
| INV1443 | AaegL5_1 | scaffold_3 | 483 | 140281304 | 140281788 | 138467871 | 138467388 |
| INV1444 | AaegL5_1 | scaffold_3 | 2573 | 153307612 | 153309588 | 151434711 | 151432138 |
| INV1445 | AaegL5_1 | scaffold_3 | 22033 | 154063033 | 154087002 | 152860801 | 152838768 |
| INV1446 | AaegL5_1 | scaffold_3 | 463 | 154853235 | 154853698 | 153536206 | 153535743 |
| INV1447 | AaegL5_1 | scaffold_3 | 343 | 155500544 | 155500887 | 154188770 | 154188427 |
| INV1448 | AaegL5_1 | scaffold_3 | 414 | 161039607 | 161040021 | 159523682 | 159523268 |
| INV1449 | AaegL5_1 | scaffold_3 | 410 | 161587116 | 161587526 | 160137140 | 160136730 |
| INV1450 | AaegL5_1 | scaffold_3 | 1030 | 165194161 | 165195190 | 163550034 | 163549004 |
| INV1451 | AaegL5_1 | scaffold_3 | 276 | 165654300 | 165654576 | 164119149 | 164118873 |
| INV1452 | AaegL5_1 | scaffold_3 | 494 | 173075674 | 173076166 | 171510941 | 171510447 |
| INV1453 | AaegL5_1 | scaffold_3 | 264 | 173771928 | 173772192 | 172211010 | 172210746 |
| INV1454 | AaegL5_1 | scaffold_3 | 1306 | 180139263 | 180140570 | 178273676 | 178272370 |
| INV1455 | AaegL5_1 | scaffold_3 | 407 | 191411264 | 191411671 | 189437923 | 189437516 |
| INV1456 | AaegL5_1 | scaffold_3 | 1654 | 191591868 | 191593527 | 189591743 | 189590089 |
| INV1457 | AaegL5_1 | scaffold_3 | 818056 | 193151103 | 193912430 | 191523686 | 192341742 |
| INV1458 | AaegL5_1 | scaffold_3 | 277 | 196721868 | 196722145 | 194991529 | 194991252 |
| INV1459 | AaegL5_1 | scaffold_3 | 925 | 205989836 | 205990755 | 204182002 | 204181077 |
| INV1460 | AaegL5_1 | scaffold_3 | 232 | 206669328 | 206669560 | 204872780 | 204872548 |
| INV1461 | AaegL5_1 | scaffold_3 | 402 | 225601617 | 225602018 | 223016500 | 223016098 |
| INV1462 | AaegL5_1 | scaffold_3 | 299 | 226574346 | 226574645 | 223860357 | 223860058 |
| INV1463 | AaegL5_1 | scaffold_3 | 1425 | 244024003 | 244025416 | 239972689 | 239971264 |
| INV1464 | AaegL5_1 | scaffold_3 | 223 | 248661175 | 248661398 | 244329899 | 244329676 |
| INV1465 | AaegL5_1 | scaffold_3 | 323 | 255208037 | 255208360 | 250837344 | 250837021 |
| INV1466 | AaegL5_1 | scaffold_3 | 18387412 | 271773706 | 286753394 | 267885290 | 286272702 |
| INV1467 | AaegL5_1 | scaffold_3 | 674 | 288024644 | 288025318 | 287752858 | 287752184 |
| INV1468 | AaegL5_1 | scaffold_3 | 15337 | 289919487 | 289937265 | 289685818 | 289701155 |
| INV1469 | AaegL5_1 | scaffold_3 | 3414 | 290900067 | 290903478 | 290564970 | 290561556 |
| INV1470 | AaegL5_1 | scaffold_3 | 366613 | 294448714 | 294861650 | 294444722 | 294811335 |
| INV1471 | AaegL5_1 | scaffold_3 | 3716 | 297056322 | 297060027 | 296993361 | 296989645 |
| INV1472 | AaegL5_1 | scaffold_3 | 431276 | 300137001 | 300462239 | 301101580 | 301532856 |
| INV1473 | AaegL5_1 | scaffold_3 | 936 | 304243775 | 304244715 | 305441035 | 305440099 |
| INV1474 | AaegL5_1 | scaffold_3 | 425867 | 307675523 | 308085041 | 310205592 | 310631459 |
| INV1475 | AaegL5_2 | scaffold_1 | 1113 | 609603 | 610717 | 621935 | 620822 |
| INV1476 | AaegL5_2 | scaffold_1 | 584155 | 8884413 | 9410554 | 9264153 | 9848308 |
| INV1477 | AaegL5_2 | scaffold_1 | 696 | 21059688 | 21060379 | 22012401 | 22011705 |
| INV1478 | AaegL5_2 | scaffold_1 | 360 | 30294940 | 30295300 | 31804160 | 31803800 |
| INV1479 | AaegL5_2 | scaffold_1 | 263 | 37869698 | 37869961 | 39479220 | 39478957 |
| INV1480 | AaegL5_2 | scaffold_1 | 353 | 38843977 | 38844330 | 40427640 | 40427287 |
| INV1481 | AaegL5_2 | scaffold_1 | 349 | 39594233 | 39594582 | 41189194 | 41188845 |
| INV1482 | AaegL5_2 | scaffold_1 | 718 | 45792527 | 45793248 | 47513939 | 47513221 |
| INV1483 | AaegL5_2 | scaffold_1 | 511 | 46367111 | 46367622 | 48104032 | 48103521 |

| | | | | | | | |
|---|---|---|---|---|---|---|---|
| INV1484 | AaegL5_2 | scaffold_1 | 731 | 48410261 | 48410995 | 49980591 | 49979860 |
| INV1485 | AaegL5_2 | scaffold_1 | 1137 | 51031777 | 51032899 | 52452844 | 52451707 |
| INV1486 | AaegL5_2 | scaffold_1 | 355 | 51282025 | 51282380 | 52673711 | 52673356 |
| INV1487 | AaegL5_2 | scaffold_1 | 632 | 68960429 | 68961063 | 69898861 | 69898229 |
| INV1488 | AaegL5_2 | scaffold_1 | 690 | 71060851 | 71061549 | 71967928 | 71967238 |
| INV1489 | AaegL5_2 | scaffold_1 | 283 | 92047095 | 92047378 | 91593411 | 91593128 |
| INV1490 | AaegL5_2 | scaffold_1 | 644 | 96365039 | 96365683 | 95521675 | 95521031 |
| INV1491 | AaegL5_2 | scaffold_1 | 569 | 111591348 | 111591918 | 109243748 | 109243179 |
| INV1492 | AaegL5_2 | scaffold_1 | 209307 | 113772656 | 113991201 | 111153595 | 111362902 |
| INV1493 | AaegL5_2 | scaffold_1 | 931 | 120977901 | 120978830 | 118235584 | 118234653 |
| INV1494 | AaegL5_2 | scaffold_1 | 495 | 132581692 | 132582187 | 129053744 | 129053249 |
| INV1495 | AaegL5_2 | scaffold_1 | 728 | 132662470 | 132663196 | 129120028 | 129119300 |
| INV1496 | AaegL5_2 | scaffold_1 | 1216 | 133410827 | 133412047 | 129783606 | 129782390 |
| INV1497 | AaegL5_2 | scaffold_1 | 1280 | 144850300 | 144851574 | 140936999 | 140935719 |
| INV1498 | AaegL5_2 | scaffold_1 | 375 | 147294864 | 147295239 | 143303916 | 143303541 |
| INV1499 | AaegL5_2 | scaffold_1 | 459 | 153366504 | 153366963 | 148939824 | 148939365 |
| INV1500 | AaegL5_2 | scaffold_1 | 1202 | 166830922 | 166832141 | 162170629 | 162169427 |
| INV1501 | AaegL5_2 | scaffold_1 | 334 | 167344858 | 167345192 | 162734523 | 162734189 |
| INV1502 | AaegL5_2 | scaffold_1 | 373 | 169727526 | 169727899 | 165137567 | 165137194 |
| INV1503 | AaegL5_2 | scaffold_1 | 373 | 172628373 | 172628746 | 167903461 | 167903088 |
| INV1504 | AaegL5_2 | scaffold_1 | 2365 | 178836265 | 178838555 | 173489935 | 173487570 |
| INV1505 | AaegL5_2 | scaffold_1 | 501 | 191929888 | 191930389 | 186148623 | 186148122 |
| INV1506 | AaegL5_2 | scaffold_1 | 967 | 195939359 | 195940316 | 190053571 | 190052604 |
| INV1507 | AaegL5_2 | scaffold_1 | 637762 | 198046260 | 198723282 | 192455915 | 193093677 |
| INV1508 | AaegL5_2 | scaffold_1 | 374 | 201002534 | 201002908 | 195522252 | 195521878 |
| INV1509 | AaegL5_2 | scaffold_1 | 1319 | 203686317 | 203687624 | 198068960 | 198067641 |
| INV1510 | AaegL5_2 | scaffold_1 | 321 | 203747329 | 203747651 | 198136409 | 198136088 |
| INV1511 | AaegL5_2 | scaffold_1 | 225 | 208799007 | 208799232 | 202971912 | 202971687 |
| INV1512 | AaegL5_2 | scaffold_1 | 727 | 210510979 | 210511717 | 204699798 | 204699071 |
| INV1513 | AaegL5_2 | scaffold_1 | 1670 | 212323668 | 212325335 | 206554064 | 206552394 |
| INV1514 | AaegL5_2 | scaffold_1 | 830 | 214188812 | 214189654 | 208335713 | 208334883 |
| INV1515 | AaegL5_2 | scaffold_1 | 652 | 219162487 | 219163139 | 213194434 | 213193782 |
| INV1516 | AaegL5_2 | scaffold_1 | 230 | 224373142 | 224373372 | 217712085 | 217711855 |
| INV1517 | AaegL5_2 | scaffold_1 | 228 | 226685067 | 226685295 | 219939826 | 219939598 |
| INV1518 | AaegL5_2 | scaffold_1 | 488 | 228146406 | 228146892 | 221154480 | 221153992 |
| INV1519 | AaegL5_2 | scaffold_1 | 853 | 228603188 | 228604041 | 221745472 | 221744619 |
| INV1520 | AaegL5_2 | scaffold_1 | 2480 | 229619048 | 229621528 | 223017166 | 223014686 |
| INV1521 | AaegL5_2 | scaffold_1 | 417 | 231672562 | 231672979 | 225021013 | 225020596 |
| INV1522 | AaegL5_2 | scaffold_1 | 318 | 232311552 | 232311870 | 225701830 | 225701512 |
| INV1523 | AaegL5_2 | scaffold_1 | 691 | 248621464 | 248622155 | 243565623 | 243564932 |
| INV1524 | AaegL5_2 | scaffold_1 | 719 | 268612523 | 268613260 | 263106633 | 263105914 |
| INV1525 | AaegL5_2 | scaffold_1 | 328 | 268675446 | 268675774 | 263174553 | 263174225 |
| INV1526 | AaegL5_2 | scaffold_1 | 933 | 272015248 | 272016170 | 266397184 | 266396251 |
| INV1527 | AaegL5_2 | scaffold_1 | 1995 | 273678242 | 273680243 | 267976583 | 267974588 |
| INV1528 | AaegL5_2 | scaffold_1 | 562 | 276339468 | 276340039 | 270634006 | 270633444 |
| INV1529 | AaegL5_2 | scaffold_1 | 378 | 293261212 | 293261590 | 287454606 | 287454228 |
| INV1530 | AaegL5_2 | scaffold_1 | 247 | 305570746 | 305570993 | 299212935 | 299212688 |
| INV1531 | AaegL5_2 | scaffold_1 | 450 | 311817822 | 311818272 | 305124860 | 305124410 |
| INV1532 | AaegL5_2 | scaffold_1 | 10364 | 312719677 | 312834456 | 306039726 | 306050090 |
| INV1533 | AaegL5_2 | scaffold_1 | 768 | 314016461 | 314017229 | 307126614 | 307125846 |
| INV1534 | AaegL5_2 | scaffold_1 | 484 | 326296081 | 326296565 | 319454423 | 319453939 |
| INV1535 | AaegL5_2 | scaffold_1 | 436 | 331131036 | 331131472 | 323912483 | 323912047 |
| INV1536 | AaegL5_2 | scaffold_1 | 1617 | 335814901 | 335816531 | 328198557 | 328196940 |
| INV1537 | AaegL5_2 | scaffold_1 | 328 | 396635242 | 396635570 | 385708586 | 385708258 |
| INV1538 | AaegL5_2 | scaffold_1 | 857 | 399393347 | 399394218 | 388284120 | 388283263 |
| INV1539 | AaegL5_2 | scaffold_1 | 453 | 439379277 | 439379727 | 426901201 | 426900748 |
| INV1540 | AaegL5_2 | scaffold_1 | 452 | 442088790 | 442089243 | 429608463 | 429608011 |
| INV1541 | AaegL5_2 | scaffold_1 | 322 | 460609941 | 460610263 | 448338515 | 448338193 |
| INV1542 | AaegL5_2 | scaffold_1 | 761 | 472826121 | 472826893 | 461149786 | 461149025 |

| | | | | | | | |
|---|---|---|---|---|---|---|---|
| INV1543 | AaegL5_3 | scaffold_2 | 1737777 | 10074843 | 11830203 | 11201268 | 12939045 |
| INV1544 | AaegL5_3 | scaffold_2 | 578 | 25689814 | 25690387 | 27465417 | 27464839 |
| INV1545 | AaegL5_3 | scaffold_2 | 1021 | 30933042 | 30934051 | 32371785 | 32370764 |
| INV1546 | AaegL5_3 | scaffold_2 | 44961 | 36135432 | 36182472 | 36188156 | 36233117 |
| INV1547 | AaegL5_3 | scaffold_2 | 331 | 37717295 | 37717626 | 38919959 | 38919628 |
| INV1548 | AaegL5_3 | scaffold_2 | 628 | 41475424 | 41476052 | 42459698 | 42459070 |
| INV1549 | AaegL5_3 | scaffold_2 | 253 | 51166022 | 51166275 | 52055308 | 52055055 |
| INV1550 | AaegL5_3 | scaffold_2 | 317 | 56708746 | 56709063 | 57595090 | 57594773 |
| INV1551 | AaegL5_3 | scaffold_2 | 308 | 57615470 | 57615778 | 58508838 | 58508530 |
| INV1552 | AaegL5_3 | scaffold_2 | 232 | 87855161 | 87855393 | 86886915 | 86886683 |
| INV1553 | AaegL5_3 | scaffold_2 | 755 | 91839726 | 91840482 | 90202544 | 90201789 |
| INV1554 | AaegL5_3 | scaffold_2 | 1055 | 99966772 | 99967827 | 96247219 | 96246164 |
| INV1555 | AaegL5_3 | scaffold_2 | 789 | 105467949 | 105468739 | 101445858 | 101445069 |
| INV1556 | AaegL5_3 | scaffold_2 | 526 | 106695708 | 106696234 | 102564161 | 102563635 |
| INV1557 | AaegL5_3 | scaffold_2 | 258950 | 108589261 | 108867146 | 104250153 | 104509103 |
| INV1558 | AaegL5_3 | scaffold_2 | 831 | 133507484 | 133508315 | 128574623 | 128573792 |
| INV1559 | AaegL5_3 | scaffold_2 | 440 | 134308117 | 134308557 | 129364472 | 129364032 |
| INV1560 | AaegL5_3 | scaffold_2 | 225 | 149562894 | 149563119 | 143557364 | 143557139 |
| INV1561 | AaegL5_3 | scaffold_2 | 939 | 153721248 | 153722189 | 147637746 | 147636807 |
| INV1562 | AaegL5_3 | scaffold_2 | 311 | 156034470 | 156034785 | 150115643 | 150115332 |
| INV1563 | AaegL5_3 | scaffold_2 | 269 | 165497574 | 165497843 | 159684855 | 159684586 |
| INV1564 | AaegL5_3 | scaffold_2 | 823 | 167765472 | 167766295 | 161785236 | 161784413 |
| INV1565 | AaegL5_3 | scaffold_2 | 491 | 183081877 | 183082369 | 176911494 | 176911003 |
| INV1566 | AaegL5_3 | scaffold_2 | 70883 | 185003506 | 185088317 | 178494690 | 178565573 |
| INV1567 | AaegL5_3 | scaffold_2 | 473 | 196960564 | 196961031 | 189878462 | 189877989 |
| INV1568 | AaegL5_3 | scaffold_2 | 570 | 201588267 | 201588837 | 196069884 | 196069314 |
| INV1569 | AaegL5_3 | scaffold_2 | 557 | 201964686 | 201965243 | 196601344 | 196600787 |
| INV1570 | AaegL5_3 | scaffold_2 | 247 | 202634192 | 202634439 | 197183684 | 197183437 |
| INV1571 | AaegL5_3 | scaffold_2 | 399 | 207212898 | 207213298 | 201532979 | 201532580 |
| INV1572 | AaegL5_3 | scaffold_2 | 756 | 207575607 | 207576363 | 201844470 | 201843714 |
| INV1573 | AaegL5_3 | scaffold_2 | 272 | 209442538 | 209442810 | 203571770 | 203571498 |
| INV1574 | AaegL5_3 | scaffold_2 | 279 | 212702637 | 212702916 | 206777755 | 206777476 |
| INV1575 | AaegL5_3 | scaffold_2 | 281 | 218300268 | 218300549 | 212264196 | 212263915 |
| INV1576 | AaegL5_3 | scaffold_2 | 518 | 220085875 | 220086393 | 214004811 | 214004293 |
| INV1577 | AaegL5_3 | scaffold_2 | 262 | 226648539 | 226648801 | 220695235 | 220694973 |
| INV1578 | AaegL5_3 | scaffold_2 | 263 | 226700537 | 226700800 | 220769830 | 220769567 |
| INV1579 | AaegL5_3 | scaffold_2 | 1189 | 240707520 | 240708698 | 234351717 | 234350528 |
| INV1580 | AaegL5_3 | scaffold_2 | 292 | 242492949 | 242493241 | 236017490 | 236017198 |
| INV1581 | AaegL5_3 | scaffold_2 | 305 | 244590513 | 244590818 | 237937460 | 237937155 |
| INV1582 | AaegL5_3 | scaffold_2 | 659 | 251717690 | 251718347 | 245045187 | 245044528 |
| INV1583 | AaegL5_3 | scaffold_2 | 181043 | 257074932 | 257245310 | 250743166 | 250924209 |
| INV1584 | AaegL5_3 | scaffold_2 | 297 | 257864376 | 257864673 | 251625806 | 251625509 |
| INV1585 | AaegL5_3 | scaffold_2 | 1047 | 273475655 | 273476693 | 266775688 | 266774641 |
| INV1586 | AaegL5_3 | scaffold_2 | 493 | 286043225 | 286043718 | 278404096 | 278403603 |
| INV1587 | AaegL5_3 | scaffold_2 | 350 | 294696773 | 294697123 | 286498481 | 286498131 |
| INV1588 | AaegL5_3 | scaffold_2 | 25964 | 305665072 | 305690710 | 296785068 | 296759104 |
| INV1589 | AaegL5_3 | scaffold_2 | 181738 | 311453958 | 311642205 | 302531981 | 302713719 |
| INV1590 | AaegL5_3 | scaffold_2 | 316 | 321937857 | 321938173 | 312541421 | 312541105 |
| INV1591 | AaegL5_3 | scaffold_2 | 685 | 323283762 | 323284455 | 313897117 | 313896432 |
| INV1592 | AaegL5_3 | scaffold_2 | 633 | 324613417 | 324614050 | 315148238 | 315147605 |
| INV1593 | AaegL5_3 | scaffold_2 | 344 | 340234449 | 340234794 | 329924976 | 329924632 |
| INV1594 | AaegL5_3 | scaffold_2 | 36456 | 341243391 | 341425706 | 330733673 | 330770129 |
| INV1595 | AaegL5_3 | scaffold_2 | 6117 | 341439743 | 341445720 | 330802548 | 330796431 |
| INV1596 | AaegL5_3 | scaffold_2 | 41970 | 341675429 | 341792295 | 330858832 | 330900802 |
| INV1597 | AaegL5_3 | scaffold_2 | 13905 | 341793095 | 341830482 | 330920016 | 330933921 |
| INV1598 | AaegL5_3 | scaffold_2 | 253 | 342305550 | 342305803 | 331127720 | 331127467 |
| INV1599 | AaegL5_3 | scaffold_2 | 1122 | 344324195 | 344325317 | 333130461 | 333129339 |
| INV1600 | AaegL5_3 | scaffold_2 | 510 | 360373764 | 360374274 | 348395836 | 348395326 |
| INV1601 | AaegL5_3 | scaffold_2 | 355 | 383295514 | 383295869 | 370990685 | 370990330 |

| | | | | | | | |
|---|---|---|---|---|---|---|---|
| INV1602 | AaegL5_3 | scaffold_2 | 1140263 | 405132013 | 406450969 | 393521328 | 394661591 |
| INV1603 | AaegL5_3 | scaffold_2 | 998914 | 407752933 | 408915508 | 395651191 | 396650105 |
| INV1604 | AaegL5_3 | scaffold_2 | 562761 | 409093692 | 409721457 | 396777357 | 397340118 |
