## Supplementary material for "From macro to micro: De novo genomes of Aedes mosquitoes enable comparative genomics among close and distant relatives": Table S6

Supplementary table S6: overrepresentation test of Aegypti group assemblies in a single analysis. Overrepresentation test performed on the three Aegypti group mosquito assemblies using the PANTHER DB overpresentation test (release 20240807). The reference annotation set is from *Anopheles gambiae*. Significance testing used Fisher's Exact test and the resulting P values were corrected by accounting for false discovery rate (FDR). Here, 'Reference' refers to the *An. gambiae* reference annotation set, 'Aaf' refers to *Aedes aegpyti formosus*, 'Am' refers *Ae. mascarensis*, and 'AaegL5' refers to the *Ae. aegypti* reference genome. 'N' in the column refers to the number of terms observed in each set, 'fold' refers to the fold enrichment. Indentation here reflects the nestedness of each GO term

| GO | GO id | Reference N | Aaf N | Aaf expected N | Aaf fold enrichment | Aaf FDR | AaegL5 N | AaegL5 expected N | AaegL5 fold enrichment | AaegL5 FDR | Am N | Am expected N | Am fold enrichment | Am FDR |
|---|---|---|---|---|---|---|---|---|---|---|---|---|---|---|
| male courtship behavior | GO:0008049 | 1488 | 41 | 3.76 | 10.92 | 1.68E-37 | 41 | 4.23 | 9.7 | 1.53E-35 | 41 | 4.58 | 8.95 | 1.12E-33 |
| male mating behavior | GO:0060179 | 1335 | 41 | 3.76 | 10.92 | 2.16E-37 | 41 | 4.23 | 9.7 | 1.81E-35 | 41 | 4.58 | 8.95 | 1.87E-33 |
| mating behavior | GO:0007617 | 852 | 41 | 3.84 | 10.68 | 1.17E-36 | 41 | 4.32 | 9.49 | 1.11E-34 | 41 | 4.68 | 8.75 | 6.65E-33 |
| reproductive behavior | GO:0019098 | 785 | 41 | 3.84 | 10.68 | 1.28E-36 | 41 | 4.32 | 9.49 | 1.19E-34 | 41 | 4.68 | 8.75 | 7.75E-33 |
| multicellular organismal reproductive process | GO:0048609 | 845 | 73 | 9.52 | 7.67 | 5.66E-47 | 57 | 10.71 | 5.32 | 1.48E-26 | 49 | 11.61 | 4.22 | 3.52E-17 |
| reproductive process | GO:0022414 | 314 | 84 | 14.94 | 5.62 | 1.91E-40 | 67 | 16.81 | 3.99 | 3.00E-22 | 58 | 18.23 | 3.18 | 1.09E-13 |
| behavior | GO:0007610 | 714 | 42 | 5.43 | 7.74 | 2.61E-27 | 44 | 6.1 | 7.21 | 6.28E-28 | 46 | 6.62 | 6.95 | 1.19E-28 |
| multicellular organismal process | GO:0032501 | 63 | 178 | 59.6 | 2.99 | 1.91E-40 | 170 | 67.06 | 2.54 | 2.12E-29 | 114 | 72.71 | 1.57 | 0.000109 |
| courtship behavior | GO:0007619 | 90 | 41 | 3.76 | 10.92 | 1.89E-37 | 41 | 4.23 | 9.7 | 1.66E-35 | 41 | 4.58 | 8.95 | 1.40E-33 |
| chemosensory behavior | GO:0007635 | 274 | 41 | 4.42 | 9.27 | 1.08E-31 | 43 | 4.98 | 8.64 | 7.72E-33 | 45 | 5.4 | 8.34 | 1.06E-33 |
| response to chemical | GO:0042221 | 245 | 59 | 26.21 | 2.25 | 2.95E-07 | 119 | 29.49 | 4.04 | 5.96E-41 | 69 | 31.98 | 2.16 | 1.20E-07 |
| sensory perception of taste | GO:0050909 | 264 | 45 | 5.59 | 8.05 | 1.89E-30 | 46 | 6.29 | 7.31 | 1.23E-29 | 52 | 6.82 | 7.62 | 3.67E-35 |
| sensory perception of chemical stimulus | GO:0007606 | 206 | 82 | 17.19 | 4.77 | 3.04E-33 | 117 | 19.35 | 6.05 | 5.10E-63 | 56 | 20.98 | 2.67 | 1.09E-09 |
| sensory perception | GO:0007600 | 179 | 96 | 20.45 | 4.69 | 2.89E-38 | 118 | 23.01 | 5.13 | 1.05E-53 | 57 | 24.95 | 2.28 | 3.45E-07 |
| nervous system process | GO:0050877 | 23 | 104 | 22.04 | 4.72 | 1.87E-41 | 127 | 24.8 | 5.12 | 1.11E-57 | 64 | 26.88 | 2.38 | 1.08E-08 |
| system process | GO:0003008 | 12 | 106 | 22.87 | 4.63 | 1.33E-41 | 128 | 25.73 | 4.97 | 1.92E-56 | 64 | 27.9 | 2.29 | 4.69E-08 |
| excitatory postsynaptic potential | GO:0060079 | 12 | 7 | 1 | 6.99 | 0.000813 | 7 | 1.13 | 6.21 | 0.00271 | 7 | 1.22 | 5.73 | 0.00476 |
| chemical synaptic transmission, postsynaptic | GO:0099565 | 114 | 7 | 1 | 6.99 | 0.000804 | 7 | 1.13 | 6.21 | 0.00266 | 7 | 1.22 | 5.73 | 0.00468 |
| synaptic signaling | GO:0099536 | 65 | 29 | 7.51 | 3.86 | 1.31E-08 | 20 | 8.45 | 2.37 | 0.0183 | 20 | 9.17 | 2.18 | 0.0378 |
| signaling | GO:0023052 | 67 | 134 | 70.53 | 1.9 | 3.96E-11 | 120 | 79.36 | 1.51 | 0.000352 | 126 | 86.05 | 1.46 | 0.000949 |
| cell communication | GO:0007154 | 53 | 134 | 71.11 | 1.88 | 5.15E-11 | 121 | 80.02 | 1.51 | 0.000295 | 126 | 86.77 | 1.45 | 0.00132 |
| signal transduction | GO:0007165 | 46 | 124 | 65.52 | 1.89 | 2.90E-10 | 110 | 73.73 | 1.49 | 0.0015 | 115 | 79.94 | 1.44 | 0.00474 |
| regulation of postsynaptic membrane potential | GO:0060078 | 46 | 11 | 1.92 | 5.73 | 4.82E-05 | 11 | 2.16 | 5.09 | 0.000263 | 11 | 2.34 | 4.7 | 0.00057 |
| regulation of membrane potential | GO:0042391 | 45 | 20 | 5.26 | 3.8 | 7.39E-06 | 18 | 5.92 | 3.04 | 0.00117 | 17 | 6.42 | 2.65 | 0.0101 |
| cellular component organization | GO:0016043 | 45 | 68 | 111.43 | 0.61 | 0.000117 | 93 | 125.38 | 0.74 | 0.0409 | 90 | 135.95 | 0.66 | 0.000594 |
| cellular component organization or biogenesis | GO:0071840 | 45 | 73 | 124.2 | 0.59 | 5.34E-06 | 101 | 139.75 | 0.72 | 0.0112 | 108 | 151.53 | 0.71 | 0.00429 |
