## Supplementary material for "From macro to micro: De novo genomes of Aedes mosquitoes enable comparative genomics among close and distant relatives": Table S7

Supplementary Table S7: All genes in inversions. List of genes, their function assigned by PANTHER DB, and their locations on the chromosomes of Aaf. We used the descriptive names assigned to AaegL5 to subset this list for the selected gene list

| Sequence name | Start | End | Aaf name | AaegL5 name | PATHER ID | PANTHER Family | PANTHER GO |
|---|---|---|---|---|---|---|---|
| chr3 | 330742477 | 330743157 | Aedes_aegypti_formosus\|g8279.t1 | XP_021712839.1 histone H1-like [Aedes aegypti] | PTHR11467 | HISTONE H1 | nucleosome binding#GO:0031491;double-stranded DNA binding#GO:0003690;chromatin DNA binding#GO:0031490;organic cyclic compound binding#GO:0097159;DNA binding#GO:0003677 |
| chr3 | 330758342 | 330759022 | Aedes_aegypti_formosus\|g8280.t1 | XP_021712839.1 histone H1-like [Aedes aegypti] | PTHR11467 | HISTONE H1 | nucleosome binding#GO:0031491;double-stranded DNA binding#GO:0003690;chromatin DNA binding#GO:0031490;organic cyclic compound binding#GO:0097159;DNA binding#GO:0003677 |
| chr1 | 14251308 | 14258651 | Aedes_aegypti_formosus\|g10879.t1 | XP_021707229.1 uncharacterized protein LOC110678537 isoform X2 [Aedes aegypti] | PTHR47027 | REVERSE TRANSCRIPTASE DOMAIN-CONTAINING PROTEIN | |
| chr3 | 330874476 | 330874850 | Aedes_aegypti_formosus\|g8292.t1 | XP_021712017.1 histone H2A-like [Aedes aegypti] | PTHR23430 | HISTONE H2A | heterocyclic compound binding#GO:1901363;organic cyclic compound binding#GO:0097159;DNA binding#GO:0003677 |
| chr1 | 5458791 | 5462399 | Aedes_aegypti_formosus\|g10682.t1 | XP_021709602.1 uncharacterized protein LOC110679408 [Aedes aegypti] | PTHR19303 | TRANSPOSON | |
| chr1 | 14144267 | 14145397 | Aedes_aegypti_formosus\|g10872.t1 | XP_001655176.1 actin-5C [Aedes aegypti] | PTHR11937 | ACTIN | |
| chr1 | 277085843 | 277087753 | Aedes_aegypti_formosus\|g16400.t1 | XP_021693655.1 heat shock protein 70 A1-like [Aedes aegypti] | PTHR19375 | HEAT SHOCK PROTEIN 70KDA | |
| chr1 | 277088834 | 277090750 | Aedes_aegypti_formosus\|g16401.t1 | XP_021693654.1 heat shock protein 70 A1-like [Aedes aegypti] | PTHR19375 | HEAT SHOCK PROTEIN 70KDA | |
| chr1 | 277110420 | 277112336 | Aedes_aegypti_formosus\|g16402.t1 | XP_021693654.1 heat shock protein 70 A1-like [Aedes aegypti] | PTHR19375 | HEAT SHOCK PROTEIN 70KDA | |
| chr1 | 277117429 | 277119345 | Aedes_aegypti_formosus\|g16403.t1 | XP_021693649.1 heat shock protein 70 A1 [Aedes aegypti] | PTHR19375 | HEAT SHOCK PROTEIN 70KDA | |
| chr1 | 277133127 | 277135043 | Aedes_aegypti_formosus\|g16404.t1 | XP_021693649.1 heat shock protein 70 A1 [Aedes aegypti] | PTHR19375 | HEAT SHOCK PROTEIN 70KDA | |
| chr1 | 277138098 | 277140014 | Aedes_aegypti_formosus\|g16405.t1 | XP_021693649.1 heat shock protein 70 A1 [Aedes aegypti] | PTHR19375 | HEAT SHOCK PROTEIN 70KDA | |
| chr1 | 280860049 | 280860648 | Aedes_aegypti_formosus\|g16468.t1 | XP_001660359.1 cuticle protein [Aedes aegypti] | PTHR12236 | STRUCTURAL CONTITUENT OF CUTICLE | |
| chr1 | 280894255 | 280895060 | Aedes_aegypti_formosus\|g16469.t1 | XP_001657546.2 larval cuticle protein A3A [Aedes aegypti] | PTHR12236 | STRUCTURAL CONTITUENT OF CUTICLE | |
| chr1 | 280895793 | 280896569 | Aedes_aegypti_formosus\|g16470.t1 | XP_001660358.1 larval cuticle protein A3A [Aedes aegypti] | PTHR12236 | STRUCTURAL CONTITUENT OF CUTICLE | |
| chr1 | 280900225 | 280901024 | Aedes_aegypti_formosus\|g16471.t1 | XP_001660357.1 larval cuticle protein A3A [Aedes aegypti] | PTHR12236 | STRUCTURAL CONTITUENT OF CUTICLE | |
| chr1 | 280906224 | 280907034 | Aedes_aegypti_formosus\|g16472.t1 | XP_001660356.1 cuticle protein [Aedes aegypti] | PTHR12236 | STRUCTURAL CONTITUENT OF CUTICLE | |
| chr1 | 280909721 | 280910628 | Aedes_aegypti_formosus\|g16473.t1 | XP_001660355.2 cuticle protein [Aedes aegypti] | PTHR12236 | STRUCTURAL CONTITUENT OF CUTICLE | |
| chr1 | 280911603 | 280912401 | Aedes_aegypti_formosus\|g16474.t1 | XP_001660354.2 cuticle protein [Aedes aegypti] | PTHR12236 | STRUCTURAL CONTITUENT OF CUTICLE | |
| chr1 | 280917897 | 280918582 | Aedes_aegypti_formosus\|g16475.t1 | XP_021693581.1 cuticle protein isoform X4 [Aedes aegypti] | PTHR12236 | STRUCTURAL CONTITUENT OF CUTICLE | |
| chr1 | 280925339 | 280926398 | Aedes_aegypti_formosus\|g16476.t1 | XP_021693570.1 cuticle protein-like isoform X2 [Aedes aegypti] | PTHR12236 | STRUCTURAL CONTITUENT OF CUTICLE | |
| chr1 | 280933315 | 280934000 | Aedes_aegypti_formosus\|g16477.t1 | XP_021693573.1 cuticle protein isoform X3 [Aedes aegypti] | PTHR12236 | STRUCTURAL CONTITUENT OF CUTICLE | |
| chr1 | 280938751 | 280939435 | Aedes_aegypti_formosus\|g16478.t1 | XP_021693573.1 cuticle protein isoform X3 [Aedes aegypti] | PTHR12236 | STRUCTURAL CONTITUENT OF CUTICLE | |
| chr1 | 280945839 | 280946531 | Aedes_aegypti_formosus\|g16479.t1 | XP_001660349.1 larval cuticle protein A2B [Aedes aegypti] | PTHR12236 | STRUCTURAL CONTITUENT OF CUTICLE | |
| chr1 | 280955374 | 280956064 | Aedes_aegypti_formosus\|g16480.t1 | XP_021693574.1 cuticle protein [Aedes aegypti] | PTHR12236 | STRUCTURAL CONTITUENT OF CUTICLE | |
| chr1 | 6413708 | 6414280 | Aedes_aegypti_formosus\|g10704.t1 | XP_021699258.1 uncharacterized protein LOC110676282 [Aedes aegypti] | PTHR46564 | TRANSPOSASE | |
| chr1 | 5130872 | 5136489 | Aedes_aegypti_formosus\|g10672.t1 | XP_021694449.1 tubulin alpha-1A chain [Aedes aegypti] | PTHR11588 | TUBULIN | ribonucleotide binding#GO:0032553;structural molecule activity#GO:0005198;organic cyclic compound binding#GO:0097159;anion binding#GO:0043168;GTP binding#GO:0005525;structural constituent of cytoskeleton#GO:0005200;guanyl ribonucleotide binding#GO:0032561;purine ribonucleoside triphosphate binding#GO:0035639 |
| chr1 | 269483497 | 269491536 | Aedes_aegypti_formosus\|g16239.t1 | XP_011493212.2 presenilin homolog isoform X2 [Aedes aegypti] | PTHR19446 | REVERSE TRANSCRIPTASES | |
| chr1 | 294548860 | 294566407 | Aedes_aegypti_formosus\|g16755.t1 | XP_001650331.1 tubulin beta chain [Aedes aegypti] | PTHR11588 | TUBULIN | ribonucleotide binding#GO:0032553;structural molecule activity#GO:0005198;organic cyclic compound binding#GO:0097159;anion binding#GO:0043168;GTP binding#GO:0005525;structural constituent of cytoskeleton#GO:0005200;guanyl ribonucleotide binding#GO:0032561;purine ribonucleoside triphosphate binding#GO:0035639 |
| chr1 | 294582455 | 294609542 | Aedes_aegypti_formosus\|g16756.t1 | XP_001650333.1 tubulin beta chain isoform X2 [Aedes aegypti] | PTHR11588 | TUBULIN | ribonucleotide binding#GO:0032553;structural molecule activity#GO:0005198;organic cyclic compound binding#GO:0097159;anion binding#GO:0043168;GTP binding#GO:0005525;structural constituent of cytoskeleton#GO:0005200;guanyl ribonucleotide binding#GO:0032561;purine ribonucleoside triphosphate binding#GO:0035639 |
| chr1 | 3468273 | 3470364 | Aedes_aegypti_formosus\|g10626.t1 | XP_021696860.1 uncharacterized protein LOC5565294 [Aedes aegypti] | PTHR34153 | SI:CH211-262H13.3-RELATED-RELATED | |
| chr1 | 13617641 | 13618798 | Aedes_aegypti_formosus\|g10855.t1 | XP_001660335.1 BTB/POZ domain-containing protein 3 [Aedes aegypti] | PTHR45774 | BTB/POZ DOMAIN-CONTAINING | |
| chr1 | 13619007 | 13620146 | Aedes_aegypti_formosus\|g10856.t1 | XP_001660336.1 kelch-like protein 40a [Aedes aegypti] | PTHR45774 | BTB/POZ DOMAIN-CONTAINING | |
| chr1 | 282350345 | 282351518 | Aedes_aegypti_formosus\|g16506.t1 | XP_021706460.1 uncharacterized protein LOC110678149 [Aedes aegypti] | PTHR46486 | CCHC-TYPE DOMAIN-CONTAINING PROTEIN | |
| chr1 | 271103742 | 271228183 | Aedes_aegypti_formosus\|g16277.t1 | XP_021693720.1 probable multidrug resistance-associated protein lethal(2)03659 [ | PTHR24223 | ATP-BINDING CASSETTE SUB-FAMILY C | ATP hydrolysis activity#GO:0016887;nucleoside-triphosphatase activity#GO:0017111;ATP-dependent activity#GO:0140657;transporter activity#GO:0005215;catalytic activity#GO:0003824;active transmembrane transporter activity#GO:0022804 |
| chr1 | 271278113 | 271282405 | Aedes_aegypti_formosus\|g16279.t1 | XP_021693721.1 multidrug resistance-associated protein 4 [Aedes aegypti] | PTHR24223 | ATP-BINDING CASSETTE SUB-FAMILY C | ATP hydrolysis activity#GO:0016887;nucleoside-triphosphatase activity#GO:0017111;ATP-dependent activity#GO:0140657;transporter activity#GO:0005215;catalytic activity#GO:0003824;active transmembrane transporter activity#GO:0022804 |
| chr1 | 271295474 | 271331820 | Aedes_aegypti_formosus\|g16280.t1 | XP_021693723.1 probable multidrug resistance-associated protein lethal(2)03659 [ | PTHR24223 | ATP-BINDING CASSETTE SUB-FAMILY C | ATP hydrolysis activity#GO:0016887;nucleoside-triphosphatase activity#GO:0017111;ATP-dependent activity#GO:0140657;transporter activity#GO:0005215;catalytic activity#GO:0003824;active transmembrane transporter activity#GO:0022804 |
| chr3 | 250830930 | 250831387 | Aedes_aegypti_formosus\|g6545.t1 | XP_001655719.1 general odorant-binding protein 56d [Aedes aegypti] | PTHR11857 | ODORANT BINDING PROTEIN-RELATED | |
| chr3 | 250855838 | 250856295 | Aedes_aegypti_formosus\|g6547.t1 | XP_001655721.2 general odorant-binding protein 56d [Aedes aegypti] | PTHR11857 | ODORANT BINDING PROTEIN-RELATED | |
| chr1 | 6149900 | 6150928 | Aedes_aegypti_formosus\|g10698.t1 | XP_021699121.1 uncharacterized protein LOC110676260 [Aedes aegypti] | PTHR23002 | ZINC FINGER CCHC DOMAIN CONTAINING PROTEIN | |
| chr3 | 394026019 | 394027489 | Aedes_aegypti_formosus\|g9683.t1 | XP_001652408.2 putative polypeptide N-acetylgalactosaminyltransferase 9 [Aedes a | PTHR11675 | N-ACETYLGALACTOSAMINYLTRANSFERASE | |
| chr1 | 8858527 | 8871531 | Aedes_aegypti_formosus\|g10757.t1 | XP_021694535.1 RNA-binding protein squid isoform X1 [Aedes aegypti] | PTHR48033 | RNA-BINDING (RRM/RBD/RNP MOTIFS) FAMILY PROTEIN | |
| chr1 | 4058246 | 4089952 | Aedes_aegypti_formosus\|g10649.t1 | XP_001655300.2 DNA polymerase subunit gamma-2, mitochondrial [Aedes aegypti] | PTHR10745 | GLYCYL-TRNA SYNTHETASE/DNA POLYMERASE SUBUNIT GAMMA-2 | catalytic activity, acting on a nucleic acid#GO:0140640;catalytic activity, acting on RNA#GO:0140098;ligase activity#GO:0016874 |
| chr1 | 281847726 | 281874144 | Aedes_aegypti_formosus\|g16498.t1 | XP_021693554.1 dynein beta chain, ciliary isoform X1 [Aedes aegypti] | PTHR10676 | DYNEIN HEAVY CHAIN FAMILY PROTEIN | microtubule motor activity#GO:0003777;ATP-dependent activity#GO:0140657;binding#GO:0005488;catalytic activity#GO:0003824;cytoskeletal motor activity#GO:0003774;protein binding#GO:0005515;nucleoside-triphosphatase activity#GO:0017111;ATP hydrolysis activity#GO:0016887 |
| chr1 | 5308030 | 5314228 | Aedes_aegypti_formosus\|g10679.t1 | XP_021694435.1 acetylcholine receptor subunit alpha-like isoform X4 [Aedes aegyp | PTHR18945 | NEUROTRANSMITTER GATED ION CHANNEL | |
| chr1 | 13938589 | 13950147 | Aedes_aegypti_formosus\|g10868.t1 | XP_001655172.1 xenotropic and polytropic retrovirus receptor 1 [Aedes aegypti] | PTHR10783 | XENOTROPIC AND POLYTROPIC RETROVIRUS RECEPTOR 1-RELATED | |
| chr1 | 284843586 | 284846281 | Aedes_aegypti_formosus\|g16536.t1 | XP_021703123.1 uncharacterized protein LOC110677020 [Aedes aegypti] | PTHR31025 | SI:CH211-196P9.1-RELATED | |
| chr1 | 268212349 | 268224306 | Aedes_aegypti_formosus\|g16196.t1 | XP_001662091.2 uncharacterized protein LOC5575602 [Aedes aegypti] | PTHR33964 | RE45066P-RELATED | |
| chr1 | 271360702 | 271362687 | Aedes_aegypti_formosus\|g16281.t1 | XP_001662532.2 zinc finger protein 431 [Aedes aegypti] | PTHR16515 | PR DOMAIN ZINC FINGER PROTEIN | |
| chr1 | 11200836 | 11201603 | Aedes_aegypti_formosus\|g10803.t1 | XP_001660128.1 alpha-(1,3)-fucosyltransferase C [Aedes aegypti] | PTHR11929 | ALPHA- 1,3 -FUCOSYLTRANSFERASE | |
| chr1 | 11204278 | 11215247 | Aedes_aegypti_formosus\|g10804.t1 | XP_001660128.1 alpha-(1,3)-fucosyltransferase C [Aedes aegypti] | PTHR11929 | ALPHA- 1,3 -FUCOSYLTRANSFERASE | |
| chr1 | 285829458 | 285832798 | Aedes_aegypti_formosus\|g16553.t1 | XP_001654379.2 putative protein tag-278 [Aedes aegypti] | PTHR23159 | CENTROSOMAL PROTEIN 2 | |
| chr1 | 274311315 | 274312709 | Aedes_aegypti_formosus\|g16348.t1 | XP_021693682.1 5-hydroxytryptamine receptor 1 [Aedes aegypti] | PTHR24247 | 5-HYDROXYTRYPTAMINE RECEPTOR | cation binding#GO:0043169;G protein-coupled amine receptor activity#GO:0008227;heterocyclic compound binding#GO:1901363;neurotransmitter receptor activity#GO:0030594;neurotransmitter binding#GO:0042165;organic cyclic compound binding#GO:0097159 |
| chr1 | 275020173 | 275021591 | Aedes_aegypti_formosus\|g16359.t1 | XP_021693677.1 5-hydroxytryptamine receptor 1 [Aedes aegypti] | PTHR24247 | 5-HYDROXYTRYPTAMINE RECEPTOR | cation binding#GO:0043169;G protein-coupled amine receptor activity#GO:0008227;heterocyclic compound binding#GO:1901363;neurotransmitter receptor activity#GO:0030594;neurotransmitter binding#GO:0042165;organic cyclic compound binding#GO:0097159 |
| chr1 | 273450212 | 273451159 | Aedes_aegypti_formosus\|g16331.t1 | XP_021693694.1 serine protease 7 isoform X1 [Aedes aegypti] | PTHR24256 | TRYPTASE-RELATED | |
| chr1 | 273466130 | 273466840 | Aedes_aegypti_formosus\|g16332.t1 | XP_021693694.1 serine protease 7 isoform X1 [Aedes aegypti] | PTHR24256 | TRYPTASE-RELATED | |
| chr1 | 273501420 | 273525037 | Aedes_aegypti_formosus\|g16334.t1 | XP_001658784.2 serine protease easter [Aedes aegypti] | PTHR24256 | TRYPTASE-RELATED | |
| chr1 | 270355858 | 270376248 | Aedes_aegypti_formosus\|g16261.t1 | XP_001649986.2 uncharacterized protein LOC5565630 [Aedes aegypti] | PTHR21177 | IP06524P-RELATED | |
| chr1 | 270389863 | 270390996 | Aedes_aegypti_formosus\|g16262.t1 | XP_001662716.2 uncharacterized protein LOC5576523 [Aedes aegypti] | PTHR21177 | IP06524P-RELATED | |
| chr1 | 14203062 | 14216992 | Aedes_aegypti_formosus\|g10877.t1 | XP_001655181.1 aminoacylase-1B isoform X1 [Aedes aegypti] | PTHR45892 | AMINOACYLASE-1 | hydrolase activity#GO:0016787;hydrolase activity, acting on carbon-nitrogen (but not peptide) bonds, in linear amides#GO:0016811 |
| chr1 | 12252992 | 12254988 | Aedes_aegypti_formosus\|g10825.t1 | XP_001660136.1 uncharacterized family 31 glucosidase KIAA1161 [Aedes aegypti] | PTHR43053 | GLYCOSIDASE FAMILY 31 | |
| chr1 | 273030432 | 273057598 | Aedes_aegypti_formosus\|g16324.t1 | XP_021693695.1 uncharacterized protein LOC5566210 [Aedes aegypti] | PTHR45598 | PROTEIN CBG11839-RELATED | |
| chr1 | 1157834 | 1182955 | Aedes_aegypti_formosus\|g10546.t1 | XP_001658560.1 uncharacterized protein LOC5569474 [Aedes aegypti] | PTHR43668 | ALLANTOINASE | hydrolase activity, acting on carbon-nitrogen (but not peptide) bonds, in cyclic amides#GO:0016812 |
| chr1 | 14483524 | 14495035 | Aedes_aegypti_formosus\|g10886.t1 | XP_001650617.1 40S ribosomal protein S14 [Aedes aegypti] | PTHR11759 | 40S RIBOSOMAL PROTEIN S14/30S RIBOSOMAL PROTEIN S11 | |
| chr1 | 277827180 | 277830338 | Aedes_aegypti_formosus\|g16418.t1 | XP_021693638.1 serine-arginine protein 55 isoform X1 [Aedes aegypti] | PTHR23003 | RNA RECOGNITION MOTIF RRM DOMAIN CONTAINING PROTEIN | mRNA binding#GO:0003729;organic cyclic compound binding#GO:0097159 |
| chr1 | 277839718 | 277841396 | Aedes_aegypti_formosus\|g16420.t1 | XP_021693640.1 serine-arginine protein 55 isoform X2 [Aedes aegypti] | PTHR23003 | RNA RECOGNITION MOTIF RRM DOMAIN CONTAINING PROTEIN | mRNA binding#GO:0003729;organic cyclic compound binding#GO:0097159 |
| chr1 | 4943300 | 4945550 | Aedes_aegypti_formosus\|g10664.t1 | XP_021698571.1 uncharacterized protein LOC110676140 [Aedes aegypti] | PTHR33053 | PROTEIN, PUTATIVE-RELATED | |
| chr1 | 5117949 | 5119457 | Aedes_aegypti_formosus\|g10670.t1 | XP_001655853.1 26S proteasome regulatory subunit 4 [Aedes aegypti] | PTHR23073 | 26S PROTEASOME REGULATORY SUBUNIT | ATP hydrolysis activity#GO:0016887;nucleoside-triphosphatase activity#GO:0017111;ATP-dependent activity#GO:0140657;catalytic activity#GO:0003824 |
| chr1 | 7230323 | 7271064 | Aedes_aegypti_formosus\|g10723.t1 | XP_021694561.1 uncharacterized protein LOC5579825 isoform X4 [Aedes aegypti] | PTHR15696 | SMG-7 SUPPRESSOR WITH MORPHOLOGICAL EFFECT ON GENITALIA PROTEIN 7 | telomeric DNA binding#GO:0042162;RNA binding#GO:0003723 |
| chr1 | 7519716 | 7542403 | Aedes_aegypti_formosus\|g10731.t1 | XP_021694561.1 uncharacterized protein LOC5579825 isoform X4 [Aedes aegypti] | PTHR15696 | SMG-7 SUPPRESSOR WITH MORPHOLOGICAL EFFECT ON GENITALIA PROTEIN 7 | telomeric DNA binding#GO:0042162;RNA binding#GO:0003723 |
| chr1 | 7638009 | 7651054 | Aedes_aegypti_formosus\|g10733.t1 | XP_021694560.1 uncharacterized protein LOC5579825 isoform X3 [Aedes aegypti] | PTHR15696 | SMG-7 SUPPRESSOR WITH MORPHOLOGICAL EFFECT ON GENITALIA PROTEIN 7 | telomeric DNA binding#GO:0042162;RNA binding#GO:0003723 |

| | | | | | | | | |
|---|---|---|---|---|---|---|---|---|
| chr1 | 4056582 | 4058206 | Aedes_aegypti_formosus\|g10648.t1 | XP_021694720.1 | origin recognition complex subunit 5 [Aedes aegypti] | PTHR12705 | ORIGIN RECOGNITION COMPLEX SUBUNIT 5 | DNA replication origin binding#GO:0003688;double-stranded DNA binding#GO:0003690 |
| chr3 | 393956725 | 393958107 | Aedes_aegypti_formosus\|g9681.t1 | XP_021711548.1 | protein kinase C, brain isozyme isoform X3 [Aedes aegypti] | PTHR35248 | PUTATIVE-RELATED | |
| chr1 | 283747727 | 283748680 | Aedes_aegypti_formosus\|g16525.t1 | XP_021704183.1 | jerky protein homolog-like [Aedes aegypti] | PTHR19303 | TRANSPOSON | |
| chr1 | 283748748 | 283749359 | Aedes_aegypti_formosus\|g16526.t1 | XP_021704183.1 | jerky protein homolog-like [Aedes aegypti] | PTHR19303 | TRANSPOSON | |
| chr3 | 394093007 | 394106948 | Aedes_aegypti_formosus\|g9687.t1 | XP_021711531.1 | serine-enriched protein isoform X2 [Aedes aegypti] | PTHR24410 | HL07962P-RELATED | |
| chr1 | 277845438 | 277859115 | Aedes_aegypti_formosus\|g16421.t1 | XP_021693634.1 | mitochondrial dicarboxylate carrier [Aedes aegypti] | PTHR45618 | MITOCHONDRIAL DICARBOXYLATE CARRIER-RELATED | |
| chr3 | 11233943 | 11235601 | Aedes_aegypti_formosus\|g1391.t1 | XP_001648679.1 | apyrase [Aedes aegypti] | PTHR11575 | 5'-NUCLEOTIDASE-RELATED | |
| chr3 | 396314566 | 396324957 | Aedes_aegypti_formosus\|g9746.t1 | XP_021704184.1 | mitochondrial import receptor subunit TOM20 homolog [Aedes aegypt | PTHR12430 | MITOCHONDRIAL IMPORT RECEPTOR SUBUNIT TOM20 | signal sequence binding#GO:0005048;protein transmembrane transporter activity#GO:0008320;binding#GO:0005488;amide binding#GO:0033218 |
| chr1 | 294471024 | 294471710 | Aedes_aegypti_formosus\|g16753.t1 | XP_021693902.1 | uncharacterized protein LOC5565952 isoform X2 [Aedes aegypti] | PTHR31553 | NF-KAPPA-B ESSENTIAL MODULATOR | K63-linked polyubiquitin modification-dependent protein binding#GO:0070530;binding#GO:0005488;polyubiquitin modification-dependent protein binding#GO:0031593 |
| chr1 | 7113249 | 7114754 | Aedes_aegypti_formosus\|g10720.t1 | XP_001651058.2 | uncharacterized protein LOC5579827 [Aedes aegypti] | PTHR45789 | FI18025P1 | RNA polymerase II cis-regulatory region sequence-specific DNA binding#GO:0000978;transcription regulator activity#GO:0140110;DNA-binding transcription factor activity, RNA polymerase II-specific#GO:0000981;organic cyclic compound binding#GO:0097159 |
| chr1 | 7142124 | 7143487 | Aedes_aegypti_formosus\|g10721.t1 | XP_021712895.1 | uncharacterized protein LOC110681442 [Aedes aegypti] | PTHR45789 | FI18025P1 | RNA polymerase II cis-regulatory region sequence-specific DNA binding#GO:0000978;transcription regulator activity#GO:0140110;DNA-binding transcription factor activity, RNA polymerase II-specific#GO:0000981;organic cyclic compound binding#GO:0097159 |
| chr1 | 430418 | 431245 | Aedes_aegypti_formosus\|g10525.t1 | XP_021694649.1 | casein kinase I [Aedes aegypti] | PTHR11909 | CASEIN KINASE-RELATED | |
| chr1 | 1668132 | 1670462 | Aedes_aegypti_formosus\|g10555.t1 | XP_021694694.1 | cell cycle checkpoint protein RAD17 [Aedes aegypti] | PTHR12172 | CELL CYCLE CHECKPOINT PROTEIN RAD17 | binding#GO:0005488;chromatin binding#GO:0003682 |
| chr1 | 2311184 | 2321882 | Aedes_aegypti_formosus\|g10598.t1 | XP_021694695.1 | solute carrier family 35 member E1 homolog [Aedes aegypti] | PTHR11132 | SOLUTE CARRIER FAMILY 35 | |
| chr1 | 2343551 | 2344552 | Aedes_aegypti_formosus\|g10599.t1 | XP_021694695.1 | solute carrier family 35 member E1 homolog [Aedes aegypti] | PTHR11132 | SOLUTE CARRIER FAMILY 35 | |
| chr1 | 3426075 | 3427190 | Aedes_aegypti_formosus\|g10625.t1 | XP_001655294.1 | NKAP family protein CG6066 [Aedes aegypti] | PTHR13087 | NF-KAPPA B ACTIVATING PROTEIN | |
| chr1 | 3777895 | 3793375 | Aedes_aegypti_formosus\|g10637.t1 | XP_021694715.1 | nuclear factor of activated T-cells 5 isoform X5 [Aedes aegypti] | PTHR21616 | CENTROSOME SPINDLE POLE ASSOCIATED PROTEIN | |
| chr1 | 3931277 | 3946565 | Aedes_aegypti_formosus\|g10644.t1 | XP_021694711.1 | nuclear factor of activated T-cells 5 isoform X1 [Aedes aegypti] | PTHR21616 | CENTROSOME SPINDLE POLE ASSOCIATED PROTEIN | |
| chr1 | 282997405 | 283103826 | Aedes_aegypti_formosus\|g16517.t1 | XP_021694322.1 | synaptogenesis protein syg-2 [Aedes aegypti] | PTHR23278 | SIDESTEP PROTEIN | |
| chr1 | 283128909 | 283221552 | Aedes_aegypti_formosus\|g16518.t1 | XP_021694322.1 | synaptogenesis protein syg-2 [Aedes aegypti] | PTHR23278 | SIDESTEP PROTEIN | |
| chr1 | 281648207 | 281649514 | Aedes_aegypti_formosus\|g16490.t1 | XP_021693560.1 | protein fork head [Aedes aegypti] | PTHR11829 | FORKHEAD BOX PROTEIN | RNA polymerase II cis-regulatory region sequence-specific DNA binding#GO:0000978;transcription regulator activity#GO:0140110;DNA-binding transcription factor activity, RNA polymerase II-specific#GO:0000981;organic cyclic compound binding#GO:0097159 |
| chr1 | 281661903 | 281663210 | Aedes_aegypti_formosus\|g16492.t1 | XP_021693559.1 | protein fork head [Aedes aegypti] | PTHR11829 | FORKHEAD BOX PROTEIN | RNA polymerase II cis-regulatory region sequence-specific DNA binding#GO:0000978;transcription regulator activity#GO:0140110;DNA-binding transcription factor activity, RNA polymerase II-specific#GO:0000981;organic cyclic compound binding#GO:0097159 |
| chr1 | 310548232 | 310589087 | Aedes_aegypti_formosus\|g17328.t1 | XP_001648552.1 | fatty-acid amide hydrolase 2 [Aedes aegypti] | PTHR43372 | FATTY-ACID AMIDE HYDROLASE | |
| chr3 | 250861418 | 250861875 | Aedes_aegypti_formosus\|g6548.t1 | XP_001655722.2 | general odorant-binding protein 56a [Aedes aegypti] | PTHR11857 | ODORANT BINDING PROTEIN-RELATED | |
| chr3 | 250877327 | 250877786 | Aedes_aegypti_formosus\|g6549.t1 | XP_001655723.2 | general odorant-binding protein 56a [Aedes aegypti] | PTHR11857 | ODORANT BINDING PROTEIN-RELATED | |
| chr3 | 396783528 | 396792345 | Aedes_aegypti_formosus\|g9758.t1 | XP_001654420.1 | polyadenylate-binding protein [Aedes aegypti] | PTHR24012 | RNA BINDING PROTEIN | |
| chr1 | 278489322 | 278505231 | Aedes_aegypti_formosus\|g16434.t1 | XP_021693609.1 | vacuolar protein sorting-associated protein 13B [Aedes aegypti] | PTHR12517 | VACUOLAR PROTEIN SORTING-ASSOCIATED PROTEIN 13B | |
| chr2 | 9500709 | 9542549 | Aedes_aegypti_formosus\|g18078.t1 | XP_021695569.1 | uncharacterized protein LOC5567816 [Aedes aegypti] | PTHR23279 | DEFECTIVE PROBOSCIS EXTENSION RESPONSE DPR -RELATED | |
| chr1 | 275132614 | 275134639 | Aedes_aegypti_formosus\|g16360.t1 | XP_001650792.2 | odorant receptor 4 [Aedes aegypti] | PTHR21137 | ODORANT RECEPTOR | signaling receptor activity#GO:0038023;olfactory receptor activity#GO:0004984;transmembrane signaling receptor activity#GO:0004888 |
| chr1 | 15767900 | 15782948 | Aedes_aegypti_formosus\|g10908.t1 | XP_021694470.1 | myc protein [Aedes aegypti] | PTHR45851 | MYC PROTO-ONCOGENE | RNA polymerase II cis-regulatory region sequence-specific DNA binding#GO:0000978;transcription regulator activity#GO:0140110;DNA-binding transcription factor activity, RNA polymerase II-specific#GO:0000981;organic cyclic compound binding#GO:0097159 |
| chr1 | 15789547 | 15790884 | Aedes_aegypti_formosus\|g10909.t1 | XP_021694470.1 | myc protein [Aedes aegypti] | PTHR45851 | MYC PROTO-ONCOGENE | RNA polymerase II cis-regulatory region sequence-specific DNA binding#GO:0000978;transcription regulator activity#GO:0140110;DNA-binding transcription factor activity, RNA polymerase II-specific#GO:0000981;organic cyclic compound binding#GO:0097159 |
| chr1 | 12935282 | 13012632 | Aedes_aegypti_formosus\|g10840.t1 | XP_021694488.1 | uncharacterized protein LOC5572379 [Aedes aegypti] | PTHR21261 | BEAT PROTEIN | |
| chr1 | 8724101 | 8731100 | Aedes_aegypti_formosus\|g10754.t1 | XP_021694539.1 | carbonic anhydrase 2 [Aedes aegypti] | PTHR18952 | CARBONIC ANHYDRASE | |
| chr1 | 8652956 | 8676381 | Aedes_aegypti_formosus\|g10751.t1 | XP_021694540.1 | IWS1-like protein [Aedes aegypti] | PTHR46010 | PROTEIN IWS1 HOMOLOG | |
| chr1 | 8433198 | 8479500 | Aedes_aegypti_formosus\|g10746.t1 | XP_021694544.1 | synaptotagmin-14 isoform X2 [Aedes aegypti] | PTHR46129 | SYNAPTOTAGMIN 14, ISOFORM D | |
| chr1 | 783900 | 806150 | Aedes_aegypti_formosus\|g10537.t1 | XP_021694658.1 | probable phosphorylase b kinase regulatory subunit alpha isoform | PTHR10749 | PHOSPHORYLASE B KINASE REGULATORY SUBUNIT | |
| chr1 | 812887 | 833596 | Aedes_aegypti_formosus\|g10538.t1 | XP_001658556.1 | phosphatidylinositide phosphatase SAC1 [Aedes aegypti] | PTHR45662 | PHOSPHATIDYLINOSITIDE PHOSPHATASE SAC1 | phosphatase activity#GO:0016791 |
| chr1 | 869008 | 886602 | Aedes_aegypti_formosus\|g10540.t1 | XP_001658556.1 | phosphatidylinositide phosphatase SAC1 [Aedes aegypti] | PTHR45662 | PHOSPHATIDYLINOSITIDE PHOSPHATASE SAC1 | phosphatase activity#GO:0016791 |
| chr1 | 895356 | 896794 | Aedes_aegypti_formosus\|g10541.t1 | XP_001658557.1 | p21-activated protein kinase-interacting protein 1-like [Aedes ae | PTHR44675 | PAK1 INTERACTING PROTEIN 1 | |
| chr1 | 840667 | 859226 | Aedes_aegypti_formosus\|g10539.t1 | XP_021694661.1 | T-complex protein 1 subunit beta [Aedes aegypti] | PTHR11353 | CHAPERONIN | protein binding#GO:0005515;unfolded protein binding#GO:0051082;binding#GO:0005488 |
| chr1 | 3998261 | 4031201 | Aedes_aegypti_formosus\|g10646.t1 | XP_021712271.1 | GPI transamidase component PIG-T-like [Aedes aegypti] | PTHR12959 | GPI TRANSAMIDASE COMPONENT PIG-T-RELATED | |
| chr1 | 4031300 | 4056281 | Aedes_aegypti_formosus\|g10647.t1 | XP_021712272.1 | gastrula zinc finger protein XlCGF57.1-like [Aedes aegypti] | PTHR24384 | FINGER PUTATIVE TRANSCRIPTION FACTOR FAMILY-RELATED | RNA polymerase II cis-regulatory region sequence-specific DNA binding#GO:0000978;transcription regulator activity#GO:0140110;DNA-binding transcription factor activity, RNA polymerase II-specific#GO:0000981;organic cyclic compound binding#GO:0097159 |
| chr1 | 48934238 | 48935395 | Aedes_aegypti_formosus\|g11708.t1 | XP_001658571.2 | 28S ribosomal protein S29, mitochondrial [Aedes aegypti] | PTHR12810 | MITOCHONDRIAL 28S RIBOSOMAL PROTEIN S29 | structural constituent of ribosome#GO:0003735 |
| chr1 | 48865887 | 48868082 | Aedes_aegypti_formosus\|g11706.t1 | XP_001658568.1 | uncharacterized protein LOC5569483 [Aedes aegypti] | PTHR24155 | OSTEOCLAST-STIMULATING FACTOR 1 | |
| chr1 | 285559509 | 285576896 | Aedes_aegypti_formosus\|g16547.t1 | XP_021693535.1 | tyrosine-protein kinase Fer isoform X1 [Aedes aegypti] | PTHR24418 | TYROSINE-PROTEIN KINASE | |
| chr1 | 280702058 | 280714457 | Aedes_aegypti_formosus\|g16464.t1 | XP_021693584.1 | uncharacterized protein LOC5572405 [Aedes aegypti] | PTHR23003 | RNA RECOGNITION MOTIF RRM DOMAIN CONTAINING PROTEIN | mRNA binding#GO:0003729;organic cyclic compound binding#GO:0097159 |
| chr1 | 280741960 | 280750423 | Aedes_aegypti_formosus\|g16465.t1 | XP_021693583.1 | uncharacterized protein LOC5572408 [Aedes aegypti] | PTHR23003 | RNA RECOGNITION MOTIF RRM DOMAIN CONTAINING PROTEIN | mRNA binding#GO:0003729;organic cyclic compound binding#GO:0097159 |
| chr1 | 268940534 | 268941206 | Aedes_aegypti_formosus\|g16220.t1 | XP_001662086.2 | probable maleylacetoacetate isomerase 2 isoform X1 [Aedes aegypti | PTHR42673 | MALEYLACETOACETATE ISOMERASE | glutathione transferase activity#GO:0004364;isomerase activity#GO:0016853 |
| chr3 | 12807560 | 12819990 | Aedes_aegypti_formosus\|g1422.t1 | XP_021706701.1 | DET1 homolog [Aedes aegypti] | PTHR13374 | DET1 HOMOLOG DE-ETIOLATED-1 HOMOLOG | molecular adaptor activity#GO:0060090;ubiquitin protein ligase binding#GO:0031625;protein-macromolecule adaptor activity#GO:0030674 |
| chr3 | 12550538 | 12551235 | Aedes_aegypti_formosus\|g1417.t1 | XP_021708952.1 | ATP-dependent Clp protease ATP-binding subunit clpX-like, mitochc | PTHR48102 | ATP-DEPENDENT CLP PROTEASE ATP-BINDING SUBUNIT CLPX-LIKE, MITOCHONDRIAL-RELATED | |
| chr3 | 397161155 | 397163158 | Aedes_aegypti_formosus\|g9769.t1 | XP_021704308.1 | uncharacterized protein LOC5573208 isoform X2 [Aedes aegypti] | PTHR22934 | PROTEIN ESC1/WETA-RELATED | |
| chr3 | 397176467 | 397186301 | Aedes_aegypti_formosus\|g9770.t1 | XP_021704307.1 | uncharacterized protein LOC5573208 isoform X1 [Aedes aegypti] | PTHR22934 | PROTEIN ESC1/WETA-RELATED | |
| chr1 | 285855878 | 285856335 | Aedes_aegypti_formosus\|g16555.t1 | XP_001654377.1 | uncharacterized protein LOC5573057 [Aedes aegypti] | PTHR23353 | RAB-GAP/TBC-RELATED | |
| chr2 | 192652422 | 192667001 | Aedes_aegypti_formosus\|g22128.t1 | XP_021699362.1 | dendritic arbor reduction protein 1 isoform X1 [Aedes aegypti] | PTHR40201 | OSMIOPHILIC BODY PROTEIN | |
| chr1 | 271626119 | 271627073 | Aedes_aegypti_formosus\|g16298.t1 | XP_021693697.1 | RNA-binding protein cabeza-like isoform X1 [Aedes aegypti] | PTHR23238 | RNA BINDING PROTEIN | transcription coregulator activity#GO:0003712;RNA binding#GO:0003723 |
| chr1 | 278703418 | 278706846 | Aedes_aegypti_formosus\|g16444.t1 | XP_021701955.1 | uncharacterized protein LOC110676845 [Aedes aegypti] | PTHR46481 | ZINC FINGER BED DOMAIN-CONTAINING PROTEIN 4 | |
| chr1 | 310468831 | 310472412 | Aedes_aegypti_formosus\|g17325.t1 | XP_021701955.1 | uncharacterized protein LOC110676845 [Aedes aegypti] | PTHR46481 | ZINC FINGER BED DOMAIN-CONTAINING PROTEIN 4 | |
| chr1 | 16407689 | 16409246 | Aedes_aegypti_formosus\|g10917.t1 | XP_021694468.1 | 5-demethoxyubiquinone hydroxylase, mitochondrial [Aedes aegypti] | PTHR11237 | COENZYME Q10 BIOSYNTHESIS PROTEIN 7 | |
| chr1 | 14626584 | 14627874 | Aedes_aegypti_formosus\|g10889.t1 | XP_001650620.1 | methionyl-tRNA formyltransferase, mitochondrial [Aedes aegypti] | PTHR11138 | METHIONYL-TRNA FORMYLTRANSFERASE | catalytic activity, acting on a nucleic acid#GO:0140640;catalytic activity, acting on RNA#GO:0140098;transferase activity#GO:0016740;transferase activity, transferring one-carbon groups#GO:0016741 |
| chr1 | 14603369 | 14617986 | Aedes_aegypti_formosus\|g10888.t1 | XP_001650619.1 | uncharacterized protein LOC5566213 isoform X1 [Aedes aegypti] | PTHR28646 | TRANSMEMBRANE PROTEIN 201 | actin filament binding#GO:0051015 |
| chr1 | 12306927 | 12307698 | Aedes_aegypti_formosus\|g10828.t1 | XP_001660139.1 | uncharacterized protein LOC5572032 [Aedes aegypti] | PTHR16296 | UNCHARACTERIZED HYPOTHALAMUS PROTEIN HT007 | |
| chr1 | 12236798 | 12238190 | Aedes_aegypti_formosus\|g10824.t1 | XP_001660135.1 | PCI domain-containing protein 2 homolog [Aedes aegypti] | PTHR12732 | UNCHARACTERIZED PROTEASOME COMPONENT REGION PCI-CONTAINING | double-stranded DNA binding#GO:0003690;RNA binding#GO:0003723;organic cyclic compound binding#GO:0097159;DNA binding#GO:0003677 |
| chr1 | 9341106 | 9374914 | Aedes_aegypti_formosus\|g10776.t1 | XP_021694527.1 | protein artichoke [Aedes aegypti] | PTHR24366 | IG(IMMUNOGLOBULIN) AND LRR(LEUCINE RICH REPEAT) DOMAINS | |
| chr1 | 8824620 | 8835685 | Aedes_aegypti_formosus\|g10756.t1 | XP_001651045.1 | ras GTPase-activating protein-binding protein 1 isoform X1 [Aedes | PTHR10693 | RAS GTPASE-ACTIVATING PROTEIN-BINDING PROTEIN | mRNA binding#GO:0003729;heterocyclic compound binding#GO:1901363;organic cyclic compound binding#GO:0097159 |
| chr1 | 7172308 | 7173426 | Aedes_aegypti_formosus\|g10722.t1 | XP_021712896.1 | paired box protein Pax-6-like isoform X1 [Aedes aegypti] | PTHR45636 | PAIRED BOX PROTEIN PAX-6-RELATED-RELATED | RNA polymerase II cis-regulatory region sequence-specific DNA binding#GO:0000978;transcription regulator activity#GO:0140110;DNA-binding transcription factor activity, RNA polymerase II-specific#GO:0000981;organic cyclic compound binding#GO:0097159 |
| chr1 | 6163604 | 6217371 | Aedes_aegypti_formosus\|g10700.t1 | XP_021694588.1 | zinc finger protein 2 [Aedes aegypti] | PTHR45891 | ZINC FINGER HOMEOBOX PROTEIN | RNA polymerase II cis-regulatory region sequence-specific DNA binding#GO:0000978;transcription regulator activity#GO:0140110;DNA-binding transcription factor activity, RNA polymerase II-specific#GO:0000981;organic cyclic compound binding#GO:0097159 |
| chr1 | 1650398 | 1651261 | Aedes_aegypti_formosus\|g10553.t1 | XP_001658562.1 | activator of basal transcription 1 [Aedes aegypti] | PTHR12311 | ACTIVATOR OF BASAL TRANSCRIPTION 1 | nucleic acid binding#GO:0003676;RNA binding#GO:0003723;organic cyclic compound binding#GO:0097159 |
| chr1 | 2355101 | 2359523 | Aedes_aegypti_formosus\|g10600.t1 | XP_001661571.1 | von Willebrand factor A domain-containing protein 8 [Aedes aegypt | PTHR21610 | VON WILLEBRAND FACTOR A DOMAIN-CONTAINING PROTEIN 8 | |
| chr1 | 2388591 | 2420085 | Aedes_aegypti_formosus\|g10601.t1 | XP_001661570.1 | elongation factor 1-gamma [Aedes aegypti] | PTHR43986 | ELONGATION FACTOR 1-GAMMA | |

| | | | | | | | | |
|---|---|---|---|---|---|---|---|---|
| chr1 | 3995565 | 3997409 | Aedes_aegypti_formosus\|g10645.t1 | XP_021694719.1 | carnitine O-acetyltransferase [Aedes aegypti] | PTHR22589 | CARNITINE O-ACYLTRANSFERASE | |
| chr1 | 285793358 | 285817183 | Aedes_aegypti_formosus\|g16552.t1 | XP_021693530.1 | AN1-type zinc finger protein 6 [Aedes aegypti] | PTHR10634 | AN1-TYPE ZINC FINGER PROTEIN | |
| chr1 | 282568317 | 282576418 | Aedes_aegypti_formosus\|g16510.t1 | XP_021693553.1 | ankyrin repeat domain-containing protein 12 isoform X6 [Aedes aeg | PTHR24178 | MOLTING PROTEIN MLT-4 | |
| chr1 | 282591976 | 282648328 | Aedes_aegypti_formosus\|g16511.t1 | XP_021693549.1 | ankyrin repeat domain-containing protein 12 isoform X2 [Aedes aeg | PTHR24178 | MOLTING PROTEIN MLT-4 | |
| chr1 | 278607082 | 278608623 | Aedes_aegypti_formosus\|g16442.t1 | XP_021693606.1 | endochitinase A [Aedes aegypti] | PTHR46827 | HEAT SHOCK PROTEIN DDB_G0288861-RELATED | |
| chr1 | 278568407 | 278569816 | Aedes_aegypti_formosus\|g16439.t1 | XP_021693611.1 | nucleotide exchange factor SIL1 [Aedes aegypti] | PTHR19316 | PROTEIN FOLDING REGULATOR | |
| chr1 | 275550970 | 275566210 | Aedes_aegypti_formosus\|g16366.t1 | XP_021693675.1 | trichohyalin [Aedes aegypti] | PTHR35970 | SODIUM CHANNEL AND CLATHRIN LINKER 1 | |
| chr1 | 271600842 | 271621143 | Aedes_aegypti_formosus\|g16297.t1 | XP_001662540.2 | protein bric-a-brac 1 isoform X1 [Aedes aegypti] | PTHR23110 | BTB DOMAIN TRANSCRIPTION FACTOR | |
| chr1 | 271461720 | 271469437 | Aedes_aegypti_formosus\|g16287.t1 | XP_021693716.1 | uncharacterized protein LOC5576202 isoform X1 [Aedes aegypti] | PTHR35420 | OS02G0198500 PROTEIN | |
| chr1 | 271494677 | 271496026 | Aedes_aegypti_formosus\|g16289.t1 | XP_021693717.1 | uncharacterized protein LOC5576202 isoform X2 [Aedes aegypti] | PTHR35420 | OS02G0198500 PROTEIN | |
| chr1 | 270244676 | 270319080 | Aedes_aegypti_formosus\|g16258.t1 | XP_021693736.1 | probable Rho GTPase-activating protein CG5521 isoform X1 [Aedes a | PTHR10063 | TUBERIN | GTPase activity#GO:0003924;nucleoside-triphosphatase regulator activity#GO:0060589;GTPase regulator activity#GO:0030695;GTPase activator activity#GO:0005096;hydrolase activity, acting on acid anhydrides#GO:0016817;protein binding#GO:0005515;enzyme activator activity#GO:0008047;nucleoside-triphosphatase activity#GO:0017111 |
| chr1 | 269308206 | 269344000 | Aedes_aegypti_formosus\|g16234.t1 | XP_021693764.1 | alpha-mannosidase 2 isoform X1 [Aedes aegypti] | PTHR11607 | ALPHA-MANNOSIDASE | hydrolase activity#GO:0016787;hydrolase activity, hydrolyzing O-glycosyl compounds#GO:0004553 |
| chr1 | 310617570 | 310629663 | Aedes_aegypti_formosus\|g17330.t1 | XP_001654115.2 | exocyst complex component 2 [Aedes aegypti] | PTHR13043 | EXOCYST COMPLEX COMPONENT SEC5 | |
| chr1 | 310263018 | 310305350 | Aedes_aegypti_formosus\|g17319.t1 | XP_021694180.1 | uncharacterized protein LOC5564410 [Aedes aegypti] | PTHR14905 | NG37 | |
| chr1 | 310313644 | 310325323 | Aedes_aegypti_formosus\|g17322.t1 | XP_021694180.1 | uncharacterized protein LOC5564410 [Aedes aegypti] | PTHR14905 | NG37 | |
| chr1 | 12436132 | 12437112 | Aedes_aegypti_formosus\|g10830.t1 | XP_001660142.1 | forkhead box protein D3 [Aedes aegypti] | PTHR11829 | FORKHEAD BOX PROTEIN | RNA polymerase II cis-regulatory region sequence-specific DNA binding#GO:0000978;transcription regulator activity#GO:0140110;DNA-binding transcription factor activity, RNA polymerase II-specific#GO:0000981;organic cyclic compound binding#GO:0097159 |
| chr3 | 12563790 | 12564424 | Aedes_aegypti_formosus\|g1418.t1 | XP_021710544.1 | tumor necrosis factor alpha-induced protein 8-like protein isofor | PTHR12757 | TUMOR NECROSIS FACTOR INDUCED PROTEIN | |
| chr3 | 12327366 | 12346960 | Aedes_aegypti_formosus\|g1412.t1 | XP_021704809.1 | DNA topoisomerase 2-binding protein 1-B [Aedes aegypti] | PTHR13561 | DNA REPLICATION REGULATOR DPB11-RELATED | |
| chr3 | 12303927 | 12320439 | Aedes_aegypti_formosus\|g1411.t1 | XP_001660595.1 | T-complex protein 1 subunit gamma [Aedes aegypti] | PTHR11353 | CHAPERONIN | protein binding#GO:0005515;unfolded protein binding#GO:0051082;binding#GO:0005488 |
| chr3 | 11816453 | 11818349 | Aedes_aegypti_formosus\|g1399.t1 | XP_001660604.1 | signal recognition particle subunit SRP68 [Aedes aegypti] | PTHR12860 | SIGNAL RECOGNITION PARTICLE 68 KDA PROTEIN | ribonucleoprotein complex binding#GO:0043021;protein-containing complex binding#GO:0044877;binding#GO:0005488 |
| chr3 | 104257265 | 104328586 | Aedes_aegypti_formosus\|g3523.t1 | XP_021706399.1 | neuronal PAS domain-containing protein 4 [Aedes aegypti] | PTHR23043 | HYPOXIA-INDUCIBLE FACTOR 1 ALPHA | |
| chr3 | 250891332 | 250891871 | Aedes_aegypti_formosus\|g6550.t1 | XP_001655725.2 | general odorant-binding protein 56d [Aedes aegypti] | PTHR11857 | ODORANT BINDING PROTEIN-RELATED | |
| chr3 | 250903556 | 250904095 | Aedes_aegypti_formosus\|g6551.t1 | XP_001655725.2 | general odorant-binding protein 56d [Aedes aegypti] | PTHR11857 | ODORANT BINDING PROTEIN-RELATED | |
| chr3 | 396823827 | 396847425 | Aedes_aegypti_formosus\|g9759.t1 | XP_001654423.1 | tudor domain-containing protein 7A isoform X1 [Aedes aegypti] | PTHR22948 | TUDOR DOMAIN CONTAINING PROTEIN | |
| chr3 | 394140482 | 394142263 | Aedes_aegypti_formosus\|g9689.t1 | XP_001652404.1 | alpha-amylase 4N [Aedes aegypti] | PTHR43447 | ALPHA-AMYLASE | |
| chr1 | 5263963 | 5264991 | Aedes_aegypti_formosus\|g10678.t1 | XP_001655863.1 | casein kinase II subunit alpha [Aedes aegypti] | PTHR24054 | CASEIN KINASE II SUBUNIT ALPHA | |
| chr1 | 5202542 | 5211942 | Aedes_aegypti_formosus\|g10677.t1 | XP_021694441.1 | synaptic vesicle glycoprotein 2C isoform X1 [Aedes aegypti] | PTHR23511 | SYNAPTIC VESICLE GLYCOPROTEIN 2 | |
| chr1 | 5185118 | 5187087 | Aedes_aegypti_formosus\|g10675.t1 | XP_001655859.2 | tetratricopeptide repeat protein 21B isoform X2 [Aedes aegypti] | PTHR14699 | STI2 PROTEIN-RELATED | |
| chr1 | 5159321 | 5163224 | Aedes_aegypti_formosus\|g10674.t1 | XP_001655856.2 | aminopeptidase N isoform X1 [Aedes aegypti] | PTHR11533 | PROTEASE M1 ZINC METALLOPROTEASE | cation binding#GO:0043169;metallopeptidase activity#GO:0008237;zinc ion binding#GO:0008270;peptide binding#GO:0042277;aminopeptidase activity#GO:0004177;exopeptidase activity#GO:0008238;peptidase activity#GO:0008233;catalytic activity, acting on a protein#GO:0140096 |
| chr1 | 5104374 | 5107987 | Aedes_aegypti_formosus\|g10669.t1 | XP_001655852.1 | zinc finger protein-like 1 homolog [Aedes aegypti] | PTHR12981 | ZINC FINGER PROTEIN-LIKE 1 | |
| chr1 | 5102924 | 5104241 | Aedes_aegypti_formosus\|g10668.t1 | XP_001655851.1 | sperm-associated antigen 1 [Aedes aegypti] | PTHR45984 | RNA (RNA) POLYMERASE II ASSOCIATED PROTEIN HOMOLOG | protein binding#GO:0005515;heat shock protein binding#GO:0031072;binding#GO:0005488 |
| chr1 | 4880182 | 4903624 | Aedes_aegypti_formosus\|g10662.t1 | XP_021694458.1 | abl interactor 2 [Aedes aegypti] | PTHR10460 | ABL INTERACTOR FAMILY MEMBER | |
| chr1 | 4547376 | 4561964 | Aedes_aegypti_formosus\|g10654.t1 | XP_021694462.1 | pyrokinin-1 receptor [Aedes aegypti] | PTHR24243 | G-PROTEIN COUPLED RECEPTOR | |
| chr1 | 4104341 | 4105238 | Aedes_aegypti_formosus\|g10651.t1 | XP_001655319.2 | vacuolar-sorting protein SNF8 [Aedes aegypti] | PTHR12806 | EAP30 SUBUNIT OF ELL COMPLEX | |
| chr1 | 16409490 | 16418317 | Aedes_aegypti_formosus\|g10918.t1 | XP_021694466.1 | protein shifted isoform X1 [Aedes aegypti] | PTHR14949 | EGF-LIKE-DOMAIN, MULTIPLE 7, 8 | |
| chr1 | 16382014 | 16398514 | Aedes_aegypti_formosus\|g10914.t1 | XP_021708813.1 | mitochondrial cardiolipin hydrolase [Aedes aegypti] | PTHR43856 | CARDIOLIPIN HYDROLASE | endoribonuclease activity#GO:0004521 |
| chr1 | 14699470 | 14729035 | Aedes_aegypti_formosus\|g10891.t1 | XP_021694471.1 | tRNA dimethylallyltransferase, mitochondrial [Aedes aegypti] | PTHR11088 | TRNA DIMETHYLALLYLTRANSFERASE | catalytic activity, acting on a nucleic acid#GO:0140640;catalytic activity, acting on RNA#GO:0140098;transferase activity, transferring alkyl or aryl (other than methyl) groups#GO:0016765 |
| chr1 | 14661473 | 14664703 | Aedes_aegypti_formosus\|g10890.t1 | XP_001650621.2 | sodium-dependent neutral amino acid transporter B(0)AT3 isoform X | PTHR11616 | SODIUM/CHLORIDE DEPENDENT TRANSPORTER | |
| chr1 | 14539778 | 14555428 | Aedes_aegypti_formosus\|g10887.t1 | XP_021694475.1 | rab11 family-interacting protein 5 isoform X1 [Aedes aegypti] | PTHR15746 | RAB11-RELATED | |
| chr1 | 14460645 | 14483215 | Aedes_aegypti_formosus\|g10885.t1 | XP_001650614.1 | dynactin subunit 4 [Aedes aegypti] | PTHR13034 | DYNACTIN P62 SUBUNIT | |
| chr1 | 14418188 | 14441537 | Aedes_aegypti_formosus\|g10884.t1 | XP_021694479.1 | glucose-induced degradation protein 4 homolog [Aedes aegypti] | PTHR14534 | VACUOLAR IMPORT AND DEGRADATION PROTEIN 24 | |
| chr1 | 14367724 | 14392074 | Aedes_aegypti_formosus\|g10883.t1 | XP_021694480.1 | uncharacterized protein LOC5574533 [Aedes aegypti] | PTHR14387 | THADA/DEATH RECEPTOR INTERACTING PROTEIN | |
| chr1 | 14176521 | 14177393 | Aedes_aegypti_formosus\|g10873.t1 | XP_001655177.1 | max-like protein X [Aedes aegypti] | PTHR15741 | BASIC HELIX-LOOP-HELIX ZIP TRANSCRIPTION FACTOR | RNA polymerase II cis-regulatory region sequence-specific DNA binding#GO:0000978;transcription regulator activity#GO:0140110;DNA-binding transcription factor activity, RNA polymerase II-specific#GO:0000981;organic cyclic compound binding#GO:0097159 |
| chr1 | 13972052 | 13986472 | Aedes_aegypti_formosus\|g10869.t1 | XP_001655173.1 | 39S ribosomal protein L38, mitochondrial [Aedes aegypti] | PTHR11362 | PHOSPHATIDYLETHANOLAMINE-BINDING PROTEIN | |
| chr1 | 13621918 | 13623862 | Aedes_aegypti_formosus\|g10857.t1 | XP_001660337.2 | xaa-Pro aminopeptidase ApepP isoform X2 [Aedes aegypti] | PTHR43763 | XAA-PRO AMINOPEPTIDASE 1 | |
| chr1 | 12339625 | 12370240 | Aedes_aegypti_formosus\|g10829.t1 | XP_021694496.1 | nuclear factor 1 isoform X1 [Aedes aegypti] | PTHR11492 | NUCLEAR FACTOR I | RNA polymerase II cis-regulatory region sequence-specific DNA binding#GO:0000978;transcription regulator activity#GO:0140110;DNA-binding transcription factor activity, RNA polymerase II-specific#GO:0000981;organic cyclic compound binding#GO:0097159 |
| chr1 | 11224519 | 11239782 | Aedes_aegypti_formosus\|g10806.t1 | XP_001660130.1 | probable cytosolic oligopeptidase A [Aedes aegypti] | PTHR11804 | PROTEASE M3 THIMET OLIGOPEPTIDASE-RELATED | metalloendopeptidase activity#GO:0004222;peptidase activity#GO:0008233;catalytic activity, acting on a protein#GO:0140096 |
| chr1 | 11215335 | 11216434 | Aedes_aegypti_formosus\|g10805.t1 | XP_001660129.1 | glyoxalase domain-containing protein 4 [Aedes aegypti] | PTHR46466 | GLYOXALASE DOMAIN-CONTAINING PROTEIN 4 | |
| chr1 | 11153015 | 11155341 | Aedes_aegypti_formosus\|g10802.t1 | XP_001660127.1 | protein stoned-A [Aedes aegypti] | PTHR10649 | ARYL HYDROCARBON RECEPTOR | signaling receptor activity#GO:0038023;transcription regulator activity#GO:0140110;RNA polymerase II transcription regulatory region sequence-specific DNA binding#GO:0000977;DNA-binding transcription factor activity, RNA polymerase II-specific#GO:0000981;organic cyclic compound binding#GO:0097159 |
| chr1 | 11148593 | 11152913 | Aedes_aegypti_formosus\|g10801.t1 | XP_021694526.1 | protein stoned-B [Aedes aegypti] | PTHR10529 | AP COMPLEX SUBUNIT MU | |
| chr1 | 8941404 | 8943013 | Aedes_aegypti_formosus\|g10761.t1 | XP_001651036.1 | venom allergen 5 [Aedes aegypti] | PTHR10334 | CYSTEINE-RICH SECRETORY PROTEIN-RELATED | |
| chr1 | 8736973 | 8783248 | Aedes_aegypti_formosus\|g10755.t1 | XP_001651046.2 | LOW QUALITY PROTEIN: rho-associated protein kinase 1 [Aedes aegyp | PTHR22988 | MYOTONIC DYSTROPHY S/T KINASE-RELATED | protein serine/threonine kinase activity#GO:0004674;phosphotransferase activity, alcohol group as acceptor#GO:0016773;catalytic activity, acting on a protein#GO:0140096;protein kinase activity#GO:0004672 |
| chr1 | 8558050 | 8581493 | Aedes_aegypti_formosus\|g10749.t1 | XP_021694542.1 | putative adenosylhomocysteinase 3 isoform X2 [Aedes aegypti] | PTHR23420 | ADENOSYLHOMOCYSTEINASE | hydrolase activity#GO:0016787;catalytic activity#GO:0003824 |
| chr1 | 8546850 | 8548406 | Aedes_aegypti_formosus\|g10748.t1 | XP_001651053.1 | lysosomal Pro-X carboxypeptidase [Aedes aegypti] | PTHR11010 | PROTEASE S28 PRO-X CARBOXYPEPTIDASE-RELATED | |
| chr1 | 8367489 | 8377488 | Aedes_aegypti_formosus\|g10743.t1 | XP_021694554.1 | microtubule-associated protein futsch isoform X1 [Aedes aegypti] | PTHR46439 | CYSTEINE-RICH MOTOR NEURON 1 PROTEIN | |
| chr1 | 7089757 | 7111561 | Aedes_aegypti_formosus\|g10719.t1 | XP_001661348.1 | protein unc-13 homolog C isoform X4 [Aedes aegypti] | PTHR10480 | PROTEIN UNC-13 HOMOLOG | syntaxin binding#GO:0019905;calmodulin binding#GO:0005516;binding#GO:0005488;SNARE binding#GO:0000149 |
| chr1 | 6619310 | 6744101 | Aedes_aegypti_formosus\|g10712.t1 | XP_021694573.1 | plexin-A2 [Aedes aegypti] | PTHR22625 | PLEXIN | |
| chr1 | 6421262 | 6438657 | Aedes_aegypti_formosus\|g10707.t1 | XP_021694580.1 | zinc finger protein 236 isoform X1 [Aedes aegypti] | PTHR24388 | ZINC FINGER PROTEIN | |
| chr1 | 6047262 | 6071420 | Aedes_aegypti_formosus\|g10694.t1 | XP_021694595.1 | tau-tubulin kinase homolog Asator [Aedes aegypti] | PTHR11909 | CASEIN KINASE-RELATED | |
| chr1 | 5581434 | 5583488 | Aedes_aegypti_formosus\|g10687.t1 | XP_001661364.1 | mannose-6-phosphate isomerase [Aedes aegypti] | PTHR10309 | MANNOSE-6-PHOSPHATE ISOMERASE | isomerase activity#GO:0016853;catalytic activity#GO:0003824 |
| chr1 | 510728 | 523628 | Aedes_aegypti_formosus\|g10529.t1 | XP_001658550.2 | putative vitellogenin receptor [Aedes aegypti] | PTHR22722 | LOW-DENSITY LIPOPROTEIN RECEPTOR-RELATED PROTEIN 2-RELATED | |
| chr1 | 1720498 | 1749952 | Aedes_aegypti_formosus\|g10558.t1 | XP_021712338.1 | proline-rich protein 36 [Aedes aegypti] | PTHR14429 | FIBROSIN FAMILY MEMBER | |
| chr1 | 2285716 | 2310302 | Aedes_aegypti_formosus\|g10596.t1 | XP_001661572.1 | vitamin K epoxide reductase complex subunit 1 [Aedes aegypti] | PTHR14519 | VITAMIN K EPOXIDE REDUCTASE COMPLEX, SUBUNIT 1 | oxidoreductase activity#GO:0016491 |
| chr1 | 2495969 | 2551653 | Aedes_aegypti_formosus\|g10602.t1 | XP_021694697.1 | uncharacterized protein LOC5574653 [Aedes aegypti] | PTHR24006 | UBIQUITIN CARBOXYL-TERMINAL HYDROLASE | |
| chr1 | 2642607 | 2645690 | Aedes_aegypti_formosus\|g10605.t1 | XP_001662326.1 | aminopeptidase N [Aedes aegypti] | PTHR11533 | PROTEASE M1 ZINC METALLOPROTEASE | cation binding#GO:0043169;metallopeptidase activity#GO:0008237;zinc ion binding#GO:0008270;peptide binding#GO:0042277;aminopeptidase activity#GO:0004177;exopeptidase activity#GO:0008238;peptidase activity#GO:0008233;catalytic activity, acting on a protein#GO:0140096 |
| chr1 | 3061264 | 3086974 | Aedes_aegypti_formosus\|g10614.t1 | XP_021694706.1 | rhophilin-2 isoform X2 [Aedes aegypti] | PTHR23031 | RHOPHILIN | |
| chr1 | 3112120 | 3181453 | Aedes_aegypti_formosus\|g10615.t1 | XP_001648600.1 | membrane-associated progesterone receptor component 1 [Aedes aegy | PTHR10281 | MEMBRANE-ASSOCIATED PROGESTERONE RECEPTOR COMPONENT-RELATED | |
| chr1 | 3218835 | 3220229 | Aedes_aegypti_formosus\|g10618.t1 | XP_001648601.2 | GDP-fucose protein O-fucosyltransferase 2 [Aedes aegypti] | PTHR13398 | GDP-FUCOSE PROTEIN O-FUCOSYLTRANSFERASE 2 | fucosyltransferase activity#GO:0008417;catalytic activity, acting on a protein#GO:0140096 |
| chr1 | 3350654 | 3351752 | Aedes_aegypti_formosus\|g10622.t1 | XP_021694709.1 | ubiquinone biosynthesis monooxygenase COQ6, mitochondrial [Aedes | PTHR43876 | UBIQUINONE BIOSYNTHESIS MONOOXYGENASE COQ6, MITOCHONDRIAL | oxidoreductase activity#GO:0016491;catalytic activity#GO:0003824 |
| chr1 | 192230151 | 192235906 | Aedes_aegypti_formosus\|g14534.t1 | XP_001660544.1 | 60S ribosomal protein L4 [Aedes aegypti] | PTHR19431 | 60S RIBOSOMAL PROTEIN L4 | structural constituent of ribosome#GO:0003735;RNA binding#GO:0003723 |
| chr1 | 192227908 | 192229816 | Aedes_aegypti_formosus\|g14533.t1 | XP_001660542.1 | pre-mRNA-splicing factor SPF27 [Aedes aegypti] | PTHR13296 | BCAS2 PROTEIN | |
| chr1 | 191549285 | 191565368 | Aedes_aegypti_formosus\|g14523.t1 | XP_021711828.1 | B-box type zinc finger protein ncl-1 [Aedes aegypti] | PTHR25462 | BONUS, ISOFORM C-RELATED | |
| chr1 | 286069505 | 286120721 | Aedes_aegypti_formosus\|g16561.t1 | XP_021693525.1 | E3 ubiquitin-protein ligase RNF19B isoform X3 [Aedes aegypti] | PTHR11685 | RBR FAMILY RING FINGER AND IBR DOMAIN-CONTAINING | ubiquitin-like protein conjugating enzyme binding#GO:0044390;catalytic activity, acting on a protein#GO:0140096;binding#GO:0005488;ubiquitin-like protein transferase activity#GO:0019787;ubiquitin protein ligase activity#GO:0061630 |
| chr1 | 285283608 | 285307983 | Aedes_aegypti_formosus\|g16544.t1 | XP_021693543.1 | phosphopantothenate--cysteine ligase [Aedes aegypti] | PTHR12290 | CORNICHON-RELATED | |
| chr1 | 281241985 | 281244684 | Aedes_aegypti_formosus\|g16486.t1 | XP_001656471.2 | translation initiation factor IF-2, mitochondrial [Aedes aegypti] | PTHR43381 | TRANSLATION INITIATION FACTOR IF-2-RELATED | |
| chr1 | 280763699 | 280801868 | Aedes_aegypti_formosus\|g16466.t1 | XP_021693565.1 | papilin isoform X3 [Aedes aegypti] | PTHR13723 | ADAMTS A DISINTEGRIN AND METALLOPROTEASE WITH THROMBOSPONDIN MOTIFS PROTEASE | |

| | | | | | | | | |
|---|---|---|---|---|---|---|---|---|
| chr1 | 278598747 | 278599718 | Aedes_aegypti_formosus\|g16441.t1 | XP_021693607.1 | thioredoxin domain-containing protein 9 [Aedes aegypti] | PTHR21148 | THIOREDOXIN DOMAIN-CONTAINING PROTEIN 9 | |
| chr1 | 278511315 | 278523029 | Aedes_aegypti_formosus\|g16436.t1 | XP_001660875.1 | serine--pyruvate aminotransferase, mitochondrial [Aedes aegypti] | PTHR21152 | AMINOTRANSFERASE CLASS V | transaminase activity#GO:0008483;catalytic activity#GO:0003824 |
| chr1 | 278093121 | 278095725 | Aedes_aegypti_formosus\|g16427.t1 | XP_021693629.1 | zinc metalloproteinase-disintegrin-like EoMP06 [Aedes aegypti] | PTHR11905 | ADAM A DISINTEGRIN AND METALLOPROTEASE DOMAIN | |
| chr1 | 277870927 | 277881155 | Aedes_aegypti_formosus\|g16423.t1 | XP_021693631.1 | uncharacterized protein LOC5576567 isoform X2 [Aedes aegypti] | PTHR11905 | ADAM A DISINTEGRIN AND METALLOPROTEASE DOMAIN | |
| chr1 | 277859320 | 277860057 | Aedes_aegypti_formosus\|g16422.t1 | XP_021693635.1 | elongator complex protein 6 isoform X1 [Aedes aegypti] | PTHR16184 | ELONGATOR COMPLEX PROTEIN 6 | |
| chr1 | 277808898 | 277810929 | Aedes_aegypti_formosus\|g16417.t1 | XP_001662752.1 | zinc finger protein 777 [Aedes aegypti] | PTHR24408 | ZINC FINGER PROTEIN | transcription regulator activity#GO:0140110;RNA polymerase II transcription regulatory region sequence-specific DNA binding#GO:0000977;DNA-binding transcription factor activity, RNA polymerase II-specific#GO:0000981;organic cyclic compound binding#GO:0097159 |
| chr1 | 277071682 | 277072660 | Aedes_aegypti_formosus\|g16399.t1 | XP_001663476.1 | cyclin-dependent kinase 2 [Aedes aegypti] | PTHR24056 | CELL DIVISION PROTEIN KINASE | protein serine/threonine kinase activity#GO:0004674;cyclin-dependent protein kinase activity#GO:0097472;cyclin binding#GO:0030332;protein kinase activity#GO:0004672;cyclin-dependent protein serine/threonine kinase activity#GO:0004693 |
| chr1 | 275698796 | 275700724 | Aedes_aegypti_formosus\|g16369.t1 | XP_001663291.2 | fibroblast growth factor receptor-like 1 [Aedes aegypti] | PTHR19890 | FIBROBLAST GROWTH FACTOR RECEPTOR | |
| chr1 | 274210060 | 274211646 | Aedes_aegypti_formosus\|g16346.t1 | XP_021693684.1 | uncharacterized protein LOC110674159 [Aedes aegypti] | PTHR31639 | F-BOX PROTEIN-LIKE | |
| chr1 | 274034383 | 274087401 | Aedes_aegypti_formosus\|g16343.t1 | XP_001660239.2 | elongation of very long chain fatty acids protein AAEL008004 isof | PTHR11157 | FATTY ACID ACYL TRANSFERASE-RELATED | acyltransferase activity#GO:0016746;acyltransferase activity, transferring groups other than amino-acyl groups#GO:0016747 |
| chr1 | 273540993 | 273575198 | Aedes_aegypti_formosus\|g16335.t1 | XP_021693693.1 | elongation of very long chain fatty acids protein AAEL008004 isof | PTHR11157 | FATTY ACID ACYL TRANSFERASE-RELATED | acyltransferase activity#GO:0016746;acyltransferase activity, transferring groups other than amino-acyl groups#GO:0016747 |
| chr1 | 271669952 | 271725792 | Aedes_aegypti_formosus\|g16301.t1 | XP_001662541.1 | N(G),N(G)-dimethylarginine dimethylaminohydrolase 1 [Aedes aegypt | PTHR12737 | DIMETHYLARGININE DIMETHYLAMINOHYDROLASE | hydrolase activity#GO:0016787;hydrolase activity, acting on carbon-nitrogen (but not peptide) bonds#GO:0016810;small molecule binding#GO:0036094;binding#GO:0005488 |
| chr1 | 271558789 | 271570879 | Aedes_aegypti_formosus\|g16294.t1 | XP_001662538.1 | uncharacterized protein LOC5576199 [Aedes aegypti] | PTHR22844 | F-BOX AND WD40 DOMAIN PROTEIN | |
| chr1 | 271520747 | 271523838 | Aedes_aegypti_formosus\|g16290.t1 | XP_001662536.1 | glucosidase 2 subunit beta [Aedes aegypti] | PTHR12630 | N-LINKED OLIGOSACCHARIDE PROCESSING | |
| chr1 | 271377444 | 271378827 | Aedes_aegypti_formosus\|g16284.t1 | XP_021693719.1 | huntingtin-interacting protein K [Aedes aegypti] | PTHR31184 | HUNTINGTIN-INTERACTING PROTEIN K FAMILY MEMBER | |
| chr1 | 270574800 | 270575222 | Aedes_aegypti_formosus\|g16269.t1 | XP_001662708.2 | putative defense protein 3 [Aedes aegypti] | PTHR45828 | CYTOCHROME B561/FERRIC REDUCTASE TRANSMEMBRANE | oxidoreductase activity, acting on metal ions#GO:0016722 |
| chr1 | 270531359 | 270532504 | Aedes_aegypti_formosus\|g16267.t1 | XP_001662710.2 | probable phosphoserine aminotransferase [Aedes aegypti] | PTHR43247 | PHOSPHOSERINE AMINOTRANSFERASE | transaminase activity#GO:0008483;heterocyclic compound binding#GO:1901363;anion binding#GO:0043168;small molecule binding#GO:0036094;catalytic activity#GO:0003824;organic cyclic compound binding#GO:0097159 |
| chr1 | 270530707 | 270531321 | Aedes_aegypti_formosus\|g16266.t1 | XP_021693731.1 | guanosine-3',5'-bis(diphosphate) 3'-pyrophosphohydrolase MESH1 is | PTHR46246 | GUANOSINE-3',5'-BIS(DIPHOSPHATE) 3'-PYROPHOSPHOHYDROLASE MESH1 | hydrolase activity#GO:0016787;phosphoric ester hydrolase activity#GO:0042578 |
| chr1 | 270500577 | 270508006 | Aedes_aegypti_formosus\|g16265.t1 | XP_001649983.1 | pyruvate kinase isoform X1 [Aedes aegypti] | PTHR11817 | PYRUVATE KINASE | carbon-oxygen lyase activity#GO:0016835;isomerase activity#GO:0016853;phosphotransferase activity, alcohol group as acceptor#GO:0016773;oxidoreductase activity#GO:0016491;intramolecular transferase activity#GO:0016866;hydro-lyase activity#GO:0016836;kinase activity#GO:0016301 |
| chr1 | 270108782 | 270133794 | Aedes_aegypti_formosus\|g16255.t1 | XP_001662725.1 | aspartate aminotransferase, cytoplasmic [Aedes aegypti] | PTHR11879 | ASPARTATE AMINOTRANSFERASE | |
| chr1 | 270080082 | 270098316 | Aedes_aegypti_formosus\|g16253.t1 | XP_001655788.1 | G patch domain-containing protein 4 [Aedes aegypti] | PTHR23149 | G PATCH DOMAIN CONTAINING PROTEIN | |
| chr1 | 269970805 | 269978133 | Aedes_aegypti_formosus\|g16250.t1 | XP_021693743.1 | scaffold protein salvador [Aedes aegypti] | PTHR47522 | SALVADOR FAMILY WW DOMAIN-CONTAINING PROTEIN 1 | |
| chr1 | 269966222 | 269966884 | Aedes_aegypti_formosus\|g16248.t1 | XP_001655791.1 | calmodulin [Aedes aegypti] | PTHR23048 | MYOSIN LIGHT CHAIN 1, 3 | |
| chr1 | 269800452 | 269821699 | Aedes_aegypti_formosus\|g16244.t1 | XP_021693747.1 | uncharacterized protein LOC5575646 [Aedes aegypti] | PTHR15451 | ERGOSTEROL BIOSYNTHETIC PROTEIN 28-RELATED | protein binding#GO:0005515;protein-macromolecule adaptor activity#GO:0030674 |
| chr1 | 269536457 | 269589658 | Aedes_aegypti_formosus\|g16241.t1 | XP_021693759.1 | proline-, glutamic acid- and leucine-rich protein 1 [Aedes aegypt | PTHR34105 | PROLINE-, GLUTAMIC ACID- AND LEUCINE-RICH PROTEIN 1 | |
| chr1 | 269480296 | 269482880 | Aedes_aegypti_formosus\|g16238.t1 | XP_001648330.1 | HSPB1-associated protein 1 [Aedes aegypti] | PTHR12461 | HYPOXIA-INDUCIBLE FACTOR 1 ALPHA INHIBITOR-RELATED | |
| chr1 | 269391696 | 269441162 | Aedes_aegypti_formosus\|g16236.t1 | XP_021693762.1 | ubiquitin carboxyl-terminal hydrolase 34 [Aedes aegypti] | PTHR24006 | UBIQUITIN CARBOXYL-TERMINAL HYDROLASE | |
| chr1 | 269245495 | 269272977 | Aedes_aegypti_formosus\|g16232.t1 | XP_021693776.1 | RNA-binding protein Nova-1 isoform X7 [Aedes aegypti] | PTHR10288 | KH DOMAIN CONTAINING RNA BINDING PROTEIN | mRNA binding#GO:0003729;organic cyclic compound binding#GO:0097159 |
| chr1 | 294484641 | 294500952 | Aedes_aegypti_formosus\|g16754.t1 | XP_021693898.1 | GTP-binding protein 2 [Aedes aegypti] | PTHR43721 | ELONGATION FACTOR TU-RELATED | translation regulator activity, nucleic acid binding#GO:0090079;translation elongation factor activity#GO:0003746;translation factor activity, RNA binding#GO:0008135 |
| chr1 | 310494345 | 310511536 | Aedes_aegypti_formosus\|g17327.t1 | XP_021694181.1 | adenylosuccinate lyase [Aedes aegypti] | PTHR43172 | ADENYLOSUCCINATE LYASE | |
| chr1 | 310238570 | 310240293 | Aedes_aegypti_formosus\|g17316.t1 | XP_001654123.2 | spindle assembly abnormal protein 6 homolog [Aedes aegypti] | PTHR44281 | SPINDLE ASSEMBLY ABNORMAL PROTEIN 6 HOMOLOG | |
| chr3 | 12710688 | 12751408 | Aedes_aegypti_formosus\|g1419.t1 | XP_001659016.2 | zinc finger RNA-binding protein isoform X1 [Aedes aegypti] | PTHR45762 | ZINC FINGER RNA-BINDING PROTEIN | single-stranded RNA binding#GO:0003727;RNA binding#GO:0003723;double-stranded RNA binding#GO:0003725 |
| chr3 | 12406200 | 12431747 | Aedes_aegypti_formosus\|g1416.t1 | XP_021705203.1 | E3 ubiquitin-protein ligase RNF126 [Aedes aegypti] | PTHR45931 | SI:CH211-59O9.10 | |
| chr3 | 12381095 | 12393371 | Aedes_aegypti_formosus\|g1415.t1 | XP_001664206.2 | probable methylcrotonoyl-CoA carboxylase beta chain, mitochondria | PTHR22855 | ACETYL, PROPIONYL, PYRUVATE, AND GLUTACONYL CARBOXYLASE-RELATED | |
| chr3 | 12369548 | 12370197 | Aedes_aegypti_formosus\|g1414.t1 | XP_001664205.1 | 39S ribosomal protein L20, mitochondrial [Aedes aegypti] | PTHR10986 | 39S RIBOSOMAL PROTEIN L20 | structural constituent of ribosome#GO:0003735 |
| chr3 | 12353800 | 12356376 | Aedes_aegypti_formosus\|g1413.t1 | XP_001664204.1 | tRNA (cytosine(34)-C(5))-methyltransferase [Aedes aegypti] | PTHR22808 | NCL1 YEAST -RELATED NOL1/NOP2/FMU SUN DOMAIN CONTAINING | methyltransferase activity#GO:0008168;transferase activity, transferring one-carbon groups#GO:0016741;catalytic activity#GO:0003824 |
| chr3 | 12296329 | 12297642 | Aedes_aegypti_formosus\|g1410.t1 | XP_021710373.1 | protein disulfide-isomerase A6 homolog [Aedes aegypti] | PTHR45815 | PROTEIN DISULFIDE-ISOMERASE A6 | protein-disulfide reductase activity#GO:0015035;catalytic activity, acting on a protein#GO:0140096 |
| chr3 | 12287288 | 12289334 | Aedes_aegypti_formosus\|g1409.t1 | XP_001660598.1 | microfibrillar-associated protein 1 [Aedes aegypti] | PTHR15327 | MICROFIBRIL-ASSOCIATED PROTEIN | RNA binding#GO:0003723;heterocyclic compound binding#GO:1901363;organic cyclic compound binding#GO:0097159 |
| chr3 | 12270189 | 12282825 | Aedes_aegypti_formosus\|g1408.t1 | XP_021704348.1 | male-specific lethal 3 homolog isoform X2 [Aedes aegypti] | PTHR10880 | MORTALITY FACTOR 4-LIKE PROTEIN | |
| chr3 | 11839180 | 11841420 | Aedes_aegypti_formosus\|g1400.t1 | XP_021705712.1 | hsp90 co-chaperone Cdc37 [Aedes aegypti] | PTHR12800 | CDC37-RELATED | heat shock protein binding#GO:0031072;unfolded protein binding#GO:0051082;chaperone binding#GO:0051087 |
| chr3 | 11791288 | 11805127 | Aedes_aegypti_formosus\|g1398.t1 | XP_001660605.1 | ATP-binding cassette sub-family E member 1 [Aedes aegypti] | PTHR19248 | ATP-BINDING TRANSPORT PROTEIN-RELATED | |
| chr1 | 4280703 | 4282156 | Aedes_aegypti_formosus\|g10652.t1 | XP_021706380.1 | uncharacterized protein LOC110678093 [Aedes aegypti] | PTHR38758 | PUTATIVE-RELATED | |
| chr3 | 250780551 | 250781320 | Aedes_aegypti_formosus\|g6544.t1 | XP_001655718.2 | general odorant-binding protein 56d [Aedes aegypti] | PTHR11857 | ODORANT BINDING PROTEIN-RELATED | |
| chr3 | 394615116 | 394623088 | Aedes_aegypti_formosus\|g9697.t1 | XP_001651094.2 | LOW QUALITY PROTEIN: uncharacterized protein LOC5566687 [Aedes ae | PTHR46542 | X-BOX BINDING PROTEIN 1 | transcription regulator activity#GO:0140110;RNA polymerase II transcription regulatory region sequence-specific DNA binding#GO:0000977;DNA-binding transcription factor activity, RNA polymerase II-specific#GO:0000981;organic cyclic compound binding#GO:0097159 |
| chr3 | 394607490 | 394608693 | Aedes_aegypti_formosus\|g9696.t1 | XP_001651093.2 | GTPase Era, mitochondrial [Aedes aegypti] | PTHR42698 | GTPASE ERA | ribonucleoprotein complex binding#GO:0043021;RNA binding#GO:0003723;protein-containing complex binding#GO:0044877;rRNA binding#GO:0019843 |
| chr3 | 394452752 | 394510570 | Aedes_aegypti_formosus\|g9694.t1 | XP_001651092.2 | sodium-dependent serotonin transporter [Aedes aegypti] | PTHR11616 | SODIUM/CHLORIDE DEPENDENT TRANSPORTER | |
| chr3 | 394174606 | 394194365 | Aedes_aegypti_formosus\|g9690.t1 | XP_021711523.1 | putative polypeptide N-acetylgalactosaminyltransferase 9 isoform | PTHR11675 | N-ACETYLGALACTOSAMINYLTRANSFERASE | |
| chr3 | 394073013 | 394088407 | Aedes_aegypti_formosus\|g9686.t1 | XP_021711532.1 | putative polypeptide N-acetylgalactosaminyltransferase 9 [Aedes a | PTHR11675 | N-ACETYLGALACTOSAMINYLTRANSFERASE | |
| chr3 | 394049501 | 394061982 | Aedes_aegypti_formosus\|g9685.t1 | XP_001652407.2 | putative polypeptide N-acetylgalactosaminyltransferase 9 [Aedes a | PTHR11675 | N-ACETYLGALACTOSAMINYLTRANSFERASE | |
| chr3 | 393942718 | 393943765 | Aedes_aegypti_formosus\|g9680.t1 | XP_021711546.1 | protein kinase C, brain isozyme isoform X1 [Aedes aegypti] | PTHR24351 | RIBOSOMAL PROTEIN S6 KINASE | |
| chr3 | 393610248 | 393630143 | Aedes_aegypti_formosus\|g9676.t1 | XP_021711564.1 | putative inorganic phosphate cotransporter isoform X1 [Aedes aegy | PTHR11662 | SOLUTE CARRIER FAMILY 17 | |
| chr3 | 396454409 | 396464991 | Aedes_aegypti_formosus\|g9750.t1 | XP_011493380.2 | ecdysone-inducible protein E75 isoform X2 [Aedes aegypti] | PTHR24082 | NUCLEAR HORMONE RECEPTOR | |
| chr3 | 396349203 | 396385978 | Aedes_aegypti_formosus\|g9747.t1 | XP_021704178.1 | roundabout homolog 3 isoform X2 [Aedes aegypti] | PTHR10075 | BASIGIN RELATED | |
| chr3 | 396198520 | 396209305 | Aedes_aegypti_formosus\|g9743.t1 | XP_001652750.1 | UDP-glucose 4-epimerase [Aedes aegypti] | PTHR43725 | UDP-GLUCOSE 4-EPIMERASE | racemase and epimerase activity, acting on carbohydrates and derivatives#GO:0016857 |
| chr3 | 395662270 | 395671756 | Aedes_aegypti_formosus\|g9736.t1 | XP_021704263.1 | sex peptide receptor [Aedes aegypti] | PTHR47023 | SEX PEPTIDE RECEPTOR | |
| chr3 | 397193318 | 397194656 | Aedes_aegypti_formosus\|g9771.t1 | XP_001654438.2 | uncharacterized protein LOC5573222 [Aedes aegypti] | PTHR11506 | LYSOSOME-ASSOCIATED MEMBRANE GLYCOPROTEIN | |
| chr3 | 397141726 | 397144477 | Aedes_aegypti_formosus\|g9768.t1 | XP_021704310.1 | 40S ribosomal protein S12 [Aedes aegypti] | PTHR11843 | 40S RIBOSOMAL PROTEIN S12 | |
| chr3 | 397131864 | 397141636 | Aedes_aegypti_formosus\|g9767.t1 | XP_021704309.1 | zinc finger MYND domain-containing protein 10 homolog [Aedes aegy | PTHR13244 | ZINC FINGER MYND DOMAIN CONTAINING PROTEIN 10 | |
| chr3 | 397118206 | 397126446 | Aedes_aegypti_formosus\|g9766.t1 | XP_001654433.1 | uncharacterized protein LOC5573228 [Aedes aegypti] | PTHR12480 | ARGININE DEMETHYLASE AND LYSYL-HYDROXYLASE JMJD | |
| chr3 | 397111988 | 397113460 | Aedes_aegypti_formosus\|g9764.t1 | XP_001654431.1 | uncharacterized protein LOC5573216 [Aedes aegypti] | PTHR11557 | PORPHOBILINOGEN DEAMINASE | transferase activity, transferring alkyl or aryl (other than methyl) groups#GO:0016765;catalytic activity#GO:0003824 |
| chr3 | 397055352 | 397065233 | Aedes_aegypti_formosus\|g9762.t1 | XP_001654429.1 | probable ATP-dependent RNA helicase DDX27 [Aedes aegypti] | PTHR47959 | ATP-DEPENDENT RNA HELICASE RHLE-RELATED | |
| chr3 | 397021438 | 397043727 | Aedes_aegypti_formosus\|g9761.t1 | XP_021704328.1 | heat shock factor protein isoform X1 [Aedes aegypti] | PTHR10015 | HEAT SHOCK TRANSCRIPTION FACTOR | |
| chr1 | 5068096 | 5092246 | Aedes_aegypti_formosus\|g10667.t1 | XP_021694451.1 | aminopeptidase N isoform X2 [Aedes aegypti] | PTHR11533 | PROTEASE M1 ZINC METALLOPROTEASE | cation binding#GO:0043169;metallopeptidase activity#GO:0008237;zinc ion binding#GO:0008270;peptide binding#GO:0042277;aminopeptidase activity#GO:0004177;exopeptidase activity#GO:0008238;peptidase activity#GO:0008233;catalytic activity, acting on a protein#GO:0140096 |
| chr1 | 14337849 | 14338588 | Aedes_aegypti_formosus\|g10882.t1 | XP_021694481.1 | protein spitz [Aedes aegypti] | PTHR12332 | KEREN-RELATED | receptor ligand activity#GO:0048018;signaling receptor activator activity#GO:0030546;signaling receptor activity#GO:0038023 |
| chr1 | 14231205 | 14251207 | Aedes_aegypti_formosus\|g10878.t1 | XP_001655183.2 | DDB1- and CUL4-associated factor 7 [Aedes aegypti] | PTHR19919 | WD REPEAT CONTAINING PROTEIN | |
| chr1 | 13832001 | 13853245 | Aedes_aegypti_formosus\|g10864.t1 | XP_021694485.1 | cholesterol 7-desaturase [Aedes aegypti] | PTHR21266 | IRON-SULFUR DOMAIN CONTAINING PROTEIN | |
| chr1 | 13648574 | 13651855 | Aedes_aegypti_formosus\|g10858.t1 | XP_021694486.1 | cytochrome P450 307a1 [Aedes aegypti] | PTHR24303 | HEME-BINDING MONOOXYGENASE FAMILY | |
| chr1 | 13601000 | 13616665 | Aedes_aegypti_formosus\|g10853.t1 | XP_001660334.1 | ethanolamine kinase [Aedes aegypti] | PTHR22603 | CHOLINE/ETHANOALAMINE KINASE | phosphotransferase activity, alcohol group as acceptor#GO:0016773;kinase activity#GO:0016301 |
| chr1 | 13587714 | 13589750 | Aedes_aegypti_formosus\|g10851.t1 | XP_001660333.1 | lipoamide acyltransferase component of branched-chain alpha-keto | PTHR43178 | DIHYDROLIPOAMIDE ACETYLTRANSFERASE COMPONENT OF PYRUVATE DEHYDROGENASE COMPLEX | fatty acid binding#GO:0005504;acyltransferase activity, transferring groups other than amino-acyl groups#GO:0016747;acyltransferase activity#GO:0016746;acetyltransferase activity#GO:0016407;small molecule binding#GO:0036094;organic cyclic compound binding#GO:0097159;carboxylic acid binding#GO:0031406;heterocyclic compound binding#GO:1901363;lipid binding#GO:0008289 |

| | | | | | | | | |
|---|---|---|---|---|---|---|---|---|
| chr1 | 12065979 | 12095174 | Aedes_aegypti_formosus\|g10821.t1 | XP_021694516.1 | potassium voltage-gated channel protein Shaker isoform X4 [Aedes | PTHR11537 | VOLTAGE-GATED POTASSIUM CHANNEL | inorganic molecular entity transmembrane transporter activity#GO:0015318;potassium channel activity#GO:0005267;ion channel activity#GO:0005216;voltage-gated cation channel activity#GO:0022843;metal ion transmembrane transporter activity#GO:0046873;voltage-gated potassium channel activity#GO:0005249 |
| chr1 | 11257752 | 11272461 | Aedes_aegypti_formosus\|g10808.t1 | XP_001660132.1 | solute carrier family 25 member 35 isoform X1 [Aedes aegypti] | PTHR45928 | RE38146P | |
| chr1 | 11239882 | 11257362 | Aedes_aegypti_formosus\|g10807.t1 | XP_001660131.1 | dynein assembly factor 4, axonemal isoform X1 [Aedes aegypti] | PTHR46492 | DYNEIN ASSEMBLY FACTOR 4, AXONEMAL | |
| chr1 | 6476483 | 6478449 | Aedes_aegypti_formosus\|g10709.t1 | XP_001661352.2 | paired box protein Pax-6 isoform X2 [Aedes aegypti] | PTHR45636 | PAIRED BOX PROTEIN PAX-6-RELATED-RELATED | RNA polymerase II cis-regulatory region sequence-specific DNA binding#GO:0000978;transcription regulator activity#GO:0140110;DNA-binding transcription factor activity, RNA polymerase II-specific#GO:0000981;organic cyclic compound binding#GO:0097159 |
| chr1 | 5566847 | 5569820 | Aedes_aegypti_formosus\|g10686.t1 | XP_021694635.1 | peroxidase [Aedes aegypti] | PTHR11475 | OXIDASE/PEROXIDASE | oxidoreductase activity#GO:0016491;catalytic activity#GO:0003824 |
| chr1 | 1063955 | 1065648 | Aedes_aegypti_formosus\|g10545.t1 | XP_001658559.1 | histidine-rich glycoprotein [Aedes aegypti] | PTHR15283 | GREMLIN 1 | receptor ligand activity#GO:0048018;signaling receptor activator activity#GO:0030546;signaling receptor activity#GO:0038023;cytokine binding#GO:0019955 |
| chr1 | 1651315 | 1652937 | Aedes_aegypti_formosus\|g10554.t1 | XP_001658563.1 | zinc finger protein 160 [Aedes aegypti] | PTHR24399 | ZINC FINGER AND BTB DOMAIN-CONTAINING | |
| chr1 | 3408801 | 3411805 | Aedes_aegypti_formosus\|g10623.t1 | XP_021694708.1 | transient receptor potential channel pyrexia [Aedes aegypti] | PTHR47143 | TRANSIENT RECEPTOR POTENTIAL CATION CHANNEL PROTEIN PAINLESS | |
| chr1 | 278779691 | 278846180 | Aedes_aegypti_formosus\|g16445.t1 | XP_021693603.1 | oxysterol-binding protein-related protein 8 isoform X5 [Aedes aeg | PTHR10972 | OXYSTEROL-BINDING PROTEIN-RELATED | lipid transporter activity#GO:0005319;sterol binding#GO:0032934 |
| chr1 | 276447649 | 276484507 | Aedes_aegypti_formosus\|g16387.t1 | XP_021693657.1 | uncharacterized protein LOC5579517 [Aedes aegypti] | PTHR14952 | ROPPORIN-1-LIKE PROTEIN | |
| chr1 | 275738292 | 275740872 | Aedes_aegypti_formosus\|g16373.t1 | XP_001663293.2 | serine/arginine repetitive matrix protein 2 [Aedes aegypti] | PTHR43670 | HEAT SHOCK PROTEIN 26 | |
| chr1 | 270721844 | 270787513 | Aedes_aegypti_formosus\|g16272.t1 | XP_021693726.1 | uncharacterized protein LOC5575947 isoform X1 [Aedes aegypti] | PTHR39069 | ECDYSONE-INDUCIBLE GENE E1, ISOFORM A | |
| chr1 | 270429664 | 270431775 | Aedes_aegypti_formosus\|g16263.t1 | XP_001662715.2 | dimethyladenosine transferase 2, mitochondrial [Aedes aegypti] | PTHR11727 | DIMETHYLADENOSINE TRANSFERASE | catalytic activity, acting on a nucleic acid#GO:0140640;N-methyltransferase activity#GO:0008170;transferase activity, transferring one-carbon groups#GO:0016741;rRNA (adenine) methyltransferase activity#GO:0016433;S-adenosylmethionine-dependent methyltransferase activity#GO:0008757;RNA methyltransferase activity#GO:0008173 |
| chr1 | 270170185 | 270174497 | Aedes_aegypti_formosus\|g16256.t1 | XP_001662724.2 | zinc finger protein 62 [Aedes aegypti] | PTHR24379 | KRAB AND ZINC FINGER DOMAIN-CONTAINING | |
| chr1 | 269980019 | 270021817 | Aedes_aegypti_formosus\|g16251.t1 | XP_021693739.1 | src substrate cortactin-like isoform X1 [Aedes aegypti] | PTHR10829 | CORTACTIN AND DREBRIN | |
| chr1 | 269635305 | 269678835 | Aedes_aegypti_formosus\|g16243.t1 | XP_021693749.1 | nuclear transcription factor Y subunit beta isoform X3 [Aedes aeg | PTHR23104 | MULTIPLE COAGULATION FACTOR DEFICIENCY PROTEIN 2 NEURAL STEM CELL DERIVED NEURONAL SURVIVAL PROTEIN | |
| chr1 | 269476949 | 269480151 | Aedes_aegypti_formosus\|g16237.t1 | XP_021693760.1 | ionotropic receptor 40a [Aedes aegypti] | PTHR42643 | IONOTROPIC RECEPTOR 20A-RELATED | |
| chr1 | 310589406 | 310599079 | Aedes_aegypti_formosus\|g17329.t1 | XP_021694173.1 | uncharacterized protein LOC5572607 [Aedes aegypti] | PTHR48017 | OS05G0424000 PROTEIN-RELATED | |
| chr1 | 310240373 | 310240898 | Aedes_aegypti_formosus\|g17317.t1 | XP_001654122.2 | coiled-coil domain-containing protein 12 [Aedes aegypti] | PTHR31551 | PRE-MRNA-SPLICING FACTOR CWF18 | |
| chr3 | 12875814 | 12877062 | Aedes_aegypti_formosus\|g1424.t1 | XP_021711486.1 | GTP-binding protein Rheb homolog [Aedes aegypti] | PTHR24070 | RAS, DI-RAS, AND RHEB FAMILY MEMBERS OF SMALL GTPASE SUPERFAMILY | GTPase activity#GO:0003924;GDP binding#GO:0019003;GTP binding#GO:0005525;purine ribonucleoside triphosphate binding#GO:0035639;catalytic activity#GO:0003824 |
| chr3 | 396411001 | 396424647 | Aedes_aegypti_formosus\|g9749.t1 | XP_001652744.1 | transmembrane protein 145 [Aedes aegypti] | PTHR23252 | INTIMAL THICKNESS RECEPTOR-RELATED | |
| chr3 | 396274052 | 396299972 | Aedes_aegypti_formosus\|g9745.t1 | XP_021704186.1 | proteoglycan 4 isoform X2 [Aedes aegypti] | PTHR13526 | TRANSCRIPTION FACTOR SPT20 HOMOLOG | transcription coregulator activity#GO:0003712;transcription regulator activity#GO:0140110 |
| chr3 | 397076257 | 397079870 | Aedes_aegypti_formosus\|g9763.t1 | XP_021704327.1 | uncharacterized protein LOC5573210 isoform X1 [Aedes aegypti] | PTHR23354 | NUCLEOLAR PROTEIN 7/ESTROGEN RECEPTOR COACTIVATOR-RELATED | |
| chr3 | 396853206 | 396861041 | Aedes_aegypti_formosus\|g9760.t1 | XP_021704337.1 | zinc finger protein 423 homolog isoform X3 [Aedes aegypti] | PTHR24379 | KRAB AND ZINC FINGER DOMAIN-CONTAINING | |
| chr1 | 4753239 | 4755608 | Aedes_aegypti_formosus\|g10660.t1 | XP_001652118.2 | sodium channel protein Nach [Aedes aegypti] | PTHR11690 | AMILORIDE-SENSITIVE SODIUM CHANNEL-RELATED | |
| chr1 | 5123413 | 5128801 | Aedes_aegypti_formosus\|g10671.t1 | XP_001655854.1 | uncharacterized protein LOC5575827 [Aedes aegypti] | PTHR12887 | NANOS PROTEIN | mRNA binding#GO:0003729;organic cyclic compound binding#GO:0097159 |
| chr1 | 4968040 | 4991064 | Aedes_aegypti_formosus\|g10666.t1 | XP_021694452.1 | homeobox protein Hox-A1-like [Aedes aegypti] | PTHR24328 | HOMEOBOX PROTEIN MOX | RNA polymerase II cis-regulatory region sequence-specific DNA binding#GO:0000978;transcription regulator activity#GO:0140110;DNA-binding transcription factor activity, RNA polymerase II-specific#GO:0000981;organic cyclic compound binding#GO:0097159 |
| chr1 | 4837137 | 4879801 | Aedes_aegypti_formosus\|g10661.t1 | XP_021694455.1 | kinesin-like protein KIF19 isoform X2 [Aedes aegypti] | PTHR47968 | CENTROMERE PROTEIN E | |
| chr1 | 4681735 | 4713156 | Aedes_aegypti_formosus\|g10659.t1 | XP_021694460.1 | pyrokinin-1 receptor [Aedes aegypti] | PTHR24243 | G-PROTEIN COUPLED RECEPTOR | |
| chr1 | 14194192 | 14195964 | Aedes_aegypti_formosus\|g10875.t1 | XP_001655180.1 | uncharacterized protein LOC5574528 [Aedes aegypti] | PTHR21028 | SI:CH211-156B7.4 | |
| chr1 | 939476 | 1010695 | Aedes_aegypti_formosus\|g10544.t1 | XP_021694665.1 | cyclic AMP response element-binding protein B isoform X1 [Aedes a | PTHR45879 | CYCLIC AMP RESPONSE ELEMENT-BINDING PROTEIN B | RNA polymerase II cis-regulatory region sequence-specific DNA binding#GO:0000978;transcription regulator activity#GO:0140110;DNA-binding transcription factor activity, RNA polymerase II-specific#GO:0000981;organic cyclic compound binding#GO:0097159 |
| chr1 | 271077796 | 271082749 | Aedes_aegypti_formosus\|g16276.t1 | XP_021693725.1 | probable multidrug resistance-associated protein lethal(2)03659 [ | PTHR10769 | 40S RIBOSOMAL PROTEIN S28 | structural constituent of ribosome#GO:0003735 |
| chr1 | 285838777 | 285850976 | Aedes_aegypti_formosus\|g16554.t1 | XP_001654378.1 | translation initiation factor eIF-2B subunit alpha [Aedes aegypti | PTHR45860 | TRANSLATION INITIATION FACTOR EIF-2B SUBUNIT ALPHA | |
| chr3 | 11878918 | 11880441 | Aedes_aegypti_formosus\|g1401.t1 | XP_001660602.1 | gustatory and odorant receptor 24 [Aedes aegypti] | PTHR21143 | INVERTEBRATE GUSTATORY RECEPTOR | |
| chr1 | 4966620 | 4967989 | Aedes_aegypti_formosus\|g10665.t1 | XP_001655849.1 | MKI67 FHA domain-interacting nucleolar phosphoprotein [Aedes aeg | PTHR46754 | MKI67 FHA DOMAIN-INTERACTING NUCLEOLAR PHOSPHOPROTEIN | heterocyclic compound binding#GO:1901363;RNA binding#GO:0003723;organic cyclic compound binding#GO:0097159 |
| chr1 | 285332909 | 285346461 | Aedes_aegypti_formosus\|g16545.t1 | XP_001648160.2 | uncharacterized protein LOC5563844 [Aedes aegypti] | PTHR21567 | CLASP | cytoskeletal protein binding#GO:0008092;protein binding#GO:0005515;microtubule binding#GO:0008017 |
| chr1 | 276538862 | 276556837 | Aedes_aegypti_formosus\|g16391.t1 | XP_001663478.2 | alkaline phosphatase, tissue-nonspecific isozyme [Aedes aegypti] | PTHR11596 | ALKALINE PHOSPHATASE | phosphatase activity#GO:0016791 |
| chr1 | 276396896 | 276397546 | Aedes_aegypti_formosus\|g16384.t1 | XP_021693660.1 | extensin [Aedes aegypti] | PTHR31927 | FI07246P-RELATED-RELATED | |
| chr1 | 271033891 | 271074590 | Aedes_aegypti_formosus\|g16275.t1 | XP_021693725.1 | probable multidrug resistance-associated protein lethal(2)03659 [ | PTHR24223 | ATP-BINDING CASSETTE SUB-FAMILY C | ATP hydrolysis activity#GO:0016887;nucleoside-triphosphatase activity#GO:0017111;ATP-dependent activity#GO:0140657;transporter activity#GO:0005215;catalytic activity#GO:0003824;active transmembrane transporter activity#GO:0022804 |
| chr3 | 394540688 | 394541713 | Aedes_aegypti_formosus\|g9695.t1 | XP_021711515.1 | vicilin-like seed storage protein At2g18540 [Aedes aegypti] | PTHR23247 | NY-REN-41 ANTIGEN L15 -RELATED | |
| chr1 | 8879127 | 8886499 | Aedes_aegypti_formosus\|g10758.t1 | XP_001651037.2 | exonuclease mut-7 homolog [Aedes aegypti] | PTHR47765 | 3'-5' EXONUCLEASE DOMAIN-CONTAINING PROTEIN | |
| chr1 | 49038093 | 49039473 | Aedes_aegypti_formosus\|g11709.t1 | XP_021702255.1 | thiamine transporter 1-like [Aedes aegypti] | PTHR10686 | FOLATE TRANSPORTER | |
| chr1 | 285860473 | 285905223 | Aedes_aegypti_formosus\|g16557.t1 | XP_021693527.1 | uncharacterized protein LOC5563999 [Aedes aegypti] | PTHR24379 | KRAB AND ZINC FINGER DOMAIN-CONTAINING | |
| chr1 | 269885210 | 269886605 | Aedes_aegypti_formosus\|g16246.t1 | XP_021693744.1 | uncharacterized protein LOC110674165 [Aedes aegypti] | PTHR16134 | F-BOX/TPR REPEAT PROTEIN POF3 | |
| chr3 | 104479096 | 104479981 | Aedes_aegypti_formosus\|g3531.t1 | XP_021709000.1 | exonuclease 3'-5' domain-containing protein 2-like [Aedes aegypti | PTHR13620 | 3-5 EXONUCLEASE | |
| chr1 | 179495 | 204899 | Aedes_aegypti_formosus\|g10521.t1 | XP_001658554.1 | calphotin [Aedes aegypti] | PTHR48148 | KERATINOCYTE PROLINE-RICH PROTEIN | |
| chr1 | 2603404 | 2619249 | Aedes_aegypti_formosus\|g10604.t1 | XP_001661567.1 | aminopeptidase N [Aedes aegypti] | PTHR11533 | PROTEASE M1 ZINC METALLOPROTEASE | cation binding#GO:0043169;metallopeptidase activity#GO:0008237;zinc ion binding#GO:0008270;peptide binding#GO:0042277;aminopeptidase activity#GO:0004177;exopeptidase activity#GO:0008238;peptidase activity#GO:0008233;catalytic activity, acting on a protein#GO:0140096 |
| chr1 | 191602341 | 191641985 | Aedes_aegypti_formosus\|g14524.t1 | XP_021711796.1 | multidrug resistance-associated protein 9 [Aedes aegypti] | PTHR24223 | ATP-BINDING CASSETTE SUB-FAMILY C | ATP hydrolysis activity#GO:0016887;nucleoside-triphosphatase activity#GO:0017111;ATP-dependent activity#GO:0140657;transporter activity#GO:0005215;catalytic activity#GO:0003824;active transmembrane transporter activity#GO:0022804 |
| chr1 | 275873667 | 275885592 | Aedes_aegypti_formosus\|g16376.t1 | XP_001663295.2 | zinc finger protein 14 [Aedes aegypti] | PTHR24379 | KRAB AND ZINC FINGER DOMAIN-CONTAINING | |
| chr1 | 310223899 | 310226461 | Aedes_aegypti_formosus\|g17315.t1 | XP_001654124.2 | zinc finger protein 678 [Aedes aegypti] | PTHR16515 | PR DOMAIN ZINC FINGER PROTEIN | |
| chr1 | 281787470 | 281787823 | Aedes_aegypti_formosus\|g16496.t1 | XP_001656475.2 | protamine [Aedes aegypti] | PTHR47658 | HIGH MOBILITY GROUP B PROTEIN 12-RELATED | |
| chr1 | 270352302 | 270354066 | Aedes_aegypti_formosus\|g16260.t1 | XP_001649987.2 | synaptic vesicle glycoprotein 2B [Aedes aegypti] | PTHR23511 | SYNAPTIC VESICLE GLYCOPROTEIN 2 | |
| chr3 | 250845296 | 250845851 | Aedes_aegypti_formosus\|g6546.t1 | XP_001655720.2 | general odorant-binding protein 56d [Aedes aegypti] | PTHR11857 | ODORANT BINDING PROTEIN-RELATED | |
| chr1 | 285856780 | 285858005 | Aedes_aegypti_formosus\|g16556.t1 | XP_001648322.2 | zinc finger protein 501 [Aedes aegypti] | PTHR24379 | KRAB AND ZINC FINGER DOMAIN-CONTAINING | |
| chr1 | 275916231 | 275923412 | Aedes_aegypti_formosus\|g16377.t1 | XP_021693662.1 | zinc finger protein 271 isoform X2 [Aedes aegypti] | PTHR24388 | ZINC FINGER PROTEIN | |
| chr1 | 270333611 | 270335699 | Aedes_aegypti_formosus\|g16259.t1 | XP_001662720.1 | putative transporter SVOPL [Aedes aegypti] | PTHR23511 | SYNAPTIC VESICLE GLYCOPROTEIN 2 | |
| chr1 | 11565346 | 11566236 | Aedes_aegypti_formosus\|g10817.t1 | XP_021694517.1 | potassium voltage-gated channel protein Shaker isoform X5 [Aedes | PTHR45093 | TRANSCRIPTION ACTIVATOR MSS11 | |
| chr1 | 280984452 | 280985214 | Aedes_aegypti_formosus\|g16482.t1 | XP_001650809.1 | larval cuticle protein A2B [Aedes aegypti] | PTHR12236 | STRUCTURAL CONTITUENT OF CUTICLE | |
| chr1 | 278505490 | 278511290 | Aedes_aegypti_formosus\|g16435.t1 | XP_021693609.1 | vacuolar protein sorting-associated protein 13B [Aedes aegypti] | PTHR12517 | VACUOLAR PROTEIN SORTING-ASSOCIATED PROTEIN 13B | |
| chr1 | 294456259 | 294458109 | Aedes_aegypti_formosus\|g16752.t1 | XP_021693900.1 | myosin-2 heavy chain-like isoform X2 [Aedes aegypti] | PTHR45615 | MYOSIN HEAVY CHAIN, NON-MUSCLE | |
| chr1 | 294456259 | 294458109 | Aedes_aegypti_formosus\|g16752.t1 | XP_021693900.1 | myosin-2 heavy chain-like isoform X2 [Aedes aegypti] | PTHR45615 | MYOSIN HEAVY CHAIN, NON-MUSCLE | |
| chr3 | 393986660 | 394003417 | Aedes_aegypti_formosus\|g9682.t1 | XP_001652408.2 | putative polypeptide N-acetylgalactosaminyltransferase 9 [Aedes a | PTHR11675 | N-ACETYLGALACTOSAMINYLTRANSFERASE | |
| chr3 | 395720745 | 395741002 | Aedes_aegypti_formosus\|g9738.t1 | XP_021704263.1 | sex peptide receptor [Aedes aegypti] | PTHR47023 | SEX PEPTIDE RECEPTOR | |
| chr3 | 396477235 | 396478323 | Aedes_aegypti_formosus\|g9752.t1 | XP_011493380.2 | ecdysone-inducible protein E75 isoform X2 [Aedes aegypti] | PTHR24082 | NUCLEAR HORMONE RECEPTOR | |
| chr1 | 3411971 | 3412756 | Aedes_aegypti_formosus\|g10624.t1 | XP_021694709.1 | ubiquinone biosynthesis monooxygenase COQ6, mitochondrial [Aedes | PTHR43876 | UBIQUINONE BIOSYNTHESIS MONOOXYGENASE COQ6, MITOCHONDRIAL | oxidoreductase activity#GO:0016491;catalytic activity#GO:0003824 |
| chr1 | 5711623 | 5725144 | Aedes_aegypti_formosus\|g10690.t1 | XP_021694614.1 | protein lap4 isoform X3 [Aedes aegypti] | PTHR23119 | DISCS LARGE | |
| chr1 | 272842240 | 272844513 | Aedes_aegypti_formosus\|g16319.t1 | XP_021700392.1 | cullin-1 [Aedes aegypti] | PTHR11932 | CULLIN | ubiquitin protein ligase binding#GO:0031625 |
| chr1 | 278526535 | 278547701 | Aedes_aegypti_formosus\|g16437.t1 | XP_021693609.1 | vacuolar protein sorting-associated protein 13B [Aedes aegypti] | PTHR12517 | VACUOLAR PROTEIN SORTING-ASSOCIATED PROTEIN 13B | |
| chr1 | 285433466 | 285441777 | Aedes_aegypti_formosus\|g16546.t1 | XP_021693539.1 | tyrosine-protein kinase Fer isoform X4 [Aedes aegypti] | PTHR15735 | FCH AND DOUBLE SH3 DOMAINS PROTEIN | |
