## Supplementary material for "From macro to micro: De novo genomes of Aedes mosquitoes enable comparative genomics among close and distant relatives": Table S8

Supplementary Table S8 Selected genes found in inversions. Subset of genes found in inverted regions of *Aedes aegypti formosus* (Aaf) assembly. We identified by creating a local BLAST database from the proteins in the AaegL5 referene genome.

| Sequence | Start | End | Inversion ID | Genes ID | Gene name | Gene type |
|---|---|---|---|---|---|---|
| chr_1 | 3408801 | 3411805 | inv_4 | XP_021694708.1 | transient receptor potential channel pyrexia | channel |
| chr_1 | 4753239 | 4755608 | inv_4 | XP_001652118.2 | sodium channel protein Nach | channel |
| chr_1 | 11565346 | 11566236 | inv_4 | XP_021694517.1 | potassium voltage-gated channel protein Shaker isoform X5 | channel |
| chr_1 | 12065979 | 12095174 | inv_4 | XP_021694516.1 | potassium voltage-gated channel protein Shaker isoform X4 | channel |
| chr_1 | 13648574 | 13651855 | inv_4 | XP_021694486.1 | cytochrome P450 307a1 | p450 |
| chr_1 | 275132614 | 275134639 | inv_5 | XP_001650792.2 | odorant receptor 4 | or |
| chr_1 | 277085843 | 277087753 | inv_5 | XP_021693655.1 | heat shock protein 70 A1-like | hsp |
| chr_1 | 277088834 | 277090750 | inv_5 | XP_021693654.1 | heat shock protein 70 A1-like | hsp |
| chr_1 | 277110420 | 277112336 | inv_5 | XP_021693654.1 | heat shock protein 70 A1-like | hsp |
| chr_1 | 277117429 | 277119345 | inv_5 | XP_021693649.1 | heat shock protein 70 A1 | hsp |
| chr_1 | 277133127 | 277135043 | inv_5 | XP_021693649.1 | heat shock protein 70 A1 | hsp |
| chr_1 | 277138098 | 277140014 | inv_5 | XP_021693649.1 | heat shock protein 70 A1 | hsp |
| chr_1 | 280860049 | 280860648 | inv_5 | XP_001660359.1 | cuticle protein | cut |
| chr_1 | 280894255 | 280895060 | inv_5 | XP_001657546.2 | larval cuticle protein A3A | cut |
| chr_1 | 280895793 | 280896569 | inv_5 | XP_001660358.1 | larval cuticle protein A3A | cut |
| chr_1 | 280900225 | 280901024 | inv_5 | XP_001660357.1 | larval cuticle protein A3A | cut |
| chr_1 | 280906224 | 280907034 | inv_5 | XP_001660356.1 | cuticle protein | cut |
| chr_1 | 280909721 | 280910628 | inv_5 | XP_001660355.2 | cuticle protein | cut |
| chr_1 | 280911603 | 280912401 | inv_5 | XP_001660354.2 | cuticle protein | cut |
| chr_1 | 280917897 | 280918582 | inv_5 | XP_021693581.1 | cuticle protein isoform X4 | cut |
| chr_1 | 280925339 | 280926398 | inv_5 | XP_021693570.1 | cuticle protein-like isoform X2 | cut |
| chr_1 | 280933315 | 280934000 | inv_5 | XP_021693573.1 | cuticle protein isoform X3 | cut |
| chr_1 | 280938751 | 280939435 | inv_5 | XP_021693573.1 | cuticle protein isoform X3 | cut |
| chr_1 | 280945839 | 280946531 | inv_5 | XP_001660349.1 | larval cuticle protein A2B | cut |
| chr_1 | 280955374 | 280956064 | inv_5 | XP_021693574.1 | cuticle protein | cut |
| chr_1 | 280984452 | 280985214 | inv_5 | XP_001650809.1 | larval cuticle protein A2B | cut |
| chr_1 | 286069505 | 286120721 | inv_5 | XP_021693525.1 | E3 ubiquitin-protein ligase RNF19B isoform X3 | ubi |
| chr_3 | 11839180 | 11841420 | inv_1 | XP_021705712.1 | hsp90 co-chaperone Cdc37 | hsp |
| chr_3 | 11878918 | 11880441 | inv_1 | XP_001660602.1 | gustatory and odorant receptor 24 | or |
| chr_3 | 12406200 | 12431747 | inv_1 | XP_021705203.1 | E3 ubiquitin-protein ligase RNF126 | ubi |
| chr_3 | 250780551 | 250781320 | inv_2 | XP_001655718.2 | general odorant-binding protein 56d | obp |
| chr_3 | 250830930 | 250831387 | inv_2 | XP_001655719.1 | general odorant-binding protein 56d | obp |
| chr_3 | 250845296 | 250845851 | inv_2 | XP_001655720.2 | general odorant-binding protein 56d | obp |
| chr_3 | 250855838 | 250856295 | inv_2 | XP_001655721.2 | general odorant-binding protein 56d | obp |
| chr_3 | 250861418 | 250861875 | inv_2 | XP_001655722.2 | general odorant-binding protein 56a | obp |
| chr_3 | 250877327 | 250877786 | inv_2 | XP_001655723.2 | general odorant-binding protein 56a | obp |
| chr_3 | 250891332 | 250891871 | inv_2 | XP_001655725.2 | general odorant-binding protein 56d | obp |
| chr_3 | 250903556 | 250904095 | inv_2 | XP_001655725.2 | general odorant-binding protein 56d | obp |
| chr_3 | 397021438 | 397043727 | inv_3 | XP_021704328.1 | heat shock factor protein isoform X1 | hsp |
